# UFMylation orchestrates chromatin engagement of core NHEJ components to promote DNA double-strand break repair

**DOI:** 10.1101/2025.06.16.659844

**Authors:** Zijuan Wang, Benjamin M. Foster, Isabelle C. da Costa, Yue Wu, Deepak Behera, Francesca Conte, Eleanor W. Trotter, Maria Jose Cabello-Lobato, Shweta Choudhary, Reuven Wiener, Petra Beli, Duncan L. Smith, William H. Banks, Steven Bagley, Shane McKee, Meenakshi Minnis, Stefan Meyer, Amanda K. Chaplin, Wolfgang Dörner, Henning D. Mootz, Iain M. Hagan, Yaron Galanty, Igor Larrosa, Matt J. Cliff, Christine K. Schmidt

**Affiliations:** Manchester Cancer Research Centre, Division of Cancer Sciences, School of Medical Sciences, Faculty of Biology, Medicine and Health, University of Manchester, Manchester, UK; Department of Chemistry, School of Natural Sciences, University of Manchester, Manchester, UK; Institute of Molecular Biology (IMB), Mainz, Germany; Cell Division Group, CRUK Manchester Institute, The University of Manchester, Manchester, UK; Department of Biochemistry and Molecular Biology, The Institute for Medical Research, Israel-Canada Hebrew University-Hadassah Medical School, Jerusalem, Israel; Institute of Developmental Biology and Neurobiology (IDN), Johannes Gutenberg-Universität, Mainz, Germany; Mass spectrometry, CRUK Manchester Institute, The University of Manchester, Manchester, UK; Microscopy, CRUK Manchester Institute, The University of Manchester, Manchester, UK; Department of Genetic Medicine, Belfast City Hospital, Belfast, UK; Manchester Academic Health Science Centre, Manchester, UK; Department of Genetic Medicine, St Mary’s Hospital, Central Manchester NHS Foundation Trust, Manchester, UK; Department of Paediatric Haematology and Oncology, Royal Manchester Children’s Hospital, Manchester, UK; Young Oncology Unit, The Christie NHS Foundation Trust, Manchester, UK; Leicester Institute for Structural and Chemical Biology, Department of Molecular and Cell Biology, University of Leicester, Leicester, UK; Institute of Biochemistry, University of Münster, Münster, Germany; The Gurdon Institute and Department of Biochemistry, University of Cambridge, Cambridge, UK; CRUK Cambridge Institute, University of Cambridge, Cambridge, UK; Manchester Institute of Biotechnology (MIB), University of Manchester, Manchester, UK

**Keywords:** UFM1, ubiquitin-like protein (UBL), DNA repair, non-homologous end-joining (NHEJ), genetic code expansion, photo-crosslinking

## Abstract

DNA double-strand breaks (DSBs) are highly cytotoxic lesions whose misrepair can lead to genomic instability, cancer and developmental disorders. Through systematic screening of understudied ubiquitin-like modifiers (UBLs), we identify UFM1 as a previously unrecognised regulator of non-homologous end-joining (NHEJ). Using a structure-guided chemical biology strategy, we develop a photo-crosslinkable UFM1 probe and, together with high-resolution NMR, uncover non-canonical UFM1-binding regions in core NHEJ components, including XRCC4. Mechanistically, proximity-dependent proteomics reveals Ku70 as a key UFMylation substrate, establishing a functional axis in which XRCC4 engages UFMylated Ku70 to promote the chromatin assembly of NHEJ factors. Perturbation of UFM1 signalling, via UFSP2 depletion or a hypomorphic UBA5 allele in patient-derived fibroblasts, impairs these processes, linking UFMylation defects to altered regulation of DSB repair. Our findings define a complete UFM1 signalling module in genome maintenance and uncover a molecular connection between hereditary UFMylation disorders and dysregulated DSB repair pathways.

## INTRODUCTION

Maintenance of genome stability is key to cellular survival and human health, yet our DNA is under constant assault from both endogenous and exogenous sources. To deal with these attacks, cells have evolved a complex and highly coordinated network of pathways known as the DNA damage response (DDR). The DDR detects, signals, and repairs DNA lesions while modulating cell cycle progression to allow time for repair or, when necessary, triggering apoptosis. Deficiencies in these repair mechanisms are linked to a wide array of human diseases, including cancer, immunodeficiency syndromes, neurodegenerative conditions, and premature ageing^1–3^.

DNA double-strand breaks (DSBs) are amongst the most cytotoxic lesions and are primarily repaired by two major pathways: homologous recombination (HR) and non-homologous end joining (NHEJ). HR is a highly accurate repair process that uses the sister chromatid as a template, restricting it to the S and G2 phases of the cell cycle. By contrast, NHEJ can function throughout interphase but can be mutagenic. Despite that, it is the predominant DSB repair pathway in mammalian cells^4,5^. NHEJ is initiated by the binding of the Ku70:Ku80 heterodimer to DNA ends, followed by recruitment of the core NHEJ kinase DNA-PKcs to form the DNA-PK holoenzyme. This complex then facilitates chromatin recruitment of a series of NHEJ factors, including XRCC4, XLF (also known as NHEJ1), PAXX, and DNA ligase IV (LIG4)^6^.

The functions of many DDR proteins, including those involved in DSB repair, are tightly regulated by posttranslational modifications (PTMs), particularly ubiquitin and ubiquitin-like proteins (UBLs). While ubiquitin, SUMO, and NEDD8 have well-established roles in DNA damage responses^7–10^, most other UBLs encoded in the human genome remain understudied. This knowledge gap limits our broader understanding of how UBL-mediated signalling networks regulate genome stability and cellular stress responses^11–13^. UBLs, like ubiquitin, are typically conjugated to substrate proteins via a conserved enzymatic cascade involving an E1 activating, an E2 conjugating, and an E3 ligating enzyme, with the modifications influencing the functionality of their targets in different ways, often mediated through selective binding by downstream reader or receptor proteins. As such, UBL receptors play a central role in translating upstream UBLylation events into specific biochemical outcomes. To fully delineate the regulatory potential of these modifications in the DDR, it is essential to define both the upstream UBL-conjugated substrates and their corresponding receptor proteins. However, these interactions are often transient and weak, presenting significant challenges for conventional biochemical identification methods. This bottleneck has limited their discovery and characterisation to often less than a handful known UBL substrate-receptor axes.

Here, we employ a two-pronged functional screening strategy to systematically interrogate the contribution of 10 understudied UBLs to DSB repair. This approach reveals diverse and pathway-specific roles for multiple UBLs across different branches of the DDR. Amongst these, UFM1 emerged as the most prominent hit across several independent DDR readouts. As the most recently identified UBL, UFM1 is garnering increasing attention for its roles beyond its canonical functions at the endoplasmic reticulum^14^. To address the challenge of defining UFM1 substrate-receptor relationships, we develop a fully functional, photo-crosslinkable UFM1 probe for receptor screening. The approach identifies a broad spectrum of UFM1-interacting proteins, spanning biological processes known to be regulated by UFM1 as well as unknown associations. Notably, several core NHEJ components ranked prominently amongst the candidates. Through site-directed mutagenesis, crosslinking mass spectrometry (XL-MS), biochemical assays, and high-resolution nuclear magnetic resonance (NMR), we validate and characterise non-canonical UFM1-binding regions on several NHEJ proteins, including XRCC4. To identify upstream UFMylated targets of these receptors, we performed proximity labelling of the UFM1 E2 enzyme UFC1 in cells, pinpointing the DNA-PK complex as a UFMylation substrate, amongst a wide range of candidate targets, with implications in DDR settings as well as broader cellular contexts. Mechanistically, we demonstrate that UFMylation is critical for facilitating the recruitment of key NHEJ components to damaged chromatin and that – within this context – XRCC4 functions as a receptor for Ku70 UFMylated at K526, defining a novel functional UFM1-Ku-XRCC4 axis in genome stability maintenance. Beyond providing structural, mechanistic and functional insights into DSB repair pathways, our findings establish a link between impaired NHEJ and UFMylation deficiency in patients, highlighting shared developmental phenotypes such as microcephaly, observed in UFMylation disorders as well as classical NHEJ syndromes^14,15^.

## RESULTS

### Systematic UBL screening reveals UFM1 functions in non-homologous end-joining (NHEJ)

To systematically assess the roles of understudied UBLs in DSB repair, we performed loss-of-function screening in U2OS cells using siRNA pools targeting 10 UBLs with limited involvement in the DDR. These included ATG12, FAT10, FUBI, ISG15, UBL3, UBL4A, UBL4B, UBL5, UFM1, and URM1. The screen was carried out using a two-pronged approach. Module 1 employed the traffic-light reporter (TLR) system to quantify HR and mutagenic end-joining (mutEJ) efficiency^16,17^(Fig. 1A), while Module 2 measured the formation of ionising radiation-induced foci (IRIF) of γH2AX, 53BP1 whose recruitment requires ubiquitin, and conjugated ubiquitin (Fig. 1B). Depletion of several UBLs, including FUBI, UBL5, ATG12, UFM1, and ISG15 resulted in reduced efficiency of both HR and mutEJ (Fig. S1A). Others, such as UBL4B, UBL3, URM1, and UBL4A, predominantly impaired HR, suggesting more pathway-specific contributions to DSB repair. Follow-up analyses indicated that the effects observed for FUBI were likely attributable to off-target activity, potentially due to sequence overlap between one of the siRNAs and UBE2I (also known as UBC9), the E2 conjugating enzyme for SUMO. Given SUMO’s well-established roles in DSB repair^18–21^, FUBI was therefore excluded from further analysis. Strikingly, depletion of several UBLs with distinct DSB repair efficiency profiles, namely UFM1, ISG15, URM1, and UBL4A, also disrupted ubiquitin conjugation and/or 53BP1 IRIF formation at DNA damage sites (Fig. S1B). Amongst these UBLs, UBL5, which scored highest in the TLR screen for promoting HR, is known to support the Fanconi anaemia pathway^22^ and maintains expression of the HR factor XRCC3^23^, providing orthogonal validation of our screening approach. UFM1 and ISG15, both of which compromised HR, mutEJ and IRIF formation upon depletion, have previously been implicated in DNA replication stress and DDR functions^13,24–33^. Furthermore, autophagy, for which ATG12 is crucial, can promote the DNA damage response in different genotoxic contexts^34–36^, and UBL4A has been reported to facilitate BRCA1 recruitment to DNA damage sites^37^. However, these previously reported functions cannot explain the complex phenotypes we observed for multiple UBLs in different DSB repair processes. Together, the findings of our screen reveal a functionally diverse involvement of UBLs in DSB repair, uncovering both established and previously unrecognised roles in genome stability maintenance and opening new avenues for mechanistic investigation.

**Figure 1.**
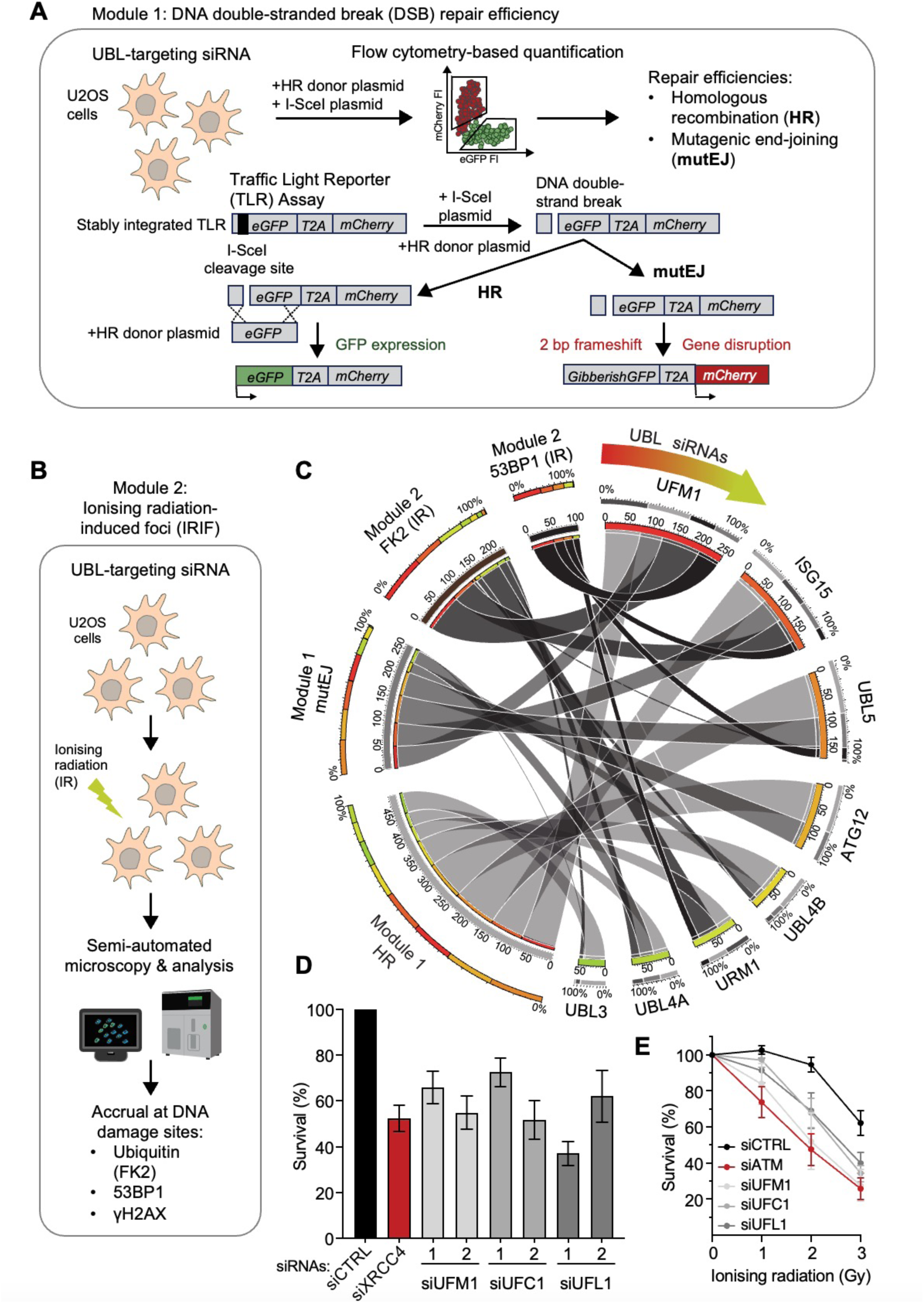
Two-module screening of UBLs reveals UFM1 as a key regulator in the DNA damage response. (**A**) Schematic of the screening pipeline for Module 1 using the traffic light reporter (TLR) system, to evaluate efficiencies of DSB repair by homologous recombination (HR) or mutagenic end-joining (mutEJ)^16,17^. (**B**) Experimental screening workflow for Module-2, encompassing high-throughput/high-content quantitative imaging and automated IRIF detection of ionising radiation-induced foci (IRIF), in response to 2 Gy monitored at different time points, of γH2AX – a DNA damage marker – 53BP1 and conjugated ubiquitin detected by FK2 antibody. (**C**) Circos plot^91^, illustrating the comparative effects of siUBL depletion on the DDR read-outs screened. Individually plotted siUBL effects, and further screening information, is available in the Supplemental Figures for Module 1 (Fig. S1A) and Module 2 (Fig. S1B). (**D**) Random plasmid integration results for two independent siRNAs targeting key components of the UFM1 pathway: UFM1, UFC1 or UFL1, compared to non-targeting control siRNA (siCTRL) and siXRCC4, targeting a core non-homologous end-joining (NHEJ) factor as a positive control. Data represent means ± SEM for n ≥ 3 independent experiments. Additional assay information is available in Figure S3. (**E**) Clonogenic survival assays of U2OS cells after IR treatment, or not, for siRNAs targeting key components of the UFM1 pathway: UFM1, UFC1 or UFL1, as indicated, compared to a non-targeting siRNA (siCTRL) and siATM as negative and positive controls, respectively. Data represent means ± SEM for n ≥ 3 independent experiments. Abbreviations: HR: homologous recombination; IR: ionising radiation; mutEJ: mutagenic end-joining; SEM: standard error of the mean; TLR: traffic light reporter (TLR).

Amongst all UBLs, UFM1 scored as the strongest candidate across multiple screening readouts (Fig. 1C), indicating an important role in diverse DDR aspects. Like ubiquitin and other well-characterised UBLs, UFM1 is conjugated to substrates via an enzymatic cascade, consisting of a single member of an activating E1 enzyme (UBA5), a conjugating E2 enzyme (UFC1) and an E3 ligating complex (UFL1 in association with DDRGK1, also known as UFBP1)^14,38–41^. Previously reported roles in ATM signalling through UFMylation of MRE11 and histone H4,^25–27^ are consistent with the HR defects we observed in our TLR screen (Fig. S1A). However, these known roles do not explain the additional impacts we observed on end-joining repair and IRIF kinetics, the latter of which we validated using two independent UFM1 siRNAs (Fig. S2A-C). To further assess the importance of UFM1 in end-joining repair, we employed a complementary assay based on random plasmid integration^19^. Using two independent siRNAs targeting UFM1, the E2 (UFC1), or the E3 (UFL1) enzyme, significantly impaired end-joining efficiency to levels comparable with depletion of XRCC4, a core NHEJ factor (Fig. 1D, Fig. S3A-B). Consistent with this, loss of UFM1 pathway components hypersensitised cells to IR (Fig. 1E). Collectively, these results highlight UFM1’s functional importance across a range of DDR readouts, including end-joining repair.

### A photo-crosslinkable probe for UFM1 receptor discovery

Given the important role of downstream UBL receptors in determining the outcomes of UBLylation events, we employed a bottom-up strategy to systematically identify UFM1 receptors. Our goal was to illuminate the mechanisms underpinning UFM1-dependent end-joining repair and to use receptor discovery as a springboard for defining upstream UFMylation axes. Recognising that UBL:receptor interactions are often transient and weak, we developed a photo-crosslinkable, non-conjugatable UFM1 probe lacking its C-terminal exposed glycine to stabilise these interactions through light-inducible covalent bonds. To facilitate stringent purification, the probe was equipped with a biotin handle conjugated to a cysteine added to UFM1’s C-terminus, enabling affinity-based enrichment and subsequent identification of covalently bound receptors via liquid chromatography tandem mass spectrometry (LC-MS/MS; Fig. 2A). In designing the probe, we leveraged structural insights into UFM1’s interaction with its E1 enzyme UBA5^42^, which is mediated through an α-β groove on UFM1, a known receptor interaction hotspot for paralogues of a related UBL, SUMO^43^. We reasoned that, analogous to SUMO, this surface may also mediate broader receptor interactions for UFM1.

**Figure 2.**
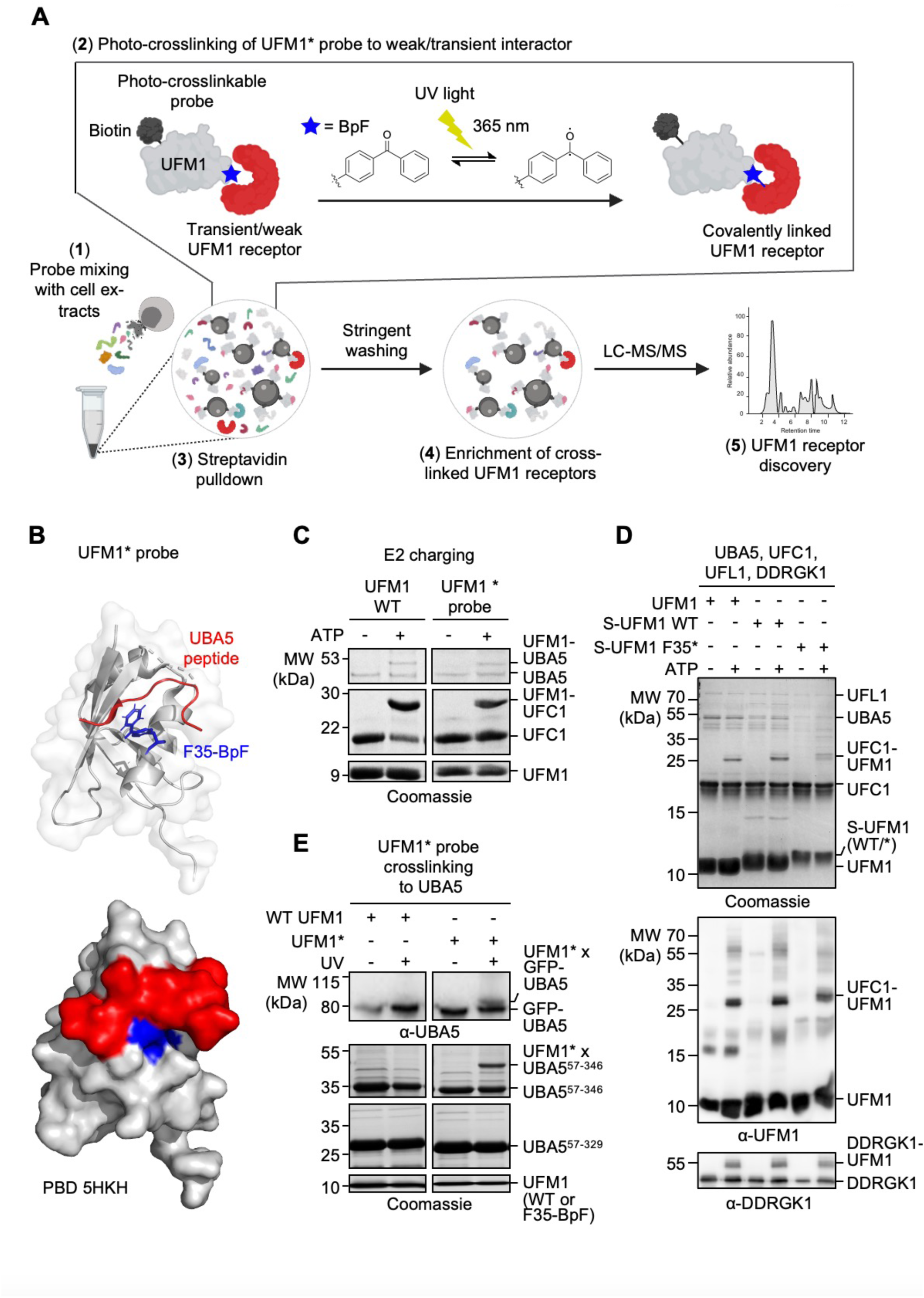
A photo-crosslinkable UFM1 probe for non-covalent interactor screening. (**A**) Schematic illustrating the workflow of inducing covalent bonds with weak and transient interactors, provided via cellular extracts, targeting a selected UFM1 surface-exposed region-of-interest, marked with a photo-crosslinkable unnatural amino acid, 4-benzoyl-(L)-phenylalanine (BpF, blue star), using UV light at a wavelength of 365 nm. Crosslinked interactors are precipitated via a biotin handle integrated into the UFM1 probe and identified by liquid chromatography tandem mass spectrometry (LC-MS/MS). (**B**) Ribbon (top) and surface (bottom) structure of UFM1 (PDB 5HKH) with BpF integrated at residue F35 (highlighted in blue), illustrating BpF’s proximity to a bound peptide of UBA5 (highlighted in red), a known non-covalent interactor targeting UFM1’s α-β groove of interest. Other UFM1 residues are shown in grey. (**C**) E2 charging assays, illustrating the functional integrity of the UFM1 F35-BpF probe for its loading to UFC1. (**D**) *In vitro* UFMylation assays, demonstrating the functional integrity of the UFM1 F35-BpF probe for its covalent attachment to acceptor sites on UFC1 and DDRGK1. (**E**) Photo-crosslinking reactions (365 nm, 2 h) of UFM1 (wildtype (WT) or F35-BpF (*)) with whole cell extracts obtained from HEK293T cells ectopically expressing GFP-UBA5 (top panel), or with recombinant UBA5, containing (UBA5^57-346^) or lacking (UBA5^57-329^) a known UFM1-interacting sequence (UIS; 2^nd^ and 3^rd^ panel). Loading controls for UFM1, WT or F35-BpF, are shown at the bottom. Abbreviations: α: anti; BpF: 4-benzoyl-(L)-phenylalanine; LC-MS/MS: liquid chromatography tandem mass spectrometry; MW: molecular weight; S-UFM1: Strep-UFM1; UFM1*: UFM1 F35-BpF probe; UIS: UFM1-interacting sequence; UV: ultraviolet; WT: wildtype.

We engineered three UFM1 probes by incorporating the photo-crosslinkable, unnatural amino acid 4-benzoyl-(L)-phenylalanine (BpF)^44^ at residues K34, F35, and E39 – each located within crosslinking distance to UBA5 and predicted to not disrupt binding based on the reported UFM1:UBA5 interaction^42^ (Fig. 2B, Fig. S4A). Amongst these, a conjugatable version of the F35 probe best preserved wild-type activity in E2 charging assays *in vitro* (Fig. 2C, Fig. S4B), possibly due to BpF more closely representing aspects of phenylalanine’s physicochemical features than those of lysine or glutamate. Further characterisation in *in vitro* UFMylation assays of a conjugatable version of UFM1 F35-BpF confirmed that the probe retained full functional activity also as a posttranslational modifier (Fig. 2D). Moreover, the UFM1 F35-BpF probe efficiently crosslinked to UBA5, both when transiently expressed in cellular extracts (Fig. 2E, top panel) or when purified as a recombinant protein (Fig. 2E, UBA5^57–346^, middle and bottom panel). The latter depended on UBA5 containing its reported UFM1-interacting sequence (UIS, Fig. 2E, UBA5^57–3^, middle panel), with no crosslinking detected in a truncated version of the protein lacking the UIS^42^ (Fig. 2E, UBA5^57–329^, bottom panel). Furthermore, using crosslinking mass spectrometry (XL-MS), the highest-confidence UBA5 peptide crosslinked to UFM1 F35-BpF via a residue within the UIS, highlighting the regioselectivity and specificity of the crosslinkable UFM1 probe. A second peptide signal within the adenylation domain may indicate the presence of additional, lower-affinity UFM1 interaction sites on UBA5, potentially contingent on UFM1 engagement with the UIS (Fig. S4C). Taken together, these findings validate UFM1 F35-BpF as a robust and highly selective tool for capturing downstream UFM1 receptors via its α-β groove.

### Photo-crosslink screening identifies UFM1 receptor candidates across varied ontologies

To identify UFM1 receptors relevant to NHEJ, we applied the photo-inducible UFM1 F35-BpF probe – hereafter referred to also as UFM1* probe – to cellular extracts from IR-treated HEK293T cells, ensuring capture under relevant DNA damage conditions. Induced crosslinking resulted in a broad smear of high-molecular-weight species, which was absent in the non-photo-induced control, confirming that the UFM1* probe successfully crosslinked to a range of cellular proteins (Fig. 3A, left panel). Affinity purification followed by LC-MS/MS analysis identified >600 high-confidence candidate receptors enriched specifically in the crosslinked condition, with minimal background in non-crosslinked controls, highlighting the effectiveness of using a stringent washing protocol (Fig. 3A, Supplementary Table S1)^43,45–47^. As expected, UFM1 itself, UBA5, and DDRGK1 were amongst the top hits, validating the approach. UFL1, the known E3 ligase, was also identified in one replicate (Fig. 3A), suggesting that additional components of the UFMylation cascade may engage in direct UFM1 recognition, consistent with observations from other UBL systems (e.g. most SUMO E3 ligases harbour SUMO-interacting motifs that bind the α-β groove)^48^.

**Figure 3.**
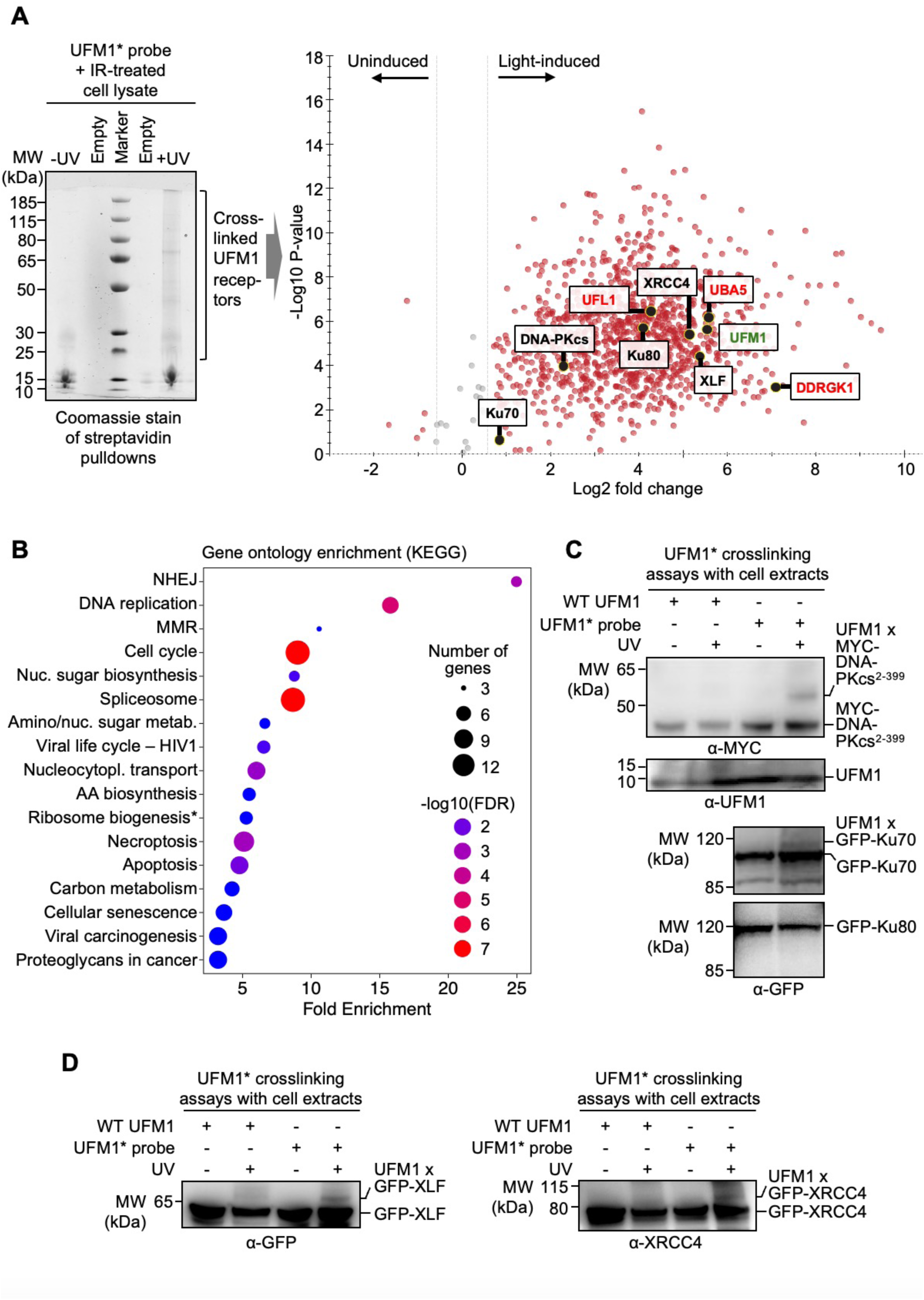
Photo-crosslink screening identifies UFM1 interactors across diverse ontologies including NHEJ. (**A**) Left: Streptavidin pulldowns, showing a broad smear of high-molecular-weight species, indicative of a diverse range of interactors crosslinked to the biotinylated UFM1 F35-BpF probe (UFM1*) detectable upon treatment with UV light (365 nm, 2 h, right lane), but not in the non-UV-treated control (left lane). Cell lysates were obtained from ionising radiation (IR)-treated HEK293T cells (30 min post 10 Gy). Right: Volcano blot, showing UFM1 interactor candidates detected in one of the three biological replicates, as a representative (rep 1, Supplementary Table S1). UFM1 (the bait) is highlighted in green, components of the UFM1 system, including known interactors in red, and select candidates-of-interest as part of the NHEJ complex in blue. (**B**) Gene ontology enrichments (KEGG) with a false discovery rate (FDR) < 0.05 for UFM1 interactor candidates detected in at least two out of the three biological replicates performed for the screen. Asterisk denotes ribosome biogenesis in eukaryotes. (**C**) Photo-crosslinking of a MYC-tagged N-terminal truncation of DNA-PKcs (residues 2-399) to UFM1 F35-BpF (UFM1*), using wildtype (WT) UFM1 as a control (top). The two bottom panels show photo-induced crosslinking experiments of the UFM1 F35-BpF probe with Ku, highlighting a crosslinked band with GFP-Ku70, but not GFP-Ku80. Photo-crosslinking reactions (365 nm, 2 h) were with whole cell extracts obtained from HEK293T cells ectopically expressing the tagged proteins, as indicated. (**D**) as (C) but for GFP-tagged XLF (left) and GFP-tagged XRCC4 (right). Abbreviations: α: anti; AA: amino acid; FDR: false discovery rate; IR: ionising radiation; Metab.: metabolism; MMR: mismatch repair MW: molecular weight; NHEJ: non-homologous end-joining; Nuc: nucleotide; Nucleocytopl.: nucleocytoplasmic; UFM1*: UFM1 F35-BpF; UV: ultraviolet; WT: wildtype.

Gene ontology (GO) analysis of the identified candidates revealed enrichment across diverse processes (Fig. 3B), including ontologies previously linked to UFM1 like apoptosis^49–53^, cell cycle^54–56^, senescence^57^, and viral life cycle regulation^58^, supporting the functional relevance of our hits. Notably, DNA replication-associated pathways also featured prominently, aligning with recent findings implicating UFMylation in the regulation of replication stress response factors such as PARP1 and PTIP^28–30^. Interestingly, ontologies related to carbon metabolism and amino acid biosynthesis – top hits in a recent UFMylation target screen^59^ – were also enriched, suggesting potential functions for UFM1 receptor interactions in these newly reported areas, as well as other pathways with un-or less established associations to UFM1, such as DNA mismatch repair, the spliceosome, and nucleocytoplasmic transport (Fig. 3B). Strikingly, the top-enriched pathway was NHEJ (Fig. 3B), with all subunits of the DNA-PK complex (the catalytic subunit DNA-PKcs, Ku70, Ku80) and the core factors XRCC4 and XLF amongst the hits (Fig. 3A). This is in strong agreement with our earlier siUBL functional screen identifying UFM1 as an important regulator of end-joining repair (Fig. 1C-E). Overall, these findings establish our UFM1* probe as a robust tool for receptor discovery and provide a comprehensive resource for exploring UFMylation roles across different cellular pathways, including a mechanistically grounded link to NHEJ.

### Core NHEJ components feature multiple, diverse UFM1-interacting modules

To dissect the molecular basis of UFM1 interactions with core NHEJ factors, we first tested whether the UFM1* probe could directly crosslink to components of the NHEJ complex when expressed in cellular extracts. We observed photo-induced crosslinking between UFM1* and multiple subunits of the DNA-PK complex, including the N-terminal 400 amino acids of DNA-PKcs, Ku70 (Fig. 3C), as well as the core NHEJ factors XLF and XRCC4 (Fig. 3D). No direct crosslinking could be detected for Ku80, despite its presence in the mass spectrometry screen. Given the high-affinity interaction between Ku70 and Ku80, which tend to remain tightly associated during affinity purification^60^, it is likely that Ku80’s co-detection reflects indirect association through Ku70 rather than direct UFM1 binding.

Focusing on XRCC4 and XLF – two structurally related scaffold proteins that are critical for the assembly of synaptic complexes^61,62^ to promote the ligation of the broken DNA – we further confirmed direct interactions of UFM1* with XRCC4 *in vitro* using recombinant proteins coupled with photo-crosslinking (Fig. S5A). To gain molecular insights into these interactions, we first performed XL-MS to identify XRCC4 peptides covalently linked to UFM1*. The analysis revealed five crosslinked peptides spanning three discrete regions of the XRCC4 dimer: (1) the globular head domain, (2) the coiled-coil stalk, and (3) its C-terminal intrinsically disordered region (IDR) (Fig. S5B).

To further explore region 1, we leveraged NMR titration experiments using a previously assigned structure of the XRCC4 head domain^63^. These studies identified significant UFM1-induced chemical shift perturbations (CSPs) and signal broadening in residues surrounding V47 – consistent with the XL-MS crosslink – highlighting a coherent surface patch encompassing residues A56, M59, V104, and S105 as likely residues mediating UFM1 interaction (Fig. 4A-B). This surface overlaps with known binding interfaces for SUMO^64^ and XLF^65^, supporting its role as a regulatory protein:protein interaction hotspot. Additional shifts in more isolated or buried residues likely reflect allosteric effects of binding rather than direct contact (Fig. 4A, Fig. S6A-B). To assess the contribution of the UIS-like motif in region 2, which has remained undetectable in previous NMR studies, we performed GST-UFM1 pulldown assays using an XRCC4 mutant in which the five key residues (FILVL) were substituted with alanines (XRCC4^180–5A^). Binding to UFM1 was markedly reduced in the mutant over wild-type XRCC4 (Fig. 4C), confirming the role of the XRCC4 180 region in UFM1 engagement. For region 3 located in the XRCC4 IDR, NMR experiments showed UFM1-induced CSPs overlapping with the crosslinked peptide (Fig. 4D, Fig. S7), and GST-UFM1 pulldown assays further identified residues 256-263 as required for proficient binding (Fig. 4E).

**Figure 4.**
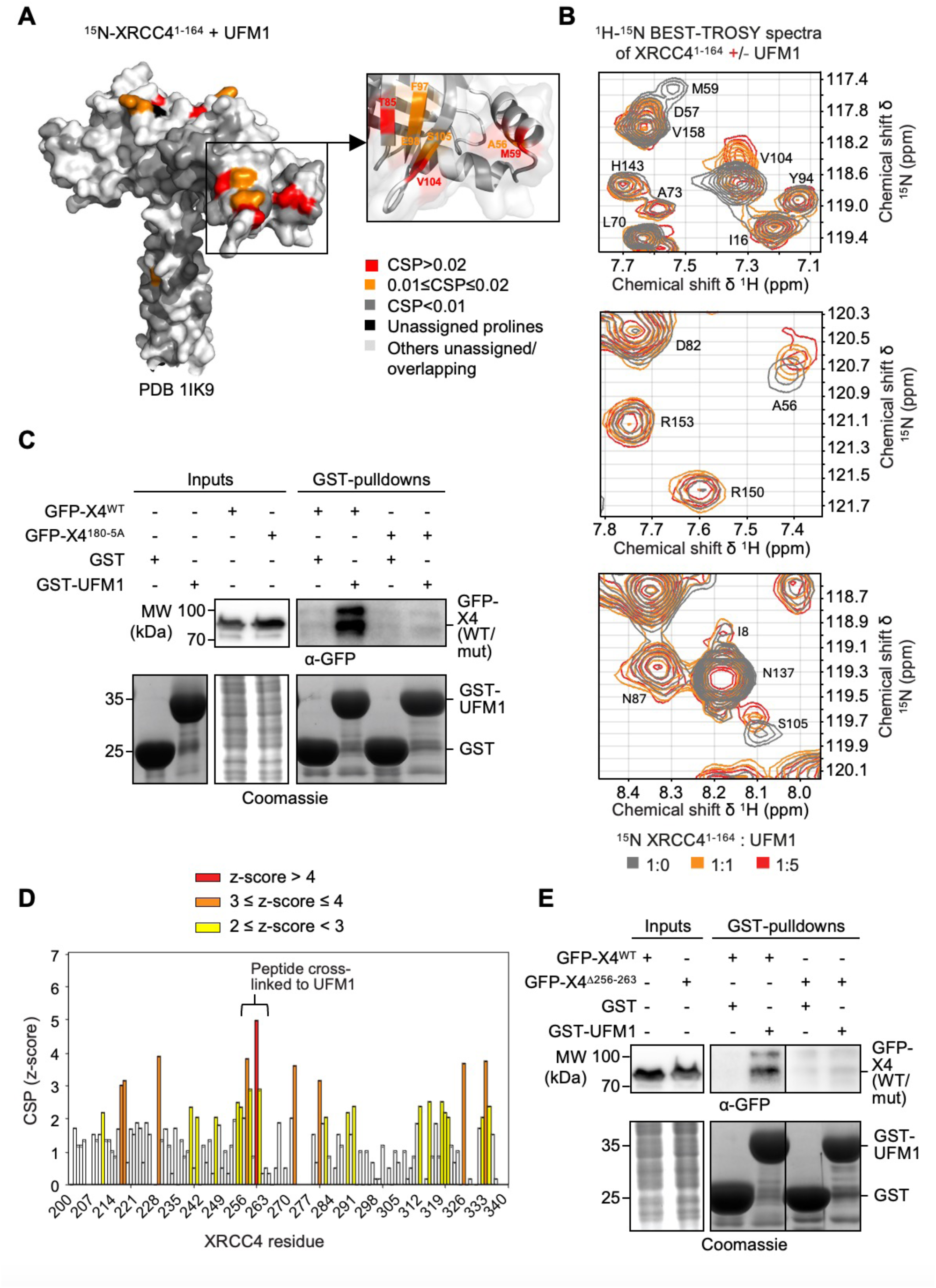
XRCC4 features multiple UFM1-binding modules. (**A**) Chemical shift perturbations (CSPs) in the ^1^H-^15^N BEST-TROSY spectra of XRCC4^1–164^ (PDB 1IK9) after addition of increasing concentrations of UFM1. Colour code ranges from red (residues most affected by binding) to dark grey (residues unaffected by binding), as indicated. Unassigned prolines are displayed in black, other unassigned/overlapping residues in light grey. Detailed view of key XRCC4^1-164^ residues affected on the sides of the XRCC4 head domain is shown on the right. (**B**) Representative panels of key interacting residues of the XRCC4^1-164 1^H-^15^N BEST-TROSY spectra (grey) overlayed with the XRCC4^1-164^ spectra after the addition of 1 (yellow) or 5 (red) molar equivalents of UFM1. (**C**) UFM1-GST-pulldowns with whole cell extracts of HEK293T cells, ectopically expressing GFP-tagged wildtype (WT) XRCC4 (X4) or XRCC4^180-5A^. (**D**) CSP profile (z-scores) in the ^1^H-^15^N HSQC spectra of XRCC4^200-334^ after addition of 1 molar equivalent of UFM1. Colour code ranges from red (residues most affected by binding) over orange to yellow (residues least affected by binding), as indicated. (**E**) UFM1-GST-pulldowns with whole cell extracts of HEK293T cells, ectopically expressing GFP-tagged wildtype (WT) XRCC4 (X4) or _XRCC4Δ256-263._ Abbreviations: CSP: chemical shift perturbation; MW: molecular weight; SD: standard deviation; WT: wildtype; X4: XRCC4.

In contrast to XRCC4, XL-MS analysis of recombinant XLF, crosslinked to UFM1 F35-BpF (Fig. S8A), did not yield high-confidence crosslinked peptides. Instead, we adopted a domain-mapping approach using a panel of XLF mutants, and identified residues 199-207, corresponding to a surface-exposed loop C-terminal to the stalk domain, as an important UFM1-binding region (Fig. S8B-C). Mutation of this region reduced UFM1 binding in GST-UFM1 pulldown assays (Fig. S8D), confirming its importance for the interaction.

Reciprocally, NMR titrations showed that both XRCC4 and XLF induced CSPs in similar UFM1 regions, overlapping with key changes observed upon binding of the UIS of UBA5 (Fig. S9A-C, Fig. S10A-B, Fig. S11A-B Fig. S12A-B)^42^. These perturbations predominantly localised to the α-β groove of UFM1, supporting a shared general binding interface. While the general interacting region was conserved, the observed perturbation patterns revealed nuanced differences in the mode of interaction across receptors with distinct sequence features. These findings align with the original design of the UFM1* probe to target this surface. In summary, these findings reveal that UFM1 engages multiple direct binding interfaces across NHEJ core factors, highlighting an unexpected level of interaction plasticity. We identify at least four bona fide UFM1 receptors – XRCC4, XLF, DNA-PKcs (N-terminal), and Ku70 – that engage UFM1 through distinct recognition modules. These results significantly expand our currently limited repertoire of around a handful of in-depth validated UFM1 receptors, and provide structural and molecular insights into UFM1 recognition modules.

### Proximity-labelling implicates the DNA-PK complex as a UFMylation target

To uncover how NHEJ components might act as functional UFM1 receptors in end-joining repair, we next sought to identify putative substrates of UFMylation in NHEJ-relevant contexts. To this end, we employed an inducible proximity biotinylation strategy in cells using engineered peroxidase APEX2^66^ fused to UFC1, the E2 conjugating enzyme for UFM1, together with a nuclear localisation signal (NLS; Fig. 5A). The NLS was used to maintain the natural localisation of UFC1 as both a cytoplasmic and nuclear protein^67^. Given that E2s pair up with E3s to engage with their substrates to facilitate UFM1 transfer^68^, we reasoned this setup would allow for spatially restricted biotinylation of potential UFM1 substrates residing in proximity to UFC1. Biotinylated proteins were then retrieved via neutravidin pulldown and identified by mass spectrometry^66^.

**Figure 5.**
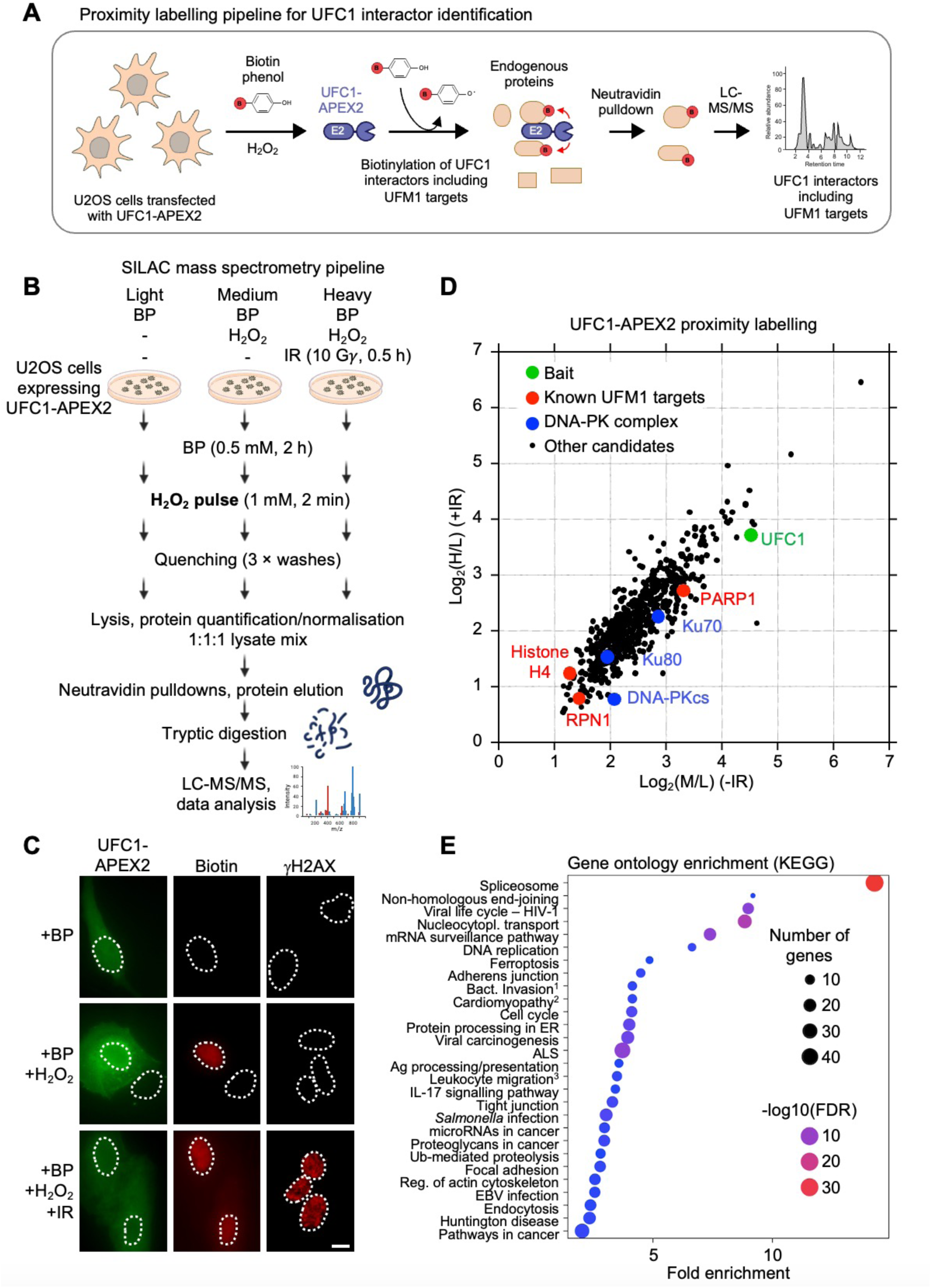
Proximity labelling identifies putative UFMylation targets across a wide range of pathways including NHEJ. (**A**) Schematic of proximity labelling strategy for UFM1 target identification. The engineered ascorbate peroxidase APEX2 was fused to the UFM1 E2 enzyme UFC1 and ectopically expressed in U2OS cells. Cells were treated with ionising radiation (IR, 10 Gy, ∼30 min) to activate DNA damage pathways. Biotinylation was initiated by 2 h pre-incubation with biotin phenol (BP, 0.5 mM), followed by a brief pulse of hydrogen peroxide (H₂O₂, 1 mM, 2 min) to catalyse proximity-dependent labelling. Biotinylated proteins, representing putative UFMylation substrates or interactors, were enriched via neutravidin pulldown and identified by liquid chromatography tandem mass spectrometry (LC-MS/MS). (**B**) Experimental workflow for SILAC-based quantitative proteomics of UFC1-APEX2 proximity-labelled interactors. U2OS cells stably expressing UFC1-APEX2 were cultured in light, medium, or heavy SILAC media corresponding to: (1) negative control (biotin phenol (BP) only, no H_2_O_2_; light), (2) APEX2-catalysed biotinylation without DNA damage (medium), and (3) APEX2-catalysed biotinylation following ionising radiation (IR, 10 Gy, ∼30 min; heavy). Following biotinylation, lysates from each condition were combined at a 1:1:1 ratio and subjected to neutravidin affinity purification. Enriched proteins were digested with trypsin and analysed by LC-MS/MS. (**C**) Fluorescence staining of U2OS cells transiently transfected with UFC1-2HA-APEX2, showing cytoplasmic as well as nuclear localisation. Biotinylation was only detected in the presence of BP, H_2_O_2_ and APEX2-fused UFC1. γH2AX was only detectable in the +IR condition. Detection of APEX2-fused UFC1 was by anti-HA, of biotinylated targets by streptavidin coupled to Alexa Fluor 594 and of γH2AX by a phospho-specific antibody. Dashed white lines mark nuclei outlines according to DAPI staining. Scale bar represents 10 μm. (**D**) Putative UFM1 targets identified via proximity labelling of APEX2-tagged UFC1 with a log2 ratio ≥ 1 in the M/L (-IR) or H/L (+IR) SILAC condition as described in (B). The bait (UFC1) is highlighted in green, known UFMylation targets in red, and select candidates of interest as part of the DNA-PK complex in blue. (**E**) Top gene ontology enrichments (KEGG) for putative UFMylation targets identified by proximity labelling, selected as described in (D). ^1^: Bacterial invasion of epithelial cells; ^2^: Arrhythmogenic right ventricular cardiomyopathy; ^3^: Leukocyte transendothelial migration. Abbreviations: ALS: amyotrophic lateral sclerosis; BP: biotin phenol; EBV: Epstein-Barr virus; ER: endoplasmic reticulum; FDR: false discovery rate; H: heavy; IR: ionising radiation; L: light; LC-MS/MS: liquid chromatography tandem mass spectrometry; M: medium; SILAC: stable isotope labelling by amino acids in cell culture; nucleocytopl.: nucleocytoplasmic; reg.: regulation; Ub: ubiquitin.

To probe how DSBs influence the UFMylation landscape, we performed APEX2-based proximity labelling in a stable isotope labelling by amino acids in cell culture (SILAC) format, integrating three conditions: (**1**) a negative control using biotin-phenol only (light), (**2**) active APEX2 labelling without DNA damage (medium), and (**3**) APEX2 labelling following IR-induced DNA damage (heavy) (Fig. 5B). Robust biotinylation was observed in conditions where APEX2-UFC1 was expressed and activated (+BP, +H_2_O_2_ ± IR, Fig. 5C, left and middle column), validating the inducibility of the system. Moreover, γH2AX induction confirmed detectable DNA damage exclusively in the IR-treated cells (Fig. 5C, right column), supporting the experimental design of the screening set-up.

Quantitative mass spectrometry identified >600 proteins significantly enriched in proximity to UFC1 under either basal (M/L) or DNA damage (H/L) conditions (Supplementary Table S2). UFC1 itself was one of the most enriched hits in both conditions (Fig. 5D, highlighted in green), as expected. Importantly, several previously validated UFMylation substrates, including histone H4^26,27^, PARP1^28^, and RPN1^69^ were enriched (Fig. 5D, highlighted in red), underpinning the validity of the labelling approach.

Gene ontology analysis revealed strong enrichment for a broad array of processes and pathways (Fig. 5E). These included both canonical UFM1-regulated pathways, such as ER-associated functions^14^, and a spectrum of emerging processes and clinical manifestations in UFM1 biology, including immunity, particularly in the context of Epstein-Barr virus infection^58,70,71^, ubiquitin-mediated proteolysis^53,72,73^, DNA replication^28–30^, ferroptosis^74^, cardiomyopathy^75,76^, amyotrophic lateral sclerosis (ALS)^59,77^, and cell cycle regulation^54–56^. The molecular details of how UFMylation regulates many of these processes remain to be elucidated, with our dataset providing an entry point for mechanistic dissection. Intriguingly, several pathways not previously linked to UFM1 were amongst the most enriched, including the spliceosome (Fig. 5E), aligning with our earlier UFM1 receptor screen (Fig. 3B), and pointing toward a possible new avenue for functional investigation. Notably, NHEJ was again amongst the most highly enriched ontologies, with all components of the DNA-PK holoenzyme – DNA-PKcs, Ku70, and Ku80 – being enriched (Fig. 5D, highlighted in blue). Together, these data establish the APEX2-UFC1 proximity labelling platform as a valid approach for mapping UFM1 substrate candidates and reinforce the notion that UFMylation intersects with a diverse range of cellular functions. Focusing on our goal of elucidating how UFMylation facilitates NHEJ, we next investigated the DNA-PK complex as a UFMylation target candidate.

### UFMylation of the DNA-PK complex establishes a UFM1-Ku70-XRCC4 axis linked to NHEJ

To validate posttranslational modification of the DNA-PK complex with UFM1, we immunoprecipitated DNA-PKcs and detected a high-molecular-weight band with an α-UFM1 antibody, migrating at a size consistent with UFMylated DNA-PKcs. Overexpression of the deUFMylase UFSP2 reduced the intensity of the band, supporting the notion that DNA-PKcs is modified by UFM1 (Fig. S13A). Furthermore, co-immunoprecipitation experiments demonstrated interactions between DNA-PKcs and the UFM1 E3 ligase UFL1 – both with exogenously expressed and endogenous UFL1 – alongside the Ku70:Ku80 heterodimer (Fig. S13B-C), further implicating the DNA-PK complex as a bona fide UFMylation substrate. To assess whether the Ku subunits themselves are UFMylated, we performed GFP pull-downs in HEK293T cells transiently expressing GFP-tagged Ku70 or Ku80, along with overexpression of the UFMylation machinery – an established strategy to detect UFMylation events^53^. In both cases, slower-migrating bands were detected with α-UFM1 immunoblotting, consistent with mono-UFMylation of Ku70 and Ku80. These bands were abolished upon co-expression of UFSP2, indicating UFMylation of both Ku subunits (Fig. S13D-E).

To identify the UFMylation sites within the DNA-PK complex, we reconstituted the reaction *in vitro* using purified components of the UFMylation cascade, containing the UBA5 E1, UFC1 E2, and UFL1:DDRGK1 E3 complex, as previously described^78^. In the presence of ATP, we observed slower-migrating forms of Ku70 and Ku80 when added either as a DNA-bound heterodimer or in complex with DNA-PKcs, consistent with UFMylation of the proteins *in vitro* (Fig. 6A). MS analysis of the modified gel bands identified several putative UFMylation sites, with lysine 526 (K526) on Ku70 emerging as the highest-confidence site, based on spectral evidence and reproducible ATP-dependence across replicates (Fig. 6B). To validate K526 as a bona fide UFMylation site in cells, we compared UFMylation levels of wild-type GFP-Ku70 and a K526R mutant. The K526R mutant showed a marked reduction in UFMylation, confirming K526 as a key UFM1 attachment site on Ku70 in cells (Fig. 6C). Given its spatial positioning within a previously reported structure of the NHEJ complex bound to DNA^79^, we reasoned that amongst the UFM1-binding regions identified, the IDR of XRCC4 spanning residues 256-263, a region within reach of the site without involving major rearrangements of the structured domains, was a likely candidate to function as a reader of K526-linked UFMylation (Fig. 6D). To test this, we performed GFP-Ku70 pulldown assays. The non-UFMylatable Ku70 K526R mutant co-precipitated markedly less XRCC4 compared to wild-type Ku70 (Fig. 6E), supporting a model in which XRCC4 engages UFMylated Ku70 through its UFM1-binding module in the IDR. Since the Ku heterodimer is amongst the first responders to DSBs – recruiting DNA-PKcs and acting as a scaffold for downstream NHEJ components – we hypothesised that UFMylation at Ku70 K526 facilitates XRCC4 recruitment via its UFM1-binding interface in the IDR. To test this, we performed laser micro-irradiation experiments and found that XRCC4 lacking its IDR UFM1-binding region was significantly impaired in its recruitment to DNA damage sites (Fig. 6F). Together, these data confirm UFMylation of the DNA-PK complex, validate our screening results, and indicate Ku70 K526 as a UFMylation site that is recognised by XRCC4’s UFM1 receptor activity to ensure proper recruitment to damaged chromatin.

**Figure 6.**
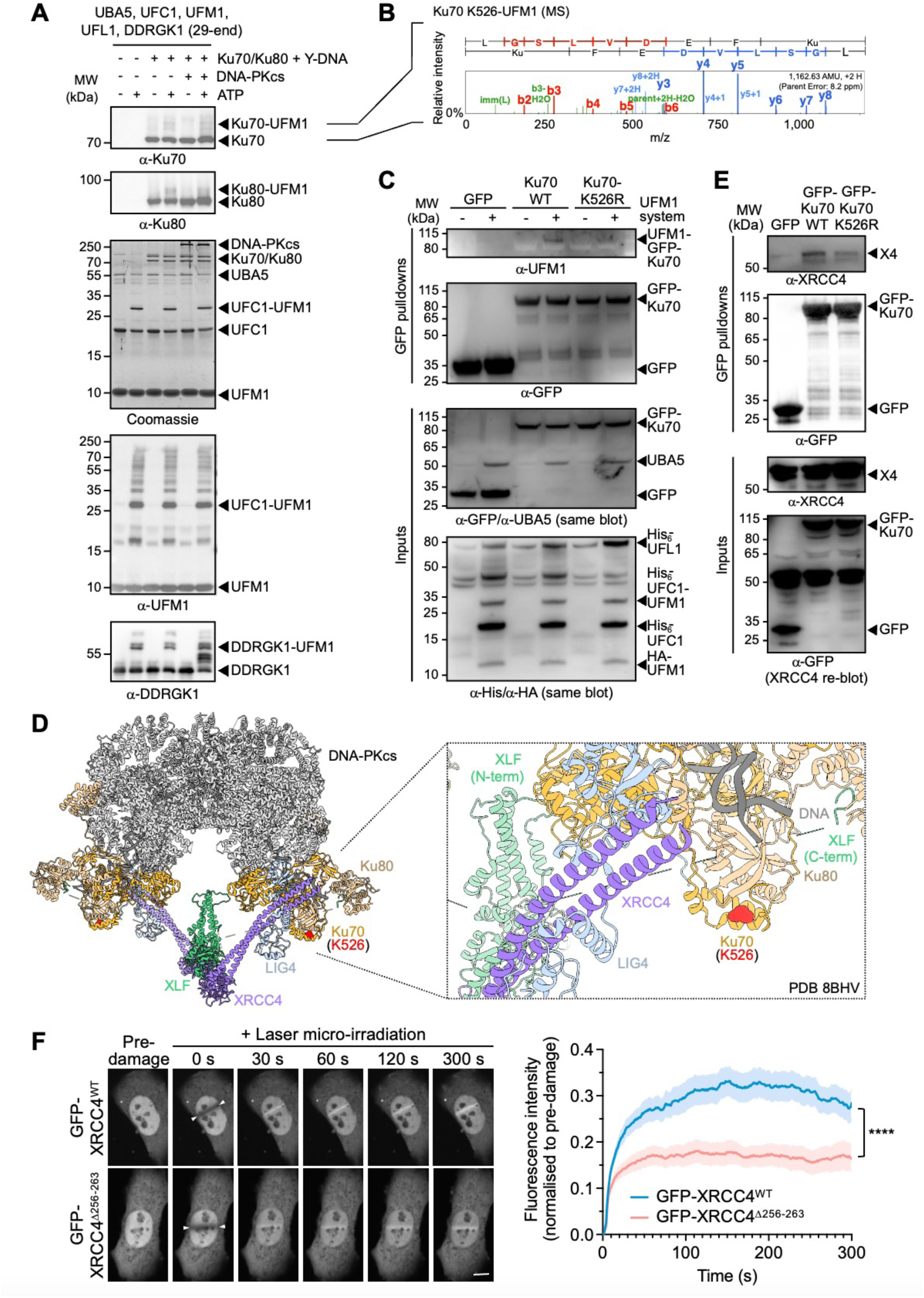
Ku70 K526 is UFMylated and recognised by XRCC4’s C-terminal IDR. (**A**) *In vitro* UFMylation assay using purified UFMylation machinery with recombinant DNA-PK complex (Ku70/80 on a DNA substrate ± DNA-PKcs) demonstrating UFMylation of the Ku complex. (**B**) Liquid chromatography tandem mass spectrometry (LC-MS/MS) example spectrum of Ku70 K526 UFMylation identified in multiple experiments only in the presence of ATP. Spectra were inspected using ScaffoldPTM with b-ions shown in red, y-ions in blue. (**C**) GFP-pulldowns of Ku70, wildtype (WT) or a K-to-R mutant version (K526R), from whole cell extracts, showing an α-UFM1-detectable band migrating at a molecular weight consistent with Ku70 UFMylation in the presence of ectopically expressed UFM1 system components (UFM1, UBA5, UFC1, UFL1 and DDRGK1), which was reduced in the Ku70 K526R mutant. (**D**) Structural representation of the NHEJ machinery (PDB 8BHV) highlighting the location of Ku70 K526 as a targeted UFMylation site in relation to neighbouring NHEJ factors including XRCC4 and XLF. (**E**) GFP-pulldowns of Ku70 (WT or K526R), from whole cell extracts obtained from HEK293T cells, showing decreased co-precipitation of XRCC4 in the mutant. (**F**) Representative images (left) pre-damage, during laser micro-irradiation, and 30 s, 60 s, 120 s and 300 s post-damage for U2OS XRCC4 knock-out cells transfected with the indicated GFP-tagged XRCC4 plasmids. Mean GFP fluorescence across the micro-irradiated site (normalised to background nuclear GFP fluorescence) with fluorescence normalised to 1 at t=0 (pre-damage) for each nucleus. Shaded area represents the error (±SEM), **** denotes *p*<0.0001 (unpaired t-test of means across the time course), scale bar represents 10 μm. White arrowheads mark the location of the end points of the laser micro-irradiated lines. Abbreviations: α: anti; IDR: intrinsically disordered region; LC-MS/MS: liquid chromatography tandem mass spectrometry; MW: molecular weight; WT: wildtype; X4: XRCC4.

### UFMylation promotes recruitment of NHEJ components to damaged chromatin

Building on our identification of a UFM1-interacting region on XRCC4 important for its recruitment to DNA damage sites, and additional UFMylation sites and UFM1-binding motifs in several key NHEJ components, we hypothesised that UFMylation facilitates the engagement of NHEJ factors with damaged chromatin more broadly. To test this, we perturbed global UFMylation levels and assessed chromatin association of core NHEJ proteins, previously shown to be covalently modified by UFM1 and/or interacting with it non-covalently. Upon treatment with calicheamicin, a potent inducer of DSBs^80,81^, we evaluated NHEJ factor loading onto chromatin. These experiments aimed to determine whether UFMylation functions as a general mechanism to regulate NHEJ complex assembly at sites of damage.

Consistent with our hypothesis, inducible depletion of the deUFMylase UFSP2 in U2OS cells – engineered via degron tagging at the endogenous *UFSP2* locus – resulted in a marked increase in the DNA damage-induced chromatin association of all NHEJ components featuring UFMylation sites and/or UFM1-interacting modules, concomitant with elevated global and chromatin-associated UFMylation (Fig. 7A). Notably, Ku, the earliest factor recruited during NHEJ, displayed >2-fold enrichment on chromatin at the earliest time point, 0 h post-calicheamicin treatment, with levels gradually declining yet remaining elevated even 18 hours after drug removal. DNA-PKcs, which is recruited by Ku, similarly showed robust and sustained chromatin association from the earliest time point onwards. XRCC4 and XLF – recruited at later stages – displayed the most significant increases in chromatin engagement at slightly later time points, peaking at 2-3 h post-damage, in UFSP2-depleted cells (Fig. 7A). These findings link the post-translational regulation of NHEJ factors by UFMylation to enhanced assembly and/or stability of the NHEJ machinery at sites of damage and reveal factor-specific kinetic differences that may reflect distinct modes of UFM1-dependent regulation across the repair timeline.

**Figure 7.**
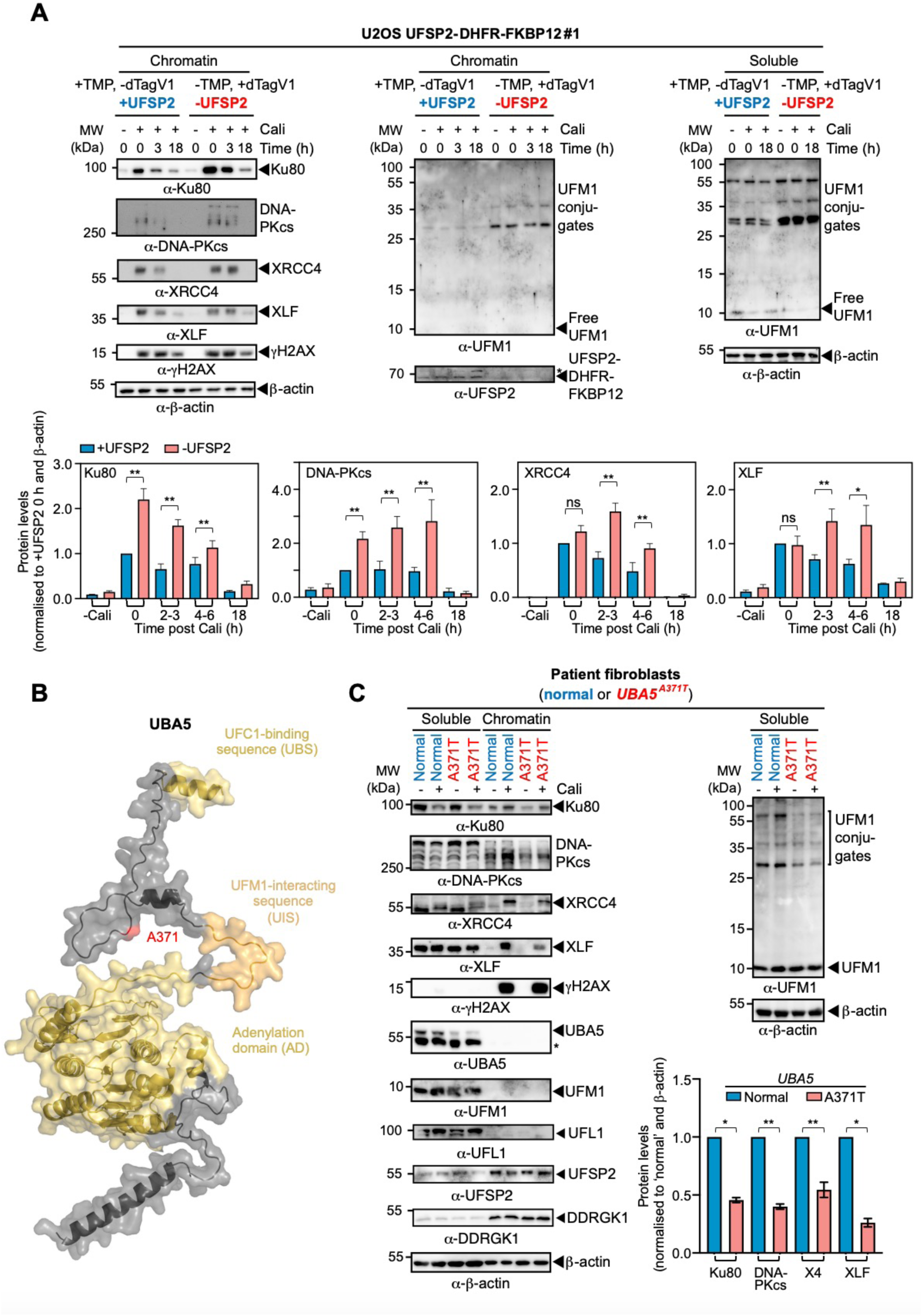
UFMylation promotes DNA damage-induced chromatin engagement of NHEJ components, linking NHEJ alterations to patients with UFM1 pathway mutations. (**A**) Chromatin association of key NHEJ factors in U2OS cells integrating a *UFSP2-DHFR-FKBP12* double-degron at the endogenous locus – induced by addition of dTagv1 and depletion of Trimethoprim (TMP) – in response to calicheamicin (Cali, 5 nM, 1 h), or not (non-treated, NT) at the indicated time points and compared to non-damaged conditions. Protein levels were quantified from immunoblots, with a representative replicate shown. Error bars represent means ± SEM of n=4 (18 h), n=8 (4-6 h, XLF -Cali), or n=10 (NT (apart from XLF -Cali), 0 h, 2-3 h) replicates, pooled from two technical and two to five biological replicates. (**B**) Alphafold model of UBA5 (Hs UBA5 AF-Q9GZZ9-F1-v4), illustrating the location of alanine 371 (A371), a residue mutated in patients with hereditary UFMylation syndromes. (**C**) Characterisation of levels and chromatin association of UFM1 pathway components and key NHEJ factors in primary fibroblasts obtained from a patient, compound heterozygous for *UBA5* mutations, carrying a missense mutation (A371T) on one, and a loss-of-function mutation on the other, allele, compared to fibroblasts obtained from a healthy individual (normal). Fibroblasts were treated, or not, with calicheamicin (Cali, 5 nM, 1 h) before their processing for chromatin fractionation. Protein levels were quantified from immunoblots, with a representative replicate shown. Error bars represent means ± SEM of n=6 (Ku80, XLF) or n=9 (DNA-PKcs, XRCC4) replicates, pooled from three technical and two to three biological replicates. Abbreviations: α: anti; AD: adenylation domain; Cali: calicheamicin; MW: molecular weight; SD: standard deviation; TMP: Trimethoprim; UBS: UFC1-binding sequence; UIS: UFM1-interacting sequence; X4: XRCC4.

### Defects in NHEJ complex accrual in cells from a patient with UFM1 pathway mutations

To examine whether these findings were clinically relevant, we turned to primary cells from a patient compound heterozygous for mutations in *UBA5*, the E1 enzyme in the UFM1 cascade. The patient carried a missense mutation (A371T) – the most frequent *UBA5* variant in the general population (allele frequency 0.27%)^82^ – on one allele, and a loss-of-function mutation on the other^83^. The A371T substitution lies within an IDR situated between the UFM1-interacting sequence (UIS) and UFC1-binding sequence (UBS) (Fig. 7B), as part of a short, relatively conserved, sequence stretch critical for UFM1 transfer^84^. Consistent with its classification as a hypomorphic allele, UBA5 protein levels were markedly reduced in patient-derived fibroblasts, leading to impaired global UFMylation while levels of free UFM1 remained unchanged (Fig. 7C). Strikingly, these fibroblasts exhibited substantial defects in the chromatin association of all key NHEJ factors previously identified as harbouring UFMylation sites or UFM1-interacting modules, with reductions of ∼50-70% compared to controls following DNA damage (Fig. 7C). These findings underscore UFMylation as an essential posttranslational modification required for the efficient recruitment and coordinated assembly of the NHEJ repair complex on damaged chromatin.

## DISCUSSION

UFMylation has emerged as a posttranslational modification (PTM) crucial for maintaining cellular homeostasis. Despite its biological significance, relatively few substrates and even fewer receptors that mediate UFM1-dependent signalling have been characterised. Here, we employ a multifaceted, bottom-up discovery strategy – integrating loss-of-function UBL screening, genetic code expansion with photo-crosslinking, and proximity labelling in cells – to trace a UFM1-mediated phenotype to its downstream receptor and ultimately to the upstream substrate and UFMylation site. This approach not only validates the DNA-PK complex as a direct UFMylation target but also uncovers mechanistic insights into how reader:writer interactions facilitate engagement of NHEJ factors with DNA damage sites, a central prerequisite for effective end-joining repair.

The identified interactions are mediated via a surface on UFM1 formed by its α-β groove analogous to a conserved receptor-binding interface on SUMO paralogues^45,64^. One of the UFM1-interacting regions on XRCC4 contains a short motif (180-FILVL-184) that shares key features with canonical UFM1-interacting sequences – namely, an aromatic residue followed by branched aliphatic side chains – but notably lacks the acidic residues found in these sequences in UBA5 (WGIEL)^42^ and DDRGK1 (FVVEE)^85^. Several additional UFM1-interacting regions we identified across XRCC4 and other NHEJ factors lacked conserved UIS features altogether, suggesting that UFM1’s α-β groove engages its receptor proteins through diverse binding modes. The presence of multiple UFM1-binding regions and UFMylation sites across several subunits of the NHEJ complex – together with our functional data showing enhanced or decreased chromatin association of these same factors upon global UFMylation increase or decrease – supports a model in which UFMylation acts as a multivalent assembly signal. This is reminiscent of SUMO’s role as a “molecular glue” that promotes higher-order protein complexes^86^. Whether UFM1 also drives phase separation in this context remains an open question, but the presence of a functional UFM1-binding region within the intrinsically disordered region (IDR) of XRCC4 – a region recently implicated in phase separation during NHEJ^87^ – makes this an intriguing possibility.

We further identify a UFM1-binding region on a functionally validated area of XRCC4’s head domain, which overlaps with a non-conventional SUMO-interacting module and also binds to XLF^64,65^. This raises compelling questions about potential crosstalk or competition between UFM1, SUMO and other pathways at this regulatory hotspot – particularly in modulating complex synapsis at DSBs – and how these pathways may fine-tune NHEJ depending on cellular context or the nature of the lesion.

Given previous reports implicating UFM1 in HR, our findings suggest UFMylation plays broader roles across DSB repair pathways. Our siUBL screen reveals UFM1 as an important player in the formation of 53BP1 and conjugated ubiquitin IRIF – processes intimately tied to DSB repair pathway choice^4^. Notably, several proteins central to pathway decision-making, including 53BP1 and RIF1 emerged as candidate UFMylation substrates in our proximity labelling screen (Supplementary Table S2), underscoring UFM1’s potential role at the HR-NHEJ crossroad. Moreover, our observation that marked levels of the deUFMylase UFSP2 associate with chromatin (Fig. 7), raises the possibility that spatially resolved deUFMylation by UFSP2 may influence repair pathway selection. Such regulation may be contingent on chromatin context, break type, cell cycle phase, and/or tissue identity, and could define distinct temporal windows of UFM1 site functionality. Whether and how deUFMylation actively restricts NHEJ complex assembly and how this intersects with other mechanisms regulating NHEJ accrual at DNA damage sites, such as recently identified roles for DNA-PKcs and NEDDylation/FBXL12 pathways^88^, remain compelling questions for future investigation. Interestingly, UFMylation has also been implicated in telomere biology^89^, where suppression of end-joining is critical to prevent telomeric fusions. Whether UFSP2 localisation at telomeres contributes to constraining aberrant NHEJ in this context presents an intriguing avenue for future investigation.

Beyond basic DNA repair biology, the dysregulation of UFMylation has been linked to cancer, acting both to suppress and drive carcinogenesis in different contexts^14,90^. Given the central role of DDR dysfunction in cancer progression, it will be important to explore how dysregulated NHEJ induced by alterations in UFMylation might modulate genome maintenance in different tumour settings, and whether intervention of the UFM1 pathway could offer therapeutic leverage, especially as pharmaceutical strategies targeting the UFMylation machinery begin to garner interest^90^.

Our study reveals a functional link between UFMylation disorders and NHEJ dysregulation. These findings are especially relevant for neurodevelopment, as phenotypic overlaps – such as microcephaly – might indicate shared underlying mechanisms. Understanding how UFMylation-dependent regulation of NHEJ manifests in different cellular contexts, in particular neural cell types, represents a key priority for uncovering disease pathogenesis and developing targeted interventions in the future.

## ACKNOWLEDGEMENTS

We thank Stephen P. Jackson (CRUK Cambridge Institute, UK) for his generous support during C.K.S.’s time in the group at the Gurdon Institute (Cambridge, UK). Further thanks go to Sébastien Britton and Patrick Calsou (both at the IPBS, Tolouse, France) for help with setting up chromatin pellet experiments and discussion, Michał Malewicz (currently at Lukasiewicz Research Network - Polish Center for Technology Development, Wroclaw, Poland) for sending the U2OS XRCC4 knockout cells, Tom Blundell (University of Cambridge, UK) and Qian Wu (University of Leeds, UK) for reagents and advice on NHEJ factor protein purification, and members of the CRUK Manchester Institute proteomics and mass spectrometry units for help with analysis. The authors acknowledge funding by the BBSRC (David Phillips BBSRC Fellowship BB/N019997/1 to C.K.S., DTP studentships), the MRC (research grants MR/X008754/1 to C.K.S., MR/X00029X/1 to A.K.C.), the Israel Science Foundation (research grant number 491/2021 to R.W.), the Israel Cancer Research Fund (award ID 21-113-PG to R.W.), CRUK (R122321 grant to D.L.S.), through a PDS award (the University of Manchester), by a Lister Institute of Preventative Medicine Prize (to A.K.C.) and a FEBS Return-to-Europe Fellowship (to C.K.S.). Research in the Steve Jackson lab at the Gurdon Institute was supported by Cancer Research UK Programme Grant C6/A11224 and ERC Advanced Researcher Grant DDRREAM (grant agreement number 268536).

## MATERIALS AND METHODS

### METHOD DETAILS

#### Cell culture

U2OS, U2OS XRCC4 knock-out – a kind gift from Michał Malewicz (currently at Lukasiewicz Research Network – Polish Center for Technology Development, Wroclaw, Poland) – HEK293T, and patient-derived cells (CB18-0280 fibroblasts from a healthy donor and CB24-0108 fibroblasts from a patient carrying the *UBA5* p.A371T mutation, a kind gift from Shane McKee^83^) were cultured in Dulbecco’s Modified Eagle Medium (DMEM, Sigma-Aldrich), supplemented with 10% foetal bovine serum (v/v) (FBS, Sigma-Aldrich) and 1% L-glutamine/penicillin/streptomycin (v/v) (100 ×, Gibco) or 1% glutamine (v/v) (100 ×, Gibco). Cultures were maintained at 37 °C in a humidified 5% CO_2_ atmosphere and routinely tested for mycoplasma. IR treatments were performed using a CellRad Faxitron instrument (Faxitron Bioptics, LLC). U2OS cell lines with a FKBP12-DHFR double degron inserted at the 3’-end of the *UFSP2* locus were generated as described in the Supplementary Methods and cultured in DMEM supplemented with 10% FBS (v/v), 1% L-glutamine/penicillin/streptomycin (v/v) and 10 μM Trimethoprim (TMP, MP Biomedicals). For double-degron treatment TMP was excluded from the media and dTagV1 (#6194, Tocris) was included at 500 nM. Ouabain (Tocris) was used at 10 μM working concentration for screening clones.

#### siRNA and plasmid transfections

siRNA oligonucleotides were designed using the Dharmacon Design Centre and purchased from Eurofins Genomics unless stated otherwise. For siRNA and plasmid co-transfections, 350,000-450,000 cells were seeded on 60 mm plastic dishes. Sterile coverslips were included in 60 mm dishes for immunofluorescence assays. Cells were transfected with siRNAs the following day to a final concentration of approximately 60 nM using Lipofectamine RNAiMAX (Life Technologies) according to the manufacturer’s guidelines. For plasmid transfections, cells were transfected using FuGENE 6 (Promega) or Lipofectamine 2000/3000 (Life Technologies), following the manufacturer’s guidelines. For siRNA and DNA co-transfections, plasmid transfections were performed around 8 h after siRNA treatment. siRNA target sequences and plasmids used for transfections are available in Supplementary Table S3.

#### Traffic Light Reporter (TLR) system

The Traffic Light Reporter (TLR) system enables simultaneous quantification of HR and mutEJ at a DSB. The reporter cassette contains a mutant GFP sequence interrupted by an I-SceI endonuclease recognition site, followed downstream by an mCherry sequence placed in a +2 out-of-frame position. Upon expression of I-SceI, DSB induction leads to one of two outcomes depending on the repair pathway engaged: HR restores functional GFP using a co-transfected donor plasmid encoding a truncated, repair-competent GFP fragment, while mutEJ can result in a 2 bp frameshift and activation of mCherry expression^16^. U2OS cells stably carrying the TLR cassette^17^ were transfected with the indicated siRNAs or treated with small-molecule inhibitors (DNA-PK inhibitor (NU7441, Tocris Bioscience, 3 μM 72 h; proteasome inhibitor MG132, 0.1 μM, 48 h; ATM inhibitor KU55933, 10 μM, 72 h; DNA-PK inhibitor AZD7648, 20 μM, 72 h; ATM inhibitor AZD1390, Cambridge Bioscience, 20 μM, 72 h). Approximately 8 h later, cells were co-transfected with expression plasmids encoding I-SceI and an infrared fluorescent protein (IFP), alongside a donor construct carrying blue fluorescent protein (BFP). Cells were collected ∼72 h post-siRNA transfection and analysed by four-colour flow cytometry on a BD LSR Fortessa (BD Biosciences). At least 10,000 IFP⁺/BFP⁺ double-positive cells were recorded per condition. Data were analysed using FlowJo (TreeStar), and the frequency of GFP⁺ and mCherry⁺ cells was used to quantify HR and mutEJ efficiencies, respectively. Repair efficiencies were normalised to control siRNA targeting firefly luciferase (siCTRL). To account for potential siUBL-induced alterations in cell cycle distribution, HR values were further normalised to the fraction of cells in S/G2 phases, as determined by propidium iodide (PI)-based DNA content profiling. For this, siRNA-treated U2OS cells were fixed in 70% ethanol, incubated with RNase A (250 μg/mL) and propidium iodide (10 μg/mL) for 30 min at 37 °C, and analysed on a FACSCalibur flow cytometer (BD Biosciences) using CellQuest software. Cell-cycle profiles were processed in FlowJo using the Watson Pragmatic and Dean-Jett-Fox algorithms.

#### Automated high-throughput/high-content microscopy for IRIF quantification

Automated high-throughput/high-content microscopy of IRIF kinetics was performed as described previously^17^. ∼48 h after siRNA transfection, U2OS cells were seeded into 96-well plates (Cell Carrier, Perkin Elmer, cat# 6005550, 15,000 cells/well). The following day, cells were irradiated using a Faxitron CellRad irradiator (Faxitron Bioptics, LLC, Tucson, AZ, USA) or directly processed to retrieve non-treated reference plates. Plates were fixed 30 min after treatment with IR (2 Gy) and stained with the respective antibodies and DAPI. A spinning-disk Perkin Elmer Opera platform equipped with a 20 × water immersion objective (0.7 numerical aperture, Olympus) was employed to acquire 6-10 confocal images (fields) for each well in a single optimised focal plane. The images were analysed using an optimised spot detection script operated by an integrated software package (Acapella, Perkin Elmer). DAPI was used to segment nuclei and create a nuclear mask. Automated mask transferral to γH2AX-, 53BP1- or conjugated ubiquitin-(recognised by the FK2 antibody) stained images allowed detection of intensities and IRIF specifically within the nuclear areas of the detected cells.

For validation of siUFM1 effects with deconvoluted siRNAs, ∼48 h post-siRNA transfection, U2OS cells were seeded into 96-well plates (Perkin Elmer; cat# 6005550) at a density of 17,500-25,000 cells per well and incubated overnight. Selected plates were treated with IR (4 Gy). Cells were fixed 30 min post-IR using 2% paraformaldehyde in phosphate-buffered saline (PBS) at room temperature (RT, 20 min), permeabilised with 0.5% Triton X-100 in PBS (15 min, RT), and stored in 0.1% Tween-20 in PBS (PBST) at 4 °C. PBST washes (2-3 times) were performed before and after each subsequent step. Cells were stained with primary antibodies, followed by appropriate Alexa Fluor-conjugated anti-rabbit or anti-mouse secondary antibodies, in combination with DAPI for nuclear co-staining. After each antibody incubation, cells were washed with PBST (2-3 times). Images were acquired using an Operetta high-content imaging system (Perkin Elmer) with a 63 × water immersion objective.

#### Circos plot analysis

siUBLs ranking within the top 25^th^ percentile, based on pooled values across all DDR read-outs and normalised to 100% for siCTRL, were selected for visualisation. In the resulting Circos plot, siUBLs are displayed on the right hemisphere, arranged clockwise by decreasing cumulative DDR impact. The outermost track indicates the specific DDR read-outs each siUBL scored in, with individual elements ordered clockwise by decreasing contribution to the overall DDR phenotype. The width of each segment reflects the magnitude of the observed phenotype. DDR read-out modules are arranged on the left hemisphere, with their corresponding outer track indicating the siUBLs scoring in each respective read-out. These modules are similarly ordered clockwise by decreasing individual siUBL effect size, and the segment width corresponding to phenotype strength^91^.

#### Random plasmid integration

Random plasmid integration assays were performed as previously described^19^ with minor modifications. Briefly, after siRNA treatment, U2OS cells were transfected with BamHI-XhoI linearised pEGFP-C1 plasmid (Clontech). The following day, cells were collected, counted and plated on two 150 mm plates for colony formation and 60 mm dishes for transfection efficiency checking. The next day, 0.5 mg/mL G418 was added to one of the 150 mm plates and the cells on the 60 mm dishes were used to assess GFP transfection efficiency by flow cytometry using a BD LSR Fortessa cell analyser (BD Biosciences). At least, 10,000 cells were assessed per siRNA condition. Cells on the 150 mm plates were incubated for 10-14 days at 37 °C to allow for colony formation. Colonies were stained with 0.5% crystal violet/20% ethanol and counted. Random plasmid integration events were normalised to transfection and plating efficiencies.

#### Clonogenic survival assays

1000-5000 siRNA-treated U2OS cells were plated in each well of 24-well plates. ∼16 h later, cells were exposed to IR (1-3 Gy) and incubated for 10-14 days at 37 °C to allow colony formation. Colonies were stained with 0.5% crystal violet/20% ethanol and counted. Results were normalised to plating efficiencies.

#### UFM1* probe photo-crosslinking assays

Crosslinking assays were performed as described previously^43,47,64^ with minor modifications. For the screen, HEK293T cells were harvested 30 min after exposure to IR (10 Gy) and lysed in PBS (137 mM NaCl, 2.7 mM KCl, 10 mM Na₂HPO₄, 1.8 mM KH₂PO₄, pH 7.4) containing 1 × cOmplete EDTA-free protease inhibitors (Roche), using three cycles of sonication (30 s on/off, 30% power). 20 µM singly C-terminally biotinylated UFM1 containing BpF at amino acid position F35 (UFM1*) was incubated with 2-10 mg/mL HEK293T cell extracts for 30 min at 4 °C in 0.2 mL thin-walled polypropylene PCR tubes (Greiner Bio-One). Samples were then irradiated for 2 h with long-wave UV light (λ = 365 nm; Herolab UV-16 L, 8 W) at a 6 mm distance. Aliquots were collected before and after UV exposure. Proteins crosslinked to UFM1 were enriched via streptavidin sepharose pulldown (Cytiva, 17-5113-01). Beads were washed five times with 500 µL of each of the following buffers using spin columns (Pierce Micro-Spin, cat#10510824): buffer A (8 M urea, 2% SDS, 100 mM Tris-HCl pH 8, 200 mM NaCl), buffer B (8 M urea, 0.2% SDS, 10% ethanol, 10% isopropanol, 100 mM Tris-HCl pH 8, 1.2 M NaCl), buffer C (same as buffer B, pH 5), buffer D (same as buffer B, pH 9), buffer E (100 mM Tris-HCl pH 8, 200 mM NaCl), and buffer F (50 mM NH₄HCO₃). Streptavidin beads were transferred to protein LoBind tubes (Eppendorf) and bound proteins eluted with 2 × SDS Laemmli buffer (120 mM Tris-HCl pH 6.8, 4% SDS, 20% glycerol, 0.02% bromophenol blue and 2.5%-mercaptoethanol) for SDS-PAGE analysis. Samples were excised from fully resolved SDS-PAGE gels, subjected to in-gel tryptic digestion, and analysed using a Vanquish Neo HPLC system coupled to an Orbitrap Lumos mass spectrometer via an EasySpray source (Thermo). The HPLC utilised mobile phases A (0.1% formic acid in water) and mobile phase B (80% acetonitrile, 0.1% formic acid). The MS was operated in data-independent acquisition (DIA) mode. DIA data was processed using Spectronaut 17 (Biognosys) in direct DIA mode using oxidation (M), deamidation (NQ) and acetylation (N-terminal) as variable modifications in a search against a human Uniprot database and only high-confidence matches were retained for downstream analysis. Gene ontology analyses were carried out for the most significantly enriched proteins (log2 fold change ≥ 0.58) using ShinyGO 0.80^92^ and KEGG^93^. For validation experiments, crosslinking was performed using 20 µM UFM1 F35-BpF with 20 µM of the recombinant interactor candidate or cell extract obtained from HEK293T cells, transiently transfected with the protein-of-interest.

#### Recombinant proteins, peptides and purification

UFM1 recombinant proteins incorporating the unnatural amino acid 4-benzoyl-(L)-phenylalanine (BpF) at position F35 (UFM1*) were expressed from a pGEX6P1 plasmid backbone in *E. coli* BL21(DE3) cells co-transformed with the plasmid pEVOL-pBpF, a gift from Peter G. Schultz (Addgene plasmid #31190; RRID: Addgene_31190)^44^. This plasmid encodes an orthogonal aminoacyl-tRNA synthetase/tRNA pair (BpFRS/BpFtRNACUA) derived from *Methanococcus jannaschii* that specifically incorporates BpF at the amber stop codon (UAG). Protein expression was induced in LB medium supplemented with 1 mM BpF, 0.05% (w/v) arabinose, and 250 µM IPTG overnight at 18 °C. Cells were lysed by sonication on ice and clarified by centrifugation at 20,000 x *g* for 20 min at 4 °C. Lysate was passed over a GSTrap column (Cytiva) and washed with 10 column volumes (CV) PBS supplemented with 1 mM DTT followed by equilibration with 10 CV Prescission/3C protease buffer (50 mM Tris-HCl pH 7.5, 150 mM NaCl, 1 mM EDTA, 1 mM DTT) before overnight on-column digestion of the GST tag with GST-3C protease (made in-house). The flow-through containing untagged UFM1 F35-BpF was collected. Biotinylation of the C-terminal cysteine was carried out using an EZ-Link biotin maleimide reaction kit (Sigma, #cat21901BID) with excess biotin removed using 7 kDa MWCO Zeba spin desalting columns (ThermoScientific, #cat89883) equilibrated with PBS.

UBA5, UFC1, and UFM1(ΔSC) for use in enzyme assays were expressed from a pOPINB vector with an N-terminal His_6_-3C tag. Protein expression was induced with 0.4 mM IPTG once transformed, growing *E. coli* (BL21(DE3)) had reached an OD_600_ between 0.5-0.7 in LB broth. Cells were harvested by centrifugation and re-suspended in lysis buffer (20 mM Tris-HCl pH 7.5, 500 mM NaCl, 20 mM imidazole, 0.1 % Triton-X100, 1 × EDTA-free cOmplete protease inhibitors (Roche)) and stored at −20 °C until use. Cells were lysed by sonication on ice and clarified by centrifugation at 20,000 x *g* for 20 min at 4 °C before filtering the lysate through a 0.45 µm PES filter. Lysate was passed over a HisTrap FF 1 ml column (Cytiva), washed with 5 CV wash buffer (20 mM Tris-HCl pH 7.5, 500 mM NaCl, 20 mM imidazole), and eluted with a linear gradient from 0-100% elution buffer (20 mM Tris-HCl pH 7.5, 500 mM NaCl, 500 mM imidazole) over 5 CV. For UBA5 and UFC1, fractions containing the His_6_-tagged protein were directly injected onto a Superdex 200 size-exclusion chromatography (SEC) column (Cytiva) and eluted by isocratic elution in 20 mM Tris pH 7.5, 100 mM NaCl, 1 mM DTT. Pure fractions (>90 % by Coomassie-stained SDS-PAGE) were pooled, concentrated in 3 kDa MWCO spin concentrators (Amicon), glycerol added to 10% (v/v), aliquoted, and stored at −70 °C. For UFM1(ΔSC), the N-terminal His_6_-tag was removed with His_6_-3C protease (made in-house) overnight at 4 °C while dialysing into 20 mM Tris-HCl pH 7.5, 100 mM NaCl before reverse purification over a HisTrap FF 1 mL column to remove the cleaved tag and 3C protease. Pure untagged UFM1(ΔSC), was further purified by SEC as above except for use of a Superdex 75 column (Cytiva).

For the E3 ligase UFL1:DDRGK1 complex, full-length UFL1 and truncated DDRGK1:DDRGK1 were expressed from a pRSFDuet1 dual expression plasmid with UFL1 inserted into MCS1 with an N-terminal His_6_-tag and DDRGK1 (29-314) in MCS2 with an N-terminal StrepII-tag. Protein expression, cell lysis, and HisTrap FF purification were carried out as above for UBA5 and UFC1 but following HisTrap elution, protein was injected onto a StrepTrap HP 1 mL column (Cytiva), washed with 20 mM Tris-HCl pH 7.5, 500 mM NaCl, 1 mM DTT and eluted in 5 mL wash buffer supplemented with 50 mM biotin. The UFL1:DDRGK1 complex was finally purified by SEC using a Superdex 200 column (Cytiva) as above for UBA5 and UFC1.

Ku70, Ku80 and DNA-PKcs proteins together with Y-shaped dsDNA were kind gifts from Amanda Chaplin (University of Leicester)^94^. His_6_-tagged XRCC4 expression plasmids (full-length XRCC4^95^, XRCC4^1-213^ (C-to-A: cysteines mutated to alanines)^96^, XRCC4^1-164^ (C-to-A: cysteines mutated to alanines)^97^), XLF^FL^ and XLF^1-224^ were generous gifts from Qian Wu (University of Leeds, UK) and Tom Blundell (University of Cambridge, UK). Recombinant proteins expressed from these plasmids were purified as described previously^96,98^.

XRCC4 C-terminal 200-334 (unlabelled and ^15^N-labelled) was expressed from a pOPINS vector with an N-terminal His_6_SUMO-tag. ^15^N-labelled proteins for NMR experiments were expressed in minimal medium supplemented with ^15^NH_4_Cl and purified equivalent to unlabelled protein. Expression, cell lysis, and HisTrap FF purification were carried out as above for UFM1 but following elution, His_6_-SENP1 SUMO protease (made in-house) was added whilst dialysing against 20 mM Tris-HCl pH 7.5, 100 mM NaCl, 20 mM imidazole. His-SENP1 and the His_6_SUMO tag were removed by reverse purification, with the flow-through containing untagged XRCC4^200-334^ that was further purified by SEC using a Superdex 75 column (Cytiva) as above for UFM1. Protein was buffer exchanged into Bis-Tris pH 6.5, 150 mM KCl, 1 mM EDTA, 1 mM DTT and concentrated to 400 µM or 200 µM (when titrating in unlabelled UFM1) for NMR experiments.

Plasmids for recombinant protein expression and purified peptides used in this study are available in Supplementary Table S3.

#### Immunoblotting, immunofluorescence, and immunoprecipitation

At the indicated time points, cells were harvested in lysis buffer containing 50 mM Tris-HCl pH 7.5, 2% sodium-dodecyl sulphate (SDS), 10 mM *N*-ethylmaleimide, 1 × EDTA-free cOmplete protease inhibitor (PI) cocktail (Roche) and 1 × phosphatase inhibitor (PPI) cocktail (50 mM NaF, 50 mM β-glycerophosphate, 10 mM Na_3_VO_4_, 250 mM sodium pyrophosphate). Whole cell extracts were incubated at 95 °C for 5 min, briefly spun down and sonicated at 30% power (QSonica Sonicators) for 50 s. Protein concentrations were determined using a Pierce BCA Assay kit (Thermo Fisher Scientific). Reactions were performed in triplicates. Absorbance was measured at 565 nm using a plate reader (Varioskan Lux, Thermo Fisher Scientific) and protein concentrations in μg/μL were calculated using a standard curve. To resolve protein bands, 30-60 μg of proteins were run on SDS-PAGE gels. Gels were then electroblotted onto PVDF membranes (GE Healthcare). PVDF membranes were blocked with 1-5% bovine serum albumin (BSA, Sigma-Aldrich) or 5% milk (w/v) in TBST (50 mM Tris-HCl pH 7.6, 150 mM NaCl, 0.1% Tween-20) at room temperature for 30-60 min prior to overnight incubation with primary antibody at 4 °C. After 3 washing steps of 10 min each with TBST, membranes were incubated under agitation with the appropriate secondary antibody for 1 h at RT. After incubation, membranes were washed three times for 15 min with TBST. Protein bands were visualised with ECL reagent (GE Healthcare) or Immobilon Forte (Sigma-Aldrich, used for pull-downs) using a ChemiDoc-Touch Imaging System (Bio-Rad).

For immunoblotting, the following primary antibodies were used against the indicated antigens: UFM1 (ab109305, Abcam, 1:500 to 1000), UBA5 (A304-115A, Bethyl, 1:1000), UFC1 (ab189252, Abcam, 1:2000), UFL1 (A303-456A, Bethyl Laboratories, 1:2000), DDRGK1/UFBP1 (21445-1-AP ProteinTech, 1:1000), UFSP2 (16999-1-AP Proteintech, 1:1000), XRCC4 (Santa Cruz, 1:500), Ku70 (10723-1-AP Proteintech, 1:1000), Ku80 (16389-1-AP Proteintech, 1:1000), LIG4 (ab193353 Abcam, 1:1000), XLF (2854 CST, 1:1000 or A300-730A Bethyl, 1:1000), γH2AX (05-636, Millipore, 1:10000), α-tubulin (14-4502-80, eBioscience, 1:5000), β-actin (A3854 Sigma, 1:20,000), H3-HRP (12648 CST, 1:5000), HA (MMS-101R, Biolegend, 1:5000), GFP (11814460001, Roche, 1:1000 or 2955 CST, 1:1000), DNA-PKcs (sc-5282, Santa Cruz, 1:250), FLAG (20543–1-AP, Proteintech, 1:2000 to 1:4000), His_6_-tag (652501 Biolegend, 1:1000) and HRP-conjugated streptavidin (43-4323, Thermo Fisher, 1:50,000). The following secondary antibodies were used: goat anti-rabbit IgG (31462, Invitrogen) and goat anti-mouse IgG (P0260, Dako) at 1:5000 to 1:10,000 dilution.

For immunofluorescence other than the siUBL IRIF evaluations, coverslips were fixed in 4% paraformaldehyde at RT for 15 min. After 3 washing steps of 5 min each using 1 × PBST, cells were permeabilised for 15 min in 0.5% Triton X-100. Following 3 washes of 5 min each in PBST, coverslips were inverted onto primary antibodies diluted in blocking buffer (5% BSA in PBST) and incubated overnight at 4 °C. Coverslips were then washed 3 × in PBST and incubated in secondary antibodies containing 1 μg/mL DAPI for 1 h at RT. Coverslips were washed 3 × in PBST, mounted on slides with Vectashield mounting media (H-1000, Vector Laboratories) and sealed with nail polish. Images were taken with an EVOS 7000 Imaging System (Thermo Fisher Scientific) using a 100 × oil lens and processed using ImageJ and Imaris (Bitplane) software packages. The following antibodies/reagents were used for staining: anti-53BP1 (NB100-034, Novus Biologicals, 1:100), anti-ubiquitin conjugates (FK2 antibody, BML-PW8810, Enzo Lifesciences, 1:100), anti-UFM1 (ab109305, Abcam, 1:100), anti-HA (16B12, MMS-101R, Biolegend, 1:200), anti-γH2AX JBW301 (05-636, Millipore, 1:100), as well as Alexa Fluor 488 goat-anti-rabbit (A11034, Invitrogen), Alexa Fluor 594 goat-anti-mouse (A11032 Invitrogen) and Alexa Fluor 594 streptavidin conjugates (S32356, Thermofisher) in a 1:250 dilution in 5% BSA in PBST.

Immunoprecipitation assays were performed using an immunoprecipitation kit (ab206996, Abcam) according to the manufacturer’s guidelines. HEK293T cells were grown in 100 mm dishes, irradiated with IR (10 Gy) prior to harvesting, and then returned to incubators at 37 °C for 1 h. Cell medium was aspirated, and cells were washed with ice-cold 1 x PBS. Cell dishes were placed on ice with 1 mL of RIPA Lysis Buffer (50 mM Tris-HCl pH 8.0, 150 mM NaCl, 1% IGEPAL CA-630, 0.5% sodium deoxycholate, 0.1% SDS) supplemented with freshly added protease inhibitor cocktail (2 μL/mL). Dishes were left on ice for 5 min to lyse cells, which were harvested directly into chilled microcentrifuge tubes. Cells were rotated end-over-end for 30 min at 4 °C. Cell lysates were cleared by centrifugation (10,000 x *g*, 10 min, 4 °C) and the supernatant was transferred to fresh chilled tubes. A specific amount of antibody (4 μg for DNA-PKcs; 10 μg for UFL1) against the target protein or an IgG-negative control antibody was added into low-binding microcentrifuge tubes. Cell lysates were diluted with 2 volumes of 50 mM Tris-HCl pH 8.0, 150 mM NaCl without detergents. Cell lysates and antibodies were mixed for 3-4 h or overnight at 4 °C on a rotary mixer. Buffer equilibrated Protein A/G Sepharose beads (25-40 μL) were added to each reaction and incubated for 1 h at 4 °C, rotating. Protein A/G Sepharose beads were pelleted and washed 3 times with 1 mL wash buffer and bound proteins were eluted in 40 μL 2 x SDS sample buffer (120 mM Tris-HCl pH 6.8, 4% SDS, 20% glycerol, 0.02% bromophenol blue, and 2.5% β-mercaptoethanol) by boiling at 95 °C for 5 min and analysed by immunoblotting.

#### Live-cell imaging and laser micro-irradiation

Live cell imaging was carried out on an Olympus (Evident) Xplore Spin system made up of an Olympus IX83 Confocal microscope body equipped with a Yokogawa CSU-W1 spinning disk. Images were captured with a 40 ×/1.4 oil objective and a Hamamatsu 2048 by 2048-pixel Orca fusion camera using cellSens software (Evident/Olympus) for image capturing. GFP fluorescence was excited with a 488 nm laser at 20% power, 1000 ms exposure. Throughout the experiment, the cells were maintained at 37 °C in HEPES-containing phenol red-free media (Gibco). Plasmids containing GFP-tagged XRCC4 (wildtype or Δ256-263) were transiently transfected in 35 mm imaging dishes with a coverslip thickness (1.5 H/170 µm) glass bottom that had been seeded with U2OS XRCC4 knock-out cells. Following transfection, cells were pre-sensitised in 10 μM BrdU for 24-48 h prior to laser micro-irradiation and imaging. Micro-irradiation was performed with a Rapp UGA-42 Caliburn laser ablation system. The 355 nm laser (with an output energy 200 µJ at 1.2 kHz and a pulse duration of 1.4 ns) was controlled by cellSens’ FRAP module in the software’s experiment manager. The laser path was created by drawing a straight-line region-of-interest (ROI) in cellSens. ROIs were drawn across the cell nuclei avoiding nucleoli, which were then ablated along the lines. The laser was set to 50%, and a 10% neutral density filter in the laser path reduced the power at the objective to 5%. 10 repetitions per ablation cycle were performed and imaging of the cells was performed in confocal mode before, during and after the ablation. The Olympus Xplore Spin system undergoes regular quality control for resolution and laser power, and a 16-point calibration was performed for the Rapp laser ablation system before imaging. GFP signal intensities at micro-irradiated sites were quantified using Fiji/ImageJ. For each time point, values were normalised to both background fluorescence (measured in an undamaged nuclear region) and the pre-damage baseline intensity at time point t=0. To reduce noise, a 15-second averaging window was applied, and the normalised intensity at t=0 was set to zero.

#### Crosslinking mass spectrometry (XL-MS)

20 µM UFM1 F35-BpF proteins were incubated with 20 µM UFM1 receptor proteins in PBS for 30 min at 4 °C in 0.2 ml thin-walled polypropylene PCR tubes (Greiner Bio-One). Samples were irradiated for 2 h with long-wave UV light (λ = 365 nm; Herolab UV-16 L, 8 W) at a 6 mm distance. Aliquots were collected before and after UV exposure for SDS-PAGE analysis. The crosslinked proteins were subjected to solution or in-gel tryptic digestion of bands excised from an SDS-PAGE gel, and analysed using a Vanquish Neo HPLC system coupled to an Orbitrap Lumos mass spectrometer via an EasySpray source (Thermo). Raw files were analysed directly using pLink2.0^99^ to identify crosslinked peptides. BpF was manually input as an amino acid (designated “X”) and the UV-inducible BpF crosslinker was manually input as crosslinking X to any other canonical amino acid with a crosslink or monolink mass of zero. Met oxidation, Gln/Asn deamidation, and N-terminal acetylation were included as variable modifications. Search parameters included a peptide length 3-100 and peptide mass of 400-10,000 Da, a maximum of 3 missed cleavages allowed, and a precursor and fragment mass tolerance of 20 ppm. An FDR of <1 % was used to filter out low-confidence crosslinks. Fasta files containing the amino acid sequences of the relevant protein constructs were used. For experiments with UFM1-UBA5, trypsin was combined with elastase digestion in tandem while for UFM1-XRCC4 experiments, trypsin was combined with chymotrypsin. The resulting high-confidence crosslinks were further manually inspected using pLabel to assess spectral evidence.

#### NMR experiments

Protein NMR spectra were recorded at 298 K on a Bruker 800 MHz spectrometer equipped with a ^1^H-^15^N TCI cryoprobe and z-axis gradients. Unless specified otherwise, samples were prepared in 10 mM HEPES, 150 mM NaCl, 50 mM L-arginine, 50 mM L-glutamate, 2 mM TCEP, and 1 mM EDTA at pH 6.8 for UFM1 and XRCC4^1-164^ or Bis-Tris pH 6.5, 150 mM KCl, 1 mM EDTA, 1 mM DTT for XRCC4^200-334^. ^1^H-^15^N spectra for XRCC4^1-164^ were acquired using the standard Bruker BEST-TROSY sequence with phase-sensitive Echo/Antiecho gradient selection. UFM1 ^1^H-^15^N and XRCC4^200-334 1^H-^15^N spectra were obtained using the standard Bruker sensitivity-enhanced, phase-sensitive HSQC sequence with Echo/Antiecho gradient selection. One-dimensional ^1^H spectra were recorded using excitation sculpting for water suppression. Spectral assignments for XRCC4 and UFM1 were based on BMRB entries 50742 (XRCC4^1-164^), 52398 (XRCC4^200-334^), and 26312, respectively. All data were processed and visualised using TopSpin 3.5 (Bruker), and protein:protein interactions were analysed using CCPN AnalysisAssign (v3.2.12). A list of plasmids used in this study is available in Supplementary Table S3.

#### Pulldown experiments

##### GFP pulldowns

HEK293T cells transfected with the indicated expression construct(s) were washed with ice-cold PBS and scraped into lysis buffer (50 mM Tris-HCl pH 7.5, 150 mM NaCl, 10% glycerol, 2 mM MgCl_2_, 10 mM *N*-ethylmaleimide) with 1 × cOmplete EDTA-free protease inhibitors (Roche) and 6 μL benzonase (Millipore), except for the GFP pulldowns shown in Figure 6E where no benzonase was used, and rotated at RT for 15 min. Subsequently, the lysates were centrifuged at 17,000 g for 10 min and the supernatant bound to 25 μL of GFP-Trap magnetic beads (Chromotek) for 4 h with end-over-end rotation at 4 °C. Protein-bound beads were then washed 5 times with lysis buffer containing 225 mM NaCl and resuspended in 2 × SDS Laemmli buffer (120 mM Tris-HCl pH 6.8, 4% SDS, 20% glycerol, 0.02% bromophenol blue and 2.5% β-mercatoethanol). 2% of input lysate were loaded unless stated otherwise.

##### GST pulldowns

For GST pulldown assays, HEK293T cell extracts were prepared as described for photo-crosslinking experiments. A total of 3-5 μg of GST-tagged proteins immobilised on glutathione magnetic beads (Promega) were incubated overnight at 4 °C with 600 μL of HEK293T cell extracts transiently transfected with the protein-of-interest, using end-over-end rotation. Beads were washed once with NETN buffer (50 mM Tris-HCl pH 8.0, 100 mM NaCl, 0.5 mM EDTA, 0.5% NP-40) and twice with PBS, then resuspended in 2 × SDS Laemmli buffer (120 mM Tris-HCl pH 6.8, 4% SDS, 20% glycerol, 0.02% bromophenol blue, 2.5% β-mercaptoethanol). Unless indicated otherwise, 4% of input lysate was loaded. Proteins were analysed by immunoblotting.

#### APEX2-based mass spectrometry screening

##### Biotin phenol (BP) synthesis^100^

The reagents and starting material were purchased from commercial sources and were carried forward without further purification. The reactions were performed under inert condition. Unless otherwise stated the reactions were monitored by thin layer chromatography. NMR was performed on 400 MHz or 500 MHz Bruker instruments, chemical shifts were reported in ppm, relative to solvents DMSO-d_6_ (δ=2.50 ppm for ^1^H NMR and δ=39.50 ppm for ^13^C NMR) or CDCl_3_ (δ=7.26 ppm for ^1^H NMR and δ=77.2 ppm for ^13^C NMR). Mass spectra were recorded using the electrospray technique. Mass spectra were obtained by the School of Chemistry Mass Spectrometry Service (University of Manchester) employing a Thermo Finnigan MAT95XP spectrometer*^100^*.

Biotin-NHS ester – to a solution of D-biotin (1.00 g, 4.09 mmol) in anhydrous DMF (12 mL), N-hydroxysuccinamide (NHS) ester (530 mg, 4.61 mmol) and 1-ethyl-3-(3-dimethylamipropyl) carbodiimide (565.8 mg, 4.91 mmol) were added and the reaction was stirred for 18 h at room temperature. After completion, DMF was removed under reduced pressure. The solid thus obtained was washed with cold ethanol to afford biotin-NHS ester (810 mg, 58%) as a white solid. **^1^H NMR** (400 MHz, DMSO-d_6_) δ 5.54 (d, *J* = 14.9 Hz, 2H), 3.43 (dd, *J* = 7.9, 5.1 Hz, 1H), 3.28 – 3.25 (m, 1H), 2.24 – 2.20 (m, 1H), 1.93 (d, *J* = 3.3 Hz, 5H), 1.78 (t, *J* = 7.4 Hz, 2H), 1.70 (t, *J* = 6.2 Hz, 1H), 0.80 – 0.73 (m, 3H), 0.64 – 0.46 (m, 3H). **^13^C NMR** (126 MHz, DMSO-d_6_) δ 170.3, 169.0, 162.8, 162.4, 61.1, 59.2, 55.3, 35.8, 30.8, 30.0, 27.9, 27.6, 25.5, 24.4. Spectral data matched those reported*^100^*.

Biotin phenol – a mixture of Biotin-NHS ester (124 mg, 0.36 mmol) and tyramine **2** (50.0 mg, 0.36 mmol) in DMF (12 mL) was prepared at room temperature. To this, Et_3_N (0.15 mL, 1.08 mmol) was added drop-wise and the reaction mixture was stirred for 18 h at ambient temperature. After completion, DMF was removed under reduced pressure, the crude mixture was purified using flash chromatography (90:10 DCM/MeOH) to yield biotin-phenol (91.6 mg, 70%) as a white solid. **^1^H NMR** (500 MHz, DMSO-d_6_) δ 9.22 (s, 1H), 7.80 (t, *J* = 5.6 Hz, 1H), 7.05 – 6.90 (m, 2H), 6.74 – 6.61 (m, 2H), 6.43 (s, 1H), 6.36 (s, 1H), 4.31 (dd, *J* = 7.8, 5.0 Hz, 1H), 4.12 (m, 1H), 3.24 – 3.13 (m, 2H), 3.10 – 3.06 (m, 1H), 2.82 (dd, *J* = 12.4, 5.1 Hz, 1H), 2.63 – 2.52 (m, 2H), 2.03 (t, *J* = 7.4 Hz, 2H), 1.67 – 1.54 (m, 1H), 1.54 – 1.37 (m, 3H), 1.27 (q, *J* = 7.0 Hz, 2H). **^13^C NMR** (126 MHz, DMSO-d_6_) δ 171.9, 162.8, 155.6, 129.5, 129.5 (2C), 115.1 (2C), 61.1, 59.2, 55.5, 40.4, 39.8, 35.2, 34.4, 28.2, 28.1, 25.3. Spectral data matched those reported*^100^*.

##### Construct design

Primers containing unique NheI and XhoI restriction sites were used to amplify UFC1 from testis cDNA by PCR and fused to APEX2 containing two N-terminal HA tags as well as two C-terminal nuclear localisation signals (NLS’s), amplified using primers containing XhoI and KpnI restriction sites. The APEX2 sequence was originally amplified from Addgene plasmid #49386, a gift from Alice Ting^101^.

##### Label-free proteomics

Samples were alkylated by addition of 5.5 mM chloroacetamide and loaded onto 4-12% gradient Bis-Tris gels. Proteins were separated by SDS-PAGE, stained using a Colloidal Blue Staining Kit (Life Technologies) and in-gel digested using trypsin. Peptides were extracted from gel and desalted on reversed-phase C18 StageTips^102^.

##### Mass spectrometry analysis

Peptides were analysed on a quadrupole Orbitrap mass spectrometer (Q Exactive Plus, Thermo Scientific) equipped with a UHPLC system (EASY-nLC 1200, Thermo Scientific) as described previously^103,104^. The mass spectrometer was operated in data-dependent mode, automatically switching between MS and MS2 acquisition. Survey full scan MS spectra (m/z 300-1,700) were acquired in the Orbitrap. The 15 most intense ions were sequentially isolated and fragmented by higher energy C-trap dissociation (HCD)^105^. An ion selection threshold of 5,000 was used. Peptides with unassigned charge states, as well as with charge states <+2, were excluded from fragmentation. Fragment spectra were acquired in the Orbitrap mass analyser.

##### Peptide identification

Raw data files were analysed using MaxQuant software (development version 1.5.2.8)^106^. Parent ion and MS2 spectra were searched against a database containing 98,566 human protein sequences obtained from UniProtKB (April 2018 release) using the Andromeda search engine^107^. Spectra were searched with a mass tolerance of 6 ppm in MS mode, 20 ppm in HCD MS2 mode, strict trypsin specificity and allowing up to two miscleavages. Cysteine carbamidomethylation was set as a fixed modification, whilst N-terminal acetylation and oxidation were set as variable modifications. The dataset was filtered based on posterior error probability (PEP) to arrive at a FDR<1% estimated using a target-decoy approach^108^.

##### Data processing

Processed data from MaxQuant were analysed in RStudio (R version 4.2.3). Proteins or peptides flagged as “reverse”, “only identified by site” or “potential contaminant” were excluded from downstream analysis. Only proteins identified by no less than two peptides and at least one unique peptide were used in downstream analysis. Statistical significance was assessed using the LIMMA package (limma-v 3.54.2)^109^. Gene ontology analyses were carried out for the most significantly enriched proteins (false discovery rate (FDR) ≤ 0.005) using ShinyGO 0.80^92^ and KEGG^93^.

##### SILAC labelling and neutravidin pulldowns

For SILAC labelling, U2OS cells were grown for two weeks prior to experiments in DMEM for SILAC (Thermofisher) containing either unlabelled amino acids (L-lysine, L-arginine), or amino acids labelled with stable isotopes (L-lysine [4,4,5,5-D4], L-arginine-U-13C6; L-lysine-U-13C6, ^15^N2, L-arginine-U-^13^C6, ^15^N4) purchased from Cambridge Isotope Laboratories. Media was supplemented with 4.5 g/L of glucose, 10% dialysed FBS (Sigma-Aldrich) and 1% L-glutamine/penicillin/streptomycin (v/v) (100 ×, Gibco). Cells transiently expressing UFC1 fused to APEX2 were irradiated with IR (10 Gy) and subsequently returned to the incubator for 30 min before adding 0.5 mM biotin phenol (BP) for 2 h prior to the start of the labelling reaction. The cells were washed with PBS and the labelling reaction initiated by adding 1 mM H_2_O_2_ for 2 min at room temperature. The reaction was terminated by washing cells thrice with a quencher solution containing 10 mM sodium azide, 10 mM sodium ascorbate, and 5 mM Trolox in PBS. Cells were lysed with ice-cold RIPA buffer (50 mM Tris-HCl pH 7.5, 150 mM NaCl, 0,1% SDS, 0,5% sodium deoxycholate, 1% Triton X-100 and freshly added protease inhibitor, 10 mM *N*-ethylmaleimide, 5 mM B-glycerophosphate, 5 mM NaF, 1 mM sodium orthovanadate). Cells were then sonicated at 30% power (QSonica Sonicators) for 50 s and centrifuged 15 min at 16,000 g at 4 °C. Protein concentrations were determined using Pierce 660 nm Protein Assay Reagent (Thermo Fisher Scientific). Reactions were performed in triplicate. Absorbance was measured at 660 nm using a plate reader (Varioskan Lux, Thermo Fisher Scientific) and protein concentrations in μg/μL were calculated using a standard curve. Lysates were mixed in 1:1:1 ratio and prepared for neutravidin pull-downs. Pierce High capacity NeutrAvidin agarose beads (29202, Thermofisher) were washed thrice with RIPA buffer and centrifuged for 1 min at 3,000 g. Beads were incubated with cell lysates (light, medium and heavy in 1:1:1) overnight at 4 °C. For elutions, the mixtures were centrifuged at 3000 g for 1 min at 4 °C. Beads were washed once with 1 mL RIPA buffer, once with 8 M urea containing 1% SDS, once with 1% SDS buffer and once with RIPA. Beads were carefully dried with a syringe to remove buffer after the last wash and then eluted by incubation for 15 min at 95 °C with 50 μL 3 × SDS loading buffer containing freshly added 1 mM DTT.

#### *In vitro* UFC1 charging assays

E1 (UBA5) (1 μM), UFM1 or UFM1 BpF-containing mutants (10 μM), and E2 (UFC1) (5 μM) were incubated in a buffer containing 50 mM Bis-Tris (pH 6.5), 100 mM NaCl and 10 mM MgCl_2_. Reactions were initiated by the addition of ATP (5 mM) and ran for a specified time at 30 °C. Before activation, an aliquot of the reaction mix was collected and taken as the control at time 0. The reactions were quenched with SDS sample buffer without β-mercaptoethanol, separated by 12% Bis-Tris non-reducing PAGE, and visualised by Coomassie brilliant blue staining.

#### *In vitro* UFMylation assays

For checking the function of UFM1 proteins and the UFM1* mutation, 1 µM His_6_-UBA5, 4 µM His_6_-UFC1, 0.25 µM UFL1-DDRGK1 (29-314) were mixed with 20 µM UFM1 protein (untagged or StrepII-tagged WT/F35* containing a free VG at the C-terminus) in 50 mM Tris pH 7.5, 10 mM MgCl_2_ and the reaction was started by addition of 10 mM ATP. The reaction was incubated for 1 h at 37 °C and quenched with 4x SDS sample buffer with reducing agent. Samples were loaded onto a 15% homemade SDS-PAGE gel and visualised with Coomassie Brilliant Blue or by immunoblotting after probing with the indicated antibodies.

For the UFMylation of DNA-PK, 0.5 µM Ku70/80 heterodimer was mixed 1:1 with Y-DNA ± 0.25 µM DNA-PKcs for 5 min at 25 °C to enable DNA-PK complex formation before the addition of 1 µM His_6_-UBA5, 4 µM His_6_-UFC1, 0.25 µM UFL1-DDRGK1, and 20 µM UFM1 in 50 mM Tris pH 7.5, 10 mM MgCl_2_ and the reaction was started by addition of 10 mM ATP. The reaction was incubated for 18 h (overnight) at 37 °C and quenched with 4 × SDS sample buffer with reducing agent. Samples were loaded onto 8% or 15% homemade SDS-PAGE gels and visualised with Coomassie Brilliant Blue or by immunoblotting after probing with the indicated antibodies.

For MS identification of the modified sites, the reaction was fully resolved by 15% SDS-PAGE with ± ATP samples from two repeats separated by empty lanes. Following brief staining with Coomassie Brilliant Blue, bands were excised at ∼70-100 kDa (for Ku70/80) and >100 kDa (for DNA-PKcs). Gel pieces were dehydrated and re-hydrated 5 times with HPLC-grade acetonitrile and HPLC-grade water, respectively, before drying to completeness in a vacuum centrifuge. Gel slices were rehydrated in 40 µl 50 mM TEAB (#catT7408, Sigma) and sequencing-grade trypsin was added at 20 ng/µl and incubated at RT for 20 min without shaking, before 100 µL 50 mM TEAB (#catT7408, Sigma) was added and incubated at 37 °C with shaking for 18 h. Overnight digests were acidified by the addition of formic acid to a final concentration of 0.2% (v/v) and dried to completeness in a vacuum centrifuge. Peptide digests were finally resuspended in 5 µL of 0.2% formic acid immediately prior to LC-MS/MS analysis. LC-MS/MS was carried out in DDA-mode using a Vanquish Neo HPLC system coupled to an Orbitrap Lumos mass spectrometer via an EasySpray source (Thermo). Data was analysed using Scaffold PTM v 4.0.2 (Proteome Software) with VG or DRVG as a variable modification on K residues to detect UFMylated sites.

#### Generation of stably inducible UFSP2 degron cells

The CRISPR-Cas9 plasmid utilised in this study was eSpCas9-hGeminin (Addgene plasmid #199344; RRID: Addgene_199344)^110^, a modified variant of eSpCas9(1.1)_No_FLAG_ATP1A1_G3_Dual_sgRNA (Addgene plasmid #86613; RRID: Addgene_86613), a gift from Yannick Doyon^111^, wherein the C-terminus of the eSpCas9 enzyme is fused to amino acids 1–110 of human Geminin. The hGeminin fusion protein facilitates degradation of eSpCas9 during G1 phase of the cell cycle, thereby preventing DSB formation in G1 phase, which would otherwise lead to indel formation via NHEJ. Consequently, Cas9 activity is restricted to the HR-favoured phases of the cell cycle (S/G)^112^. The eSpCas9-hGeminin plasmid also encodes the ATP1A1 G3 guide RNA (gRNA), enabling precise editing of the ATP1A1 gene, with an additional ATP1A1 template plasmid included to induce naturally occurring mutations in the ATP1A1 gene (encoding the sodium/potassium pump Na^+^/K ^+^ATPase). These mutations confer resistance to the drug ouabain (Tocris, #1076) without changing enzyme function^111^. The double degron sequence used comprises both dihydrofolate reductase (DHFR) and FK506-binding protein 12 (FKBP12). Previous studies have demonstrated that fusing DHFR and FKBP12 to a native protein at either the C- or N-terminus induces protein degradation upon removal of TMP (#T795615) and/or the addition of the degrader dtagV1 (Tocris, #6914), respectively^113,114^.

*UFSP2* gRNAs were designed using CRISPOR software (http://crispor.tefor.net/crispor.py) with 20 bp guides within 100 bp of the target region selected. The following guides were generated to target the 3’-end of the gene: gRNA1: 5’-TAAATCATATTTGGTCGCTG-3’ and gRNA2: 5’-AGTCAAAGACTGCAGTAGAG-3’. Guide RNAs were synthesised using GeneArt (Invitrogen) to be expressed using a tRNA and H1 promoter, respectively, and cloned into the eSpCas9-hGeminin plasmid. Correct insertion of gRNAs was validated by Sanger sequencing with the primer 5’-ACCGTAAATACTCCACCCAT-3’. The homology donor template to insert the DHFR-FKBP12 coding sequences into the 3’-end of *UFSP2* were designed using Benchling, including silent mutations in the PAM and gRNA targets to reduce re-editing, and similarly synthesised using GeneArt (Invitrogen). The template was inserted into a plasmid backbone and verified by Sanger Sequencing using the forward primer 5’-GTAAAACGACGGCCAGT-3’ and the reverse primer 5’-AGGAAACAGCTATGACC-3’. Successfully verified plasmids for guides and template were subsequently amplified using a Maxiprep Kit (Qiagen, #12163) for transfection.

For transfection, U2OS cells were harvested by trypsinisation at 70-90% confluence. 5 × 10^6^ cells were co-transfected with 3 plasmids: eSpCas9-hGeminin containing the gRNAs, ATP1A1 template and repair template for double-degron insertion, using 10 µL of each all at 1 µg/µL. Cells were electroporated using the Neon Transfection System (Invitrogen, #MPK5000) at 1400 V, 30 ms, 1 pulse. Electroporated cells were incubated in 150 mm round dishes for 48-72 h, then treated with ouabain selection media (10 µM) which was replaced every few days for 5-7 days. Colonies were picked under the microscope and placed in 96-well plates, then expanded up to 6-well plates for cryopreservation and DNA and/or protein analysis.

For screening successful insertion of the double-degron tag at the *UFSP2* locus, cell pellets were lysed in PCR direct lysis reagent (302-C, Viagen Biotech) plus Proteinase K (BIP4206, Apollo Scientific) for 4 h at 55 °C then 95 °C for 1 h. 1 µl of this mixture was used to amplify the target locus by PCR with the primers: F: 5’-AATAACTCAGTGGCCCTAGGAG-3’ and R: 5’-CTGAAGATCTCGCTTGGTCATG-3’ using HotStartTaq Master Mix (203443, Qiagen) using the following conditions: 95 °C for 15 min (to activate the HotStartTaq); 35 cycles of 95 °C 1 min, 52 °C 45 s, 72 °C 2.5 min; 72 °C 10 min. Gel-purified PCR products were sequenced by Sanger sequencing using the primers: F-seq: 5’-CGTCTTCTCTATCCAGGGCTG-3’ and R: 5’-CAAACTAGAACCAGGATTTAATGTGA-3’.

#### Chromatin fractionation following DNA double-strand break induction

Chromatin fractionation experiments broadly followed a previously established method^80,81^. Stock solution of calicheamicin γ1 (Cali, CAS: 108212-75-5, #HY-19609, MedChem Express) was made at 250 μM in DMSO and stored at −70 °C. For drug exposure, exponentially growing cells in 60 mm or 100 mm (U2OS and HEK293T cells) or 150 mm (fibroblasts) diameter dishes were either mock-treated (DMSO) or treated with 5 nM Cali in fresh medium for 1 h at 37 °C. In chase experiments, cells were washed with fresh media and incubated in fresh media without Cali at 37 °C for the indicated time points. Cells were washed with PBS and harvested using trypsin-EDTA followed by two washes with ice-cold PBS. For chromatin fractionation, cells were first resuspended for 7 min on ice in 100 µL of extraction buffer I (50 mM Hepes-KOH pH 7.5, 150 mM NaCl, 1 mM EDTA, 0.1% Triton X-100, 1 × EDTA-free cOmplete protease inhibitors, 1 × protein phosphatase inhibitors) with intermittent gentle vortexing. Following centrifugation at 17,000 x *g* for 3 min, the supernatant was removed and stored on ice, and pellets were gently resuspended with pipette tips in 100 µL of extraction buffer II (50 mM Hepes-KOH pH 7.5, 75 mM NaCl, 1 mM EDTA, 0.025% Triton X-100, 30 U RNAseA/T1 (NEB)), incubated for 15 min at 25 °C under agitation and centrifuged as above. The supernatants from both centrifugation steps were pooled (soluble protein fraction). Pellets (chromatin fractions) were resuspended in 100 µL extraction buffer III (50 mM Tris-HCl pH 8.1, 10 mM EDTA, 0.1% Triton X-100, 0.3% SDS) and sonicated for 20 s at 30% amplitude in 10 s pulses (QSonica Q125 sonicator with a 2 mm diameter probe). Protein content was measured with BCA reagent (Pierce). Immunoblotting was performed as described. For quantification of chromatin-bound NHEJ factors, bands were quantified in Image Lab (Bio-Rad) or Image J and normalised to the β-actin loading control from the same blot after subtracting background blot intensity. Amounts were normalised to the “wildtype” or “normal” condition within the same antibody blot.

## REFERENCES

1. Blackford, A. N. & Jackson, S. P. ATM, ATR, and DNA-PK: The Trinity at the Heart of the DNA Damage Response. Mol Cell 66, 801–817 (2017).

2. Jackson, S. P. & Bartek, J. The DNA-damage response in human biology and disease. Nature 461, 1071–1078 (2009).

3. Ciccia, A. & Elledge, S. J. The DNA Damage Response: Making It Safe to Play with Knives. Mol Cell 40, 179–204 (2010).

4. Scully, R., Panday, A., Elango, R. & Willis, N. A. DNA double-strand break repair-pathway choice in somatic mammalian cells. Nat Rev Mol Cell Biol 20, 698–714 (2019).

5. Blackford, A. N. & Stucki, M. How Cells Respond to DNA Breaks in Mitosis. Trends Biochem Sci 45, 321–331 (2020).

6. Chang, H. H. Y., Pannunzio, N. R., Adachi, N. & Lieber, M. R. Non-homologous DNA end joining and alternative pathways to double-strand break repair. Nat Rev Mol Cell Biol 18, 495–506 (2017).

7. Schwertman, P., Bekker-Jensen, S. & Mailand, N. Regulation of DNA double-strand break repair by ubiquitin and ubiquitin-like modifiers. Nat Rev Mol Cell Biol 17, 379– 394 (2016).

8. Garvin, A. J. & Morris, J. R. SUMO, a small, but powerful, regulator of double-strand break repair. Philosophical Transactions of the Royal Society B: Biological Sciences 372, 20160281 (2017).

9. Bhachoo, J. S. & Garvin, A. J. SUMO and the DNA damage response. Biochem Soc Trans 52, 773–792 (2024).

10. Brown, J. S. & Jackson, S. P. Ubiquitylation, neddylation and the DNA damage response. Open Biol 5, 150018 (2015).

11. Cappadocia, L. & Lima, C. D. Ubiquitin-like Protein Conjugation: Structures, Chemistry, and Mechanism. Chem Rev 118, 889–918 (2018).

12. Da Costa, I. C. & Schmidt, C. K. Ubiquitin-like proteins in the DNA damage response: the next generation. Essays Biochem 64, 737–752 (2020).

13. Sandy, Z., da Costa, I. C. & Schmidt, C. K. More than Meets the ISG15: Emerging Roles in the DNA Damage Response and Beyond. Biomolecules 10, 1557 (2020).

14. Komatsu, M., Inada, T. & Noda, N. N. The UFM1 system: Working principles, cellular functions, and pathophysiology. Mol Cell 84, 156–169 (2024).

15. Zhao, B., Rothenberg, E., Ramsden, D. A. & Lieber, M. R. The molecular basis and disease relevance of non-homologous DNA end joining. Nat Rev Mol Cell Biol 21, 765– 781 (2020).

16. Certo, M. T. et al. Tracking genome engineering outcome at individual DNA breakpoints. Nat Methods 8, 671–676 (2011).

17. Schmidt, C. K. et al. Systematic E2 screening reveals a UBE2D–RNF138–CtIP axis promoting DNA repair. Nat Cell Biol 17, 1458–1470 (2015).

18. Morris, J. R. et al. The SUMO modification pathway is involved in the BRCA1 response to genotoxic stress. Nature 462, 886–890 (2009).

19. Galanty, Y. et al. Mammalian SUMO E3-ligases PIAS1 and PIAS4 promote responses to DNA double-strand breaks. Nature 462, 935–939 (2009).

20. Morris, J. R. & Garvin, A. J. SUMO in the DNA Double-Stranded Break Response: Similarities, Differences, and Cooperation with Ubiquitin. J Mol Biol 429, 3376–3387 (2017).

21. Garvin, A. J. et al. SUMO4 promotes SUMO deconjugation required for DNA double-strand-break repair. Mol Cell 85, 877–893.e9 (2025).

22. Oka, Y., Bekker-Jensen, S. & Mailand, N. Ubiquitin-like protein UBL5 promotes the functional integrity of the Fanconi anemia pathway. EMBO J 34, 1385–1398 (2015).

23. Oka, Y. et al. UBL5 is essential for pre-mRNA splicing and sister chromatid cohesion in human cells. EMBO Rep 15, 956–964 (2014).

24. Panichnantakul, P. et al. Protein UFMylation regulates early events during ribosomal DNA-damage response. Cell Rep 43, 114738 (2024).

25. Wang, Z. et al. MRE11 UFMylation promotes ATM activation. Nucleic Acids Res 47, 4124–4135 (2019).

26. Qin, B. et al. STK38 promotes ATM activation by acting as a reader of histone H4 ufmylation. Sci Adv 6, (2020).

27. Qin, B. et al. UFL1 promotes histone H4 ufmylation and ATM activation. Nat Commun 10, 1242 (2019).

28. Gong, Y. et al. PARP1 UFMylation ensures the stability of stalled replication forks. Proceedings of the National Academy of Sciences 121, (2024).

29. Tan, Q. & Xu, X. PTIP UFMylation promotes replication fork degradation in BRCA1-deficient cells. Journal of Biological Chemistry 300, 107312 (2024).

30. Tian, T. et al. UFL1 triggers replication fork degradation by MRE11 in BRCA1/2-deficient cells. Nat Chem Biol 20, 1650–1661 (2024).

31. Raso, M. C. et al. Interferon-stimulated gene 15 accelerates replication fork progression inducing chromosomal breakage. Journal of Cell Biology 219, (2020).

32. Wardlaw, C. P. & Petrini, J. H. J. ISG15 conjugation to proteins on nascent DNA mitigates DNA replication stress. Nat Commun 13, 5971 (2022).

33. Moro, R. N. et al. Interferon restores replication fork stability and cell viability in BRCA-defective cells via ISG15. Nat Commun 14, 6140 (2023).

34. Wang, Y. et al. Autophagy Regulates Chromatin Ubiquitination in DNA Damage Response through Elimination of SQSTM1/p62. Mol Cell 63, 34–48 (2016).

35. Xu, F. et al. Autophagy Promotes the Repair of Radiation-Induced DNA Damage in Bone Marrow Hematopoietic Cells via Enhanced STAT3 Signaling. Radiat Res 187, 382 (2017).

36. Ge, R. et al. XPA promotes autophagy to facilitate cisplatin resistance in melanoma cells through the activation of PARP1. Journal of Investigative Dermatology 136, S115 (2016).

37. Krenciute, G. et al. Nuclear BAG6-UBL4A-GET4 Complex Mediates DNA Damage Signaling and Cell Death. Journal of Biological Chemistry 288, 20547–20557 (2013).

38. Tatsumi, K. et al. A Novel Type of E3 Ligase for the Ufm1 Conjugation System. Journal of Biological Chemistry 285, 5417–5427 (2010).

39. Cai, Y. et al. UFBP1, a Key Component of the Ufm1 Conjugation System, Is Essential for Ufmylation-Mediated Regulation of Erythroid Development. PLoS Genet 11, e1005643 (2015).

40. Peter, J. J. et al. A non-canonical scaffold-type E3 ligase complex mediates protein UFMylation. EMBO J 41, e111015 (2022).

41. Ishimura, R. et al. The UFM1 system regulates ER-phagy through the ufmylation of CYB5R3. Nat Commun 13, 7857 (2022).

42. Habisov, S. et al. Structural and Functional Analysis of a Novel Interaction Motif within UFM1-activating Enzyme 5 (UBA5) Required for Binding to Ubiquitin-like Proteins and Ufmylation. Journal of Biological Chemistry 291, 9025–9041 (2016).

43. Brüninghoff, K. et al. Identification of SUMO Binding Proteins Enriched after Covalent Photo-Cross-Linking. ACS Chem Biol 15, 2406–2414 (2020).

44. Chin, J. W., Martin, A. B., King, D. S., Wang, L. & Schultz, P. G. Addition of a photocrosslinking amino acid to the genetic code of Escherichia coli. Proceedings of the National Academy of Sciences 99, 11020–11024 (2002).

45. Taupitz, K. F., Dörner, W. & Mootz, H. D. Covalent Capturing of Transient SUMO–SIM Interactions Using Unnatural Amino Acid Mutagenesis and Photocrosslinking. Chemistry – A European Journal 23, 5978–5982 (2017).

46. Brüninghoff, K., Wulff, S., Dörner, W., Geiss-Friedlander, R. & Mootz, H. D. A Photo-Crosslinking Approach to Identify Class II SUMO-1 Binders. Front Chem 10, (2022).

47. Sandy, Z. et al. Site-selective photo-crosslinking for the characterisation of transient ubiquitin-like protein-protein interactions. PLoS One 20, e0316321 (2025).

48. Kaur, K., Park, H., Pandey, N., Azuma, Y. & De Guzman, R. N. Identification of a new small ubiquitin-like modifier (SUMO)-interacting motif in the E3 ligase PIASy. Journal of Biological Chemistry 292, 10230–10238 (2017).

49. Zhang, M. et al. RCAD/Ufl1, a Ufm1 E3 ligase, is essential for hematopoietic stem cell function and murine hematopoiesis. Cell Death Differ 22, 1922–1934 (2015).

50. HU, X., et al. Ubiquitin-fold modifier 1 inhibits apoptosis by suppressing the endoplasmic reticulum stress response in Raw264.7 cells. Int J Mol Med 33, 1539– 1546 (2014).

51. Lemaire, K. et al. Ubiquitin Fold Modifier 1 (UFM1) and Its Target UFBP1 Protect Pancreatic Beta Cells from ER Stress-Induced Apoptosis. PLoS One 6, e18517 (2011).

52. Zhang, X. et al. Ufmylation regulates granulosa cell apoptosis via ER stress but not oxidative stress during goat follicular atresia. Theriogenology 169, 47–55 (2021).

53. Liu, J. et al. UFMylation maintains tumour suppressor p53 stability by antagonizing its ubiquitination. Nat Cell Biol 22, 1056–1063 (2020).

54. Li, G. et al. Eg5 UFMylation promotes spindle organization during mitosis. Cell Death Dis 15, 544 (2024).

55. Yu, L. et al. The UFM1 cascade times mitosis entry associated with microcephaly. The FASEB Journal 34, 1319–1330 (2020).

56. Gak, I. A., et al. UFMylation regulates translational homeostasis and cell cycle progression. bioRxiv (2020).

57. Liu, Y. et al. UFMylation maintains YAP stability to promote vascular endothelial cell senescence. iScience 28, 111854 (2025).

58. Balce, D. R. et al. UFMylation inhibits the proinflammatory capacity of interferon-γ– activated macrophages. Proceedings of the National Academy of Sciences 118, (2021).

59. Blazev, R. et al. Site-specific quantification of the in vivo UFMylome reveals myosin modification in ALS. Cell Reports Methods 5, 101048 (2025).

60. Dynan, W. S. & Yoo, S. Interaction of Ku protein and DNA-dependent protein kinase catalytic subunit with nucleic acids. Nucleic Acids Res 26, 1551–1559 (1998).

61. Chaplin, A. K. et al. Cryo-EM of NHEJ supercomplexes provides insights into DNA repair. Mol Cell 81, 3400–3409.e3 (2021).

62. Chen, S. et al. Structural basis of long-range to short-range synaptic transition in NHEJ. Nature 593, 294–298 (2021).

63. Cabello-Lobato, M. J., Schmidt, C. K. & Cliff, M. J. 1H, 13C, 15N backbone resonance assignment for the 1–164 construct of human XRCC4. Biomol NMR Assign 15, 389–395 (2021).

64. Cabello-Lobato, M. J. et al. Microarray screening reveals two non-conventional SUMO-binding modules linked to DNA repair by non-homologous end-joining. Nucleic Acids Res 50, 4732–4754 (2022).

65. Andres, S. N. et al. A human XRCC4–XLF complex bridges DNA. Nucleic Acids Res 40, 1868–1878 (2012).

66. Hung, V. et al. Spatially resolved proteomic mapping in living cells with the engineered peroxidase APEX2. Nat Protoc 11, 456–475 (2016).

67. Komatsu, M. et al. A novel protein-conjugating system for Ufm1, a ubiquitin-fold modifier. EMBO J 23, 1977–1986 (2004).

68. Osborne, H. C., Irving, E., Forment, J. V. & Schmidt, C. K. E2 enzymes in genome stability: pulling the strings behind the scenes. Trends Cell Biol 31, 628–643 (2021).

69. Liang, J. R. et al. A Genome-wide ER-phagy Screen Highlights Key Roles of Mitochondrial Metabolism and ER-Resident UFMylation. Cell 180, 1160–1177.e20 (2020).

70. Yiu, S. P. T. et al. An Epstein-Barr virus protein interaction map reveals NLRP3 inflammasome evasion via MAVS UFMylation. Mol Cell 83, 2367–2386.e15 (2023).

71. Snider, D. L., Park, M., Murphy, K. A., Beachboard, D. C. & Horner, S. M. Signaling from the RNA sensor RIG-I is regulated by ufmylation. Proceedings of the National Academy of Sciences 119, (2022).

72. Mao, M. et al. Modification of PLAC8 by UFM1 affects tumorous proliferation and immune response by impacting PD-L1 levels in triple-negative breast cancer. J Immunother Cancer 10, e005668 (2022).

73. Zhu, J. et al. P4HB UFMylation regulates mitochondrial function and oxidative stress. Free Radic Biol Med 188, 277–286 (2022).

74. Yang, J. et al. Metformin induces Ferroptosis by inhibiting UFMylation of SLC7A11 in breast cancer. Journal of Experimental & Clinical Cancer Research 40, 206 (2021).

75. Li, J. et al. Ufm1-Specific Ligase Ufl1 Regulates Endoplasmic Reticulum Homeostasis and Protects Against Heart Failure. Circ Heart Fail 11, (2018).

76. Sutherland, J. D. & Barrio, R. Putting the Stress on UFM1 (Ubiquitin-Fold Modifier 1). Circ Heart Fail 11, (2018).

77. Molendijk, J. et al. Proteome-wide systems genetics identifies UFMylation as a regulator of skeletal muscle function. Elife 11, (2022).

78. Peter, J. J. et al. A non-canonical scaffold-type E3 ligase complex mediates protein UFMylation. EMBO J 41, (2022).

79. Seif-El-Dahan, M. et al. PAXX binding to the NHEJ machinery explains functional redundancy with XLF. Sci Adv 9, (2023).

80. Britton, S. et al. DNA damage triggers SAF-A and RNA biogenesis factors exclusion from chromatin coupled to R-loops removal. Nucleic Acids Res 42, 9047–9062 (2014).

81. Elmroth, K. Cleavage of cellular DNA by calicheamicin γ1. DNA Repair (Amst) 2, 363– 374 (2003).

82. Karczewski, K. J. et al. The mutational constraint spectrum quantified from variation in 141,456 humans. Nature 581, 434–443 (2020).

83. Muona, M. et al. Biallelic Variants in UBA5 Link Dysfunctional UFM1 Ubiquitin-like Modifier Pathway to Severe Infantile-Onset Encephalopathy. The American Journal of Human Genetics 99, 683–694 (2016).

84. Kumar, M. et al. Structural basis for UFM1 transfer from UBA5 to UFC1. Nat Commun 12, 5708 (2021).

85. Ishimura, R. et al. The UFM1 system regulates ER-phagy through the ufmylation of CYB5R3. Nat Commun 13, 7857 (2022).

86. Matunis, M. J., Zhang, X.-D. & Ellis, N. A. SUMO: The Glue that Binds. Dev Cell 11, 596–597 (2006).

87. Vu, D.-D. et al. Multivalent interactions of the disordered regions of XLF and XRCC4 foster robust cellular NHEJ and drive the formation of ligation-boosting condensates in vitro. Nat Struct Mol Biol 31, 1732–1744 (2024).

88. Bossaert, M. et al. Identification of the main barriers to Ku accumulation in chromatin. Cell Rep 43, 114538 (2024).

89. Lee, L. et al. UFMylation of MRE11 is essential for telomere length maintenance and hematopoietic stem cell survival. Sci Adv 7, (2021).

90. Chung, C. H. & Yoo, H. M. Emerging role of protein modification by UFM1 in cancer. Biochem Biophys Res Commun 633, 61–63 (2022).

91. Krzywinski, M. et al. Circos: An information aesthetic for comparative genomics. Genome Res 19, 1639–1645 (2009).

92. Ge, S. X., Jung, D. & Yao, R. ShinyGO: a graphical gene-set enrichment tool for animals and plants. Bioinformatics 36, 2628–2629 (2020).

93. Kanehisa, M., Furumichi, M., Sato, Y., Ishiguro-Watanabe, M. & Tanabe, M. KEGG: integrating viruses and cellular organisms. Nucleic Acids Res 49, D545–D551 (2021).

94. Chaplin, A. K. et al. Dimers of DNA-PK create a stage for DNA double-strand break repair. Nat Struct Mol Biol 28, 13–19 (2021).

95. Wang, J. L. et al. Dissection of DNA double-strand-break repair using novel single-molecule forceps. Nat Struct Mol Biol 25, 482–487 (2018).

96. Sibanda, B. L. et al. Crystal structure of an Xrcc4-DNA ligase IV complex. Nat Struct Biol 8, 1015–1019 (2001).

97. Wu, Q. et al. Non-homologous end-joining partners in a helical dance: structural studies of XLF–XRCC4 interactions. Biochem Soc Trans 39, 1387–1392 (2011).

98. Li, Y. et al. Crystal structure of human XLF/Cernunnos reveals unexpected differences from XRCC4 with implications for NHEJ. EMBO J 27, 290–300 (2008).

99. Chen, Z.-L. et al. A high-speed search engine pLink 2 with systematic evaluation for proteome-scale identification of cross-linked peptides. Nat Commun 10, 3404 (2019).

100. Zhou, Y. et al. Expanding APEX2 Substrates for Proximity-Dependent Labeling of Nucleic Acids and Proteins in Living Cells. Angewandte Chemie International Edition 58, 11763–11767 (2019).

101. Lam, S. S. et al. Directed evolution of APEX2 for electron microscopy and proximity labeling. Nat Methods 12, 51–54 (2015).

102. Rappsilber, J., Mann, M. & Ishihama, Y. Protocol for micro-purification, enrichment, pre-fractionation and storage of peptides for proteomics using StageTips. Nat Protoc 2, 1896–1906 (2007).

103. Kelstrup, C. D., Young, C., Lavallee, R., Nielsen, M. L. & Olsen, J. V. Optimized Fast and Sensitive Acquisition Methods for Shotgun Proteomics on a Quadrupole Orbitrap Mass Spectrometer. J Proteome Res 11, 3487–3497 (2012).

104. Bekker-Jensen, D. B. et al. A Compact Quadrupole-Orbitrap Mass Spectrometer with FAIMS Interface Improves Proteome Coverage in Short LC Gradients. Molecular & Cellular Proteomics 19, 716–729 (2020).

105. Olsen, J. V et al. Higher-energy C-trap dissociation for peptide modification analysis. Nat Methods 4, 709–712 (2007).

106. Cox, J. & Mann, M. MaxQuant enables high peptide identification rates, individualized p.p.b.-range mass accuracies and proteome-wide protein quantification. Nat Biotechnol 26, 1367–1372 (2008).

107. Cox, J. et al. Andromeda: A Peptide Search Engine Integrated into the MaxQuant Environment. J Proteome Res 10, 1794–1805 (2011).

108. Elias, J. E. & Gygi, S. P. Target-decoy search strategy for increased confidence in large-scale protein identifications by mass spectrometry. Nat Methods 4, 207–214 (2007).

109. Smyth, G. K. Linear Models and Empirical Bayes Methods for Assessing Differential Expression in Microarray Experiments. Stat Appl Genet Mol Biol 3, 1–25 (2004).

110. Morrison, K. R. et al. Elevated basal AMP-activated protein kinase activity sensitizes colorectal cancer cells to growth inhibition by metformin. Open Biol 13, (2023).

111. Agudelo, D. et al. Marker-free coselection for CRISPR-driven genome editing in human cells. Nat Methods 14, 615–620 (2017).

112. Gutschner, T., Haemmerle, M., Genovese, G., Draetta, G. F. & Chin, L. Post-translational Regulation of Cas9 during G1 Enhances Homology-Directed Repair. Cell Rep 14, 1555–1566 (2016).

113. Iwamoto, M., Björklund, T., Lundberg, C., Kirik, D. & Wandless, T. J. A General Chemical Method to Regulate Protein Stability in the Mammalian Central Nervous System. Chem Biol 17, 981–988 (2010).

114. de Azevedo, M. F. et al. Systematic Analysis of FKBP Inducible Degradation Domain Tagging Strategies for the Human Malaria Parasite Plasmodium falciparum. PLoS One 7, e40981 (2012).

