## Supplementary material for "UFMylation orchestrates chromatin engagement of core NHEJ components to promote DNA double-strand break repair": Text and Figures

### Supplemental Figure S1

**A** **Module 1 results - TLR**

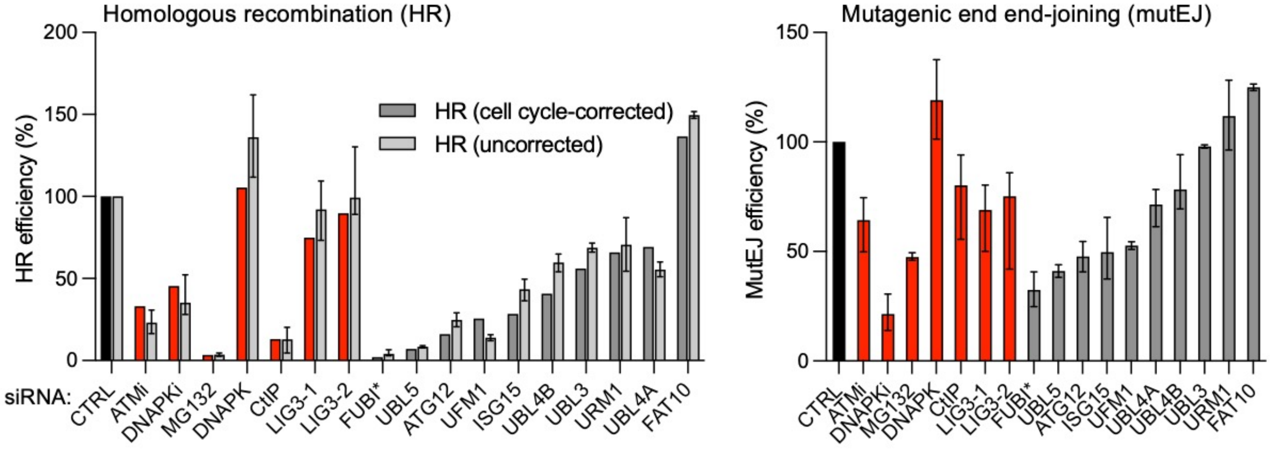


**B** **Module 2 results – IRIF kinetics**

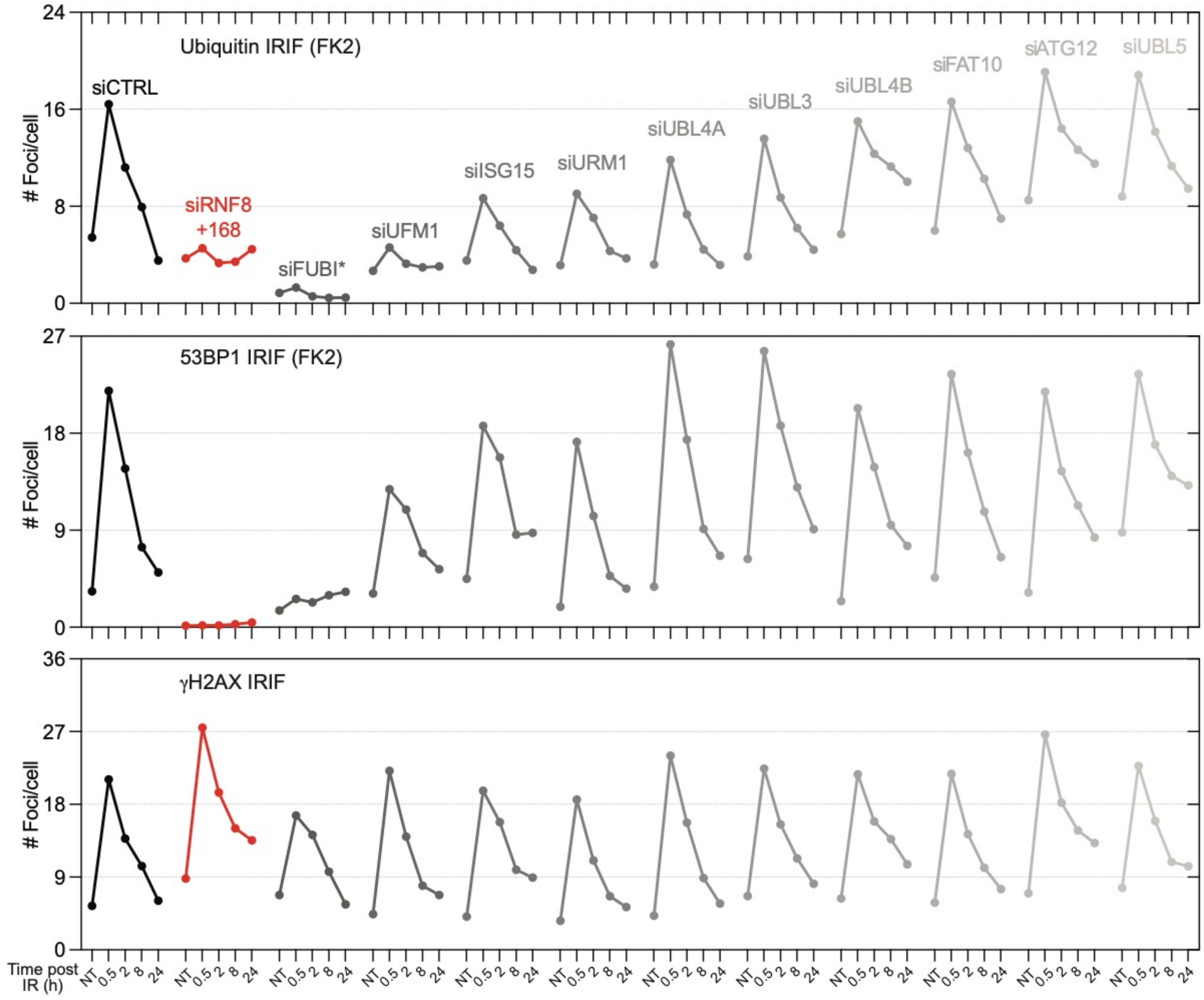

**Figure S1**. **Individual readouts for two-module screening of UBLs for roles in the DNA damage response.** (**A**) Left: Module 1 screening results for homologous recombination (HR) efficiency, using the traffic light reporter (TLR) system integrated into U2OS cells. Given HR's limitation to S/G2, results are presented corrected to flow-cytometry S/G2 values (left bars) in comparison to uncorrected values (right bars). Non-targeting control (CTRL) siRNA targeting luciferase (black) and small molecule inhibitors targeting ATM, DNA-PK and the proteasome (MG132) as well as siRNAs targeting DNAP-PK, CtIP and LIG3 (red) were used as negative and positive controls, respectively. Control conditions were published previously, due to forming part of the same screening pipeline[17]. Data represent means ± ranges of n=9 (siCTRL), n=8 (siCtIP), n=6 (siLIG3-1), n=5 (DNAPKi, siDNA-PK, siLIG3-2), n=4 (ATMi), n=3 (MG132, siFUBI, siISG15, siUBL4A, siUBL4B, siURM1) or n=2 (siUBL5, siATG12, siUFM1, siUBL3, siFAT10) biological replicates. Right: Module 1 screening results for DNA double-strand break repair by mutagenic end-joining (mutEJ), using the traffic light reporter (TLR system) integrated into U2OS cells. Controls are as described in (A). Data represent means ± ranges of n=9 (siCTRL), n=8 (siCtIP), n=6 (siLIG3-1), n=5 (DNAPKi, siDNA-PK, siLIG3-2), n=4 (ATMi), n=3 (MG132, siFUBI, siISG15, siUBL4A, siUBL4B, siURM1) or n=2 (siUBL5, siATG12, siUFM1, siUBL3, siFAT10) biological replicates. (**B**) Recruitment and resolution kinetics of conjugated ubiquitin recognised by the FK2 antibody, 53BP1 and γH2AX ionising radiation-induced foci (IRIF) in U2OS cells at the indicated time points after IR treatment (2 Gy), compared to the non-treated condition. Non-targeting control siRNA (siCTRL, against luciferase) and an siRNA mix targeting RNF8 plus RNF168 were used as negative and positive controls, respectively. Control data were published previously, due to forming part of the same screening pipeline[17]. Plots represent means ± range for n=9 (siCTRL and siRNF8+168) or n=2 (UBL siRNAs) 96-plate wells integrating, on average, >1000 cells imaged per UBL, and >8000 per control siRNA, per condition, with >400,000 cells imaged in total in one screening experiment.

Abbreviations: CTRL: control; HR: homologous recombination; IRIF: ionising radiation-induced foci; mutEJ: mutagenic end-joining; NT: non-treated; TLR: traffic light reporter.

*Follow-up analyses indicated that the effects observed for FUBI were likely attributable to off-target activity, potentially due to sequence overlap between one of the siRNAs and UBE2I (also known as UBC9), the E2 conjugating enzyme for SUMO. Given SUMO's well-established role in DSB repair[18–21], FUBI was excluded from further analysis.

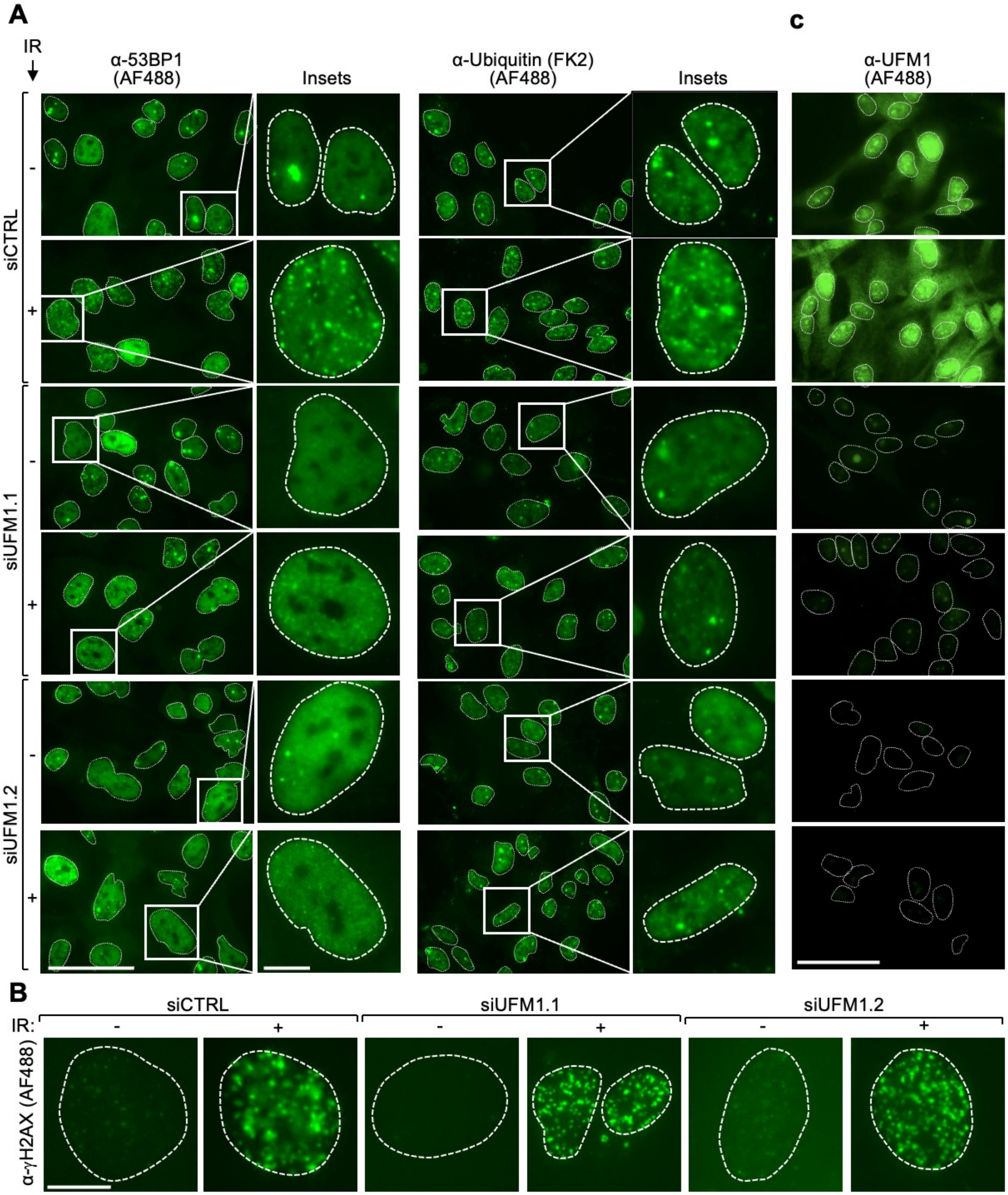


**Figure S2. UFM1 promotes recruitment of 53BP1 and conjugated ubiquitin to DNA damage sites.** Formation of ionising radiation-induced foci (IRIF) of 53BP1 (left) and ubiquitin (FK2 antibody, right) (**A**) as well as γH2AX (**B**) in siUFM1-depleted U2OS cells, using two independent siRNAs (siUFM1.1, siUFM1.2), compared to non-targeting control siRNA (siCTRL, against luciferase). (**C**) siRNA depletion efficiencies of two independent siRNAs targeting UFM1 (siUFM1.1, siUFM1.2), used in (A), as illustrated by diminished immunofluorescent staining in U2OS cells, using an α-UFM1 antibody. Dashed white lines mark nuclei outlines according to DAPI staining. Scale bar represents 50 µm for scaled-out images and 10 µm for insets in (A), 10 µm in (B), and 50 µm in (C).

Abbreviations: α: anti; AF: Alexa Fluor; IRIF: ionising radiation-induced foci.

#### Supplemental Figure S3

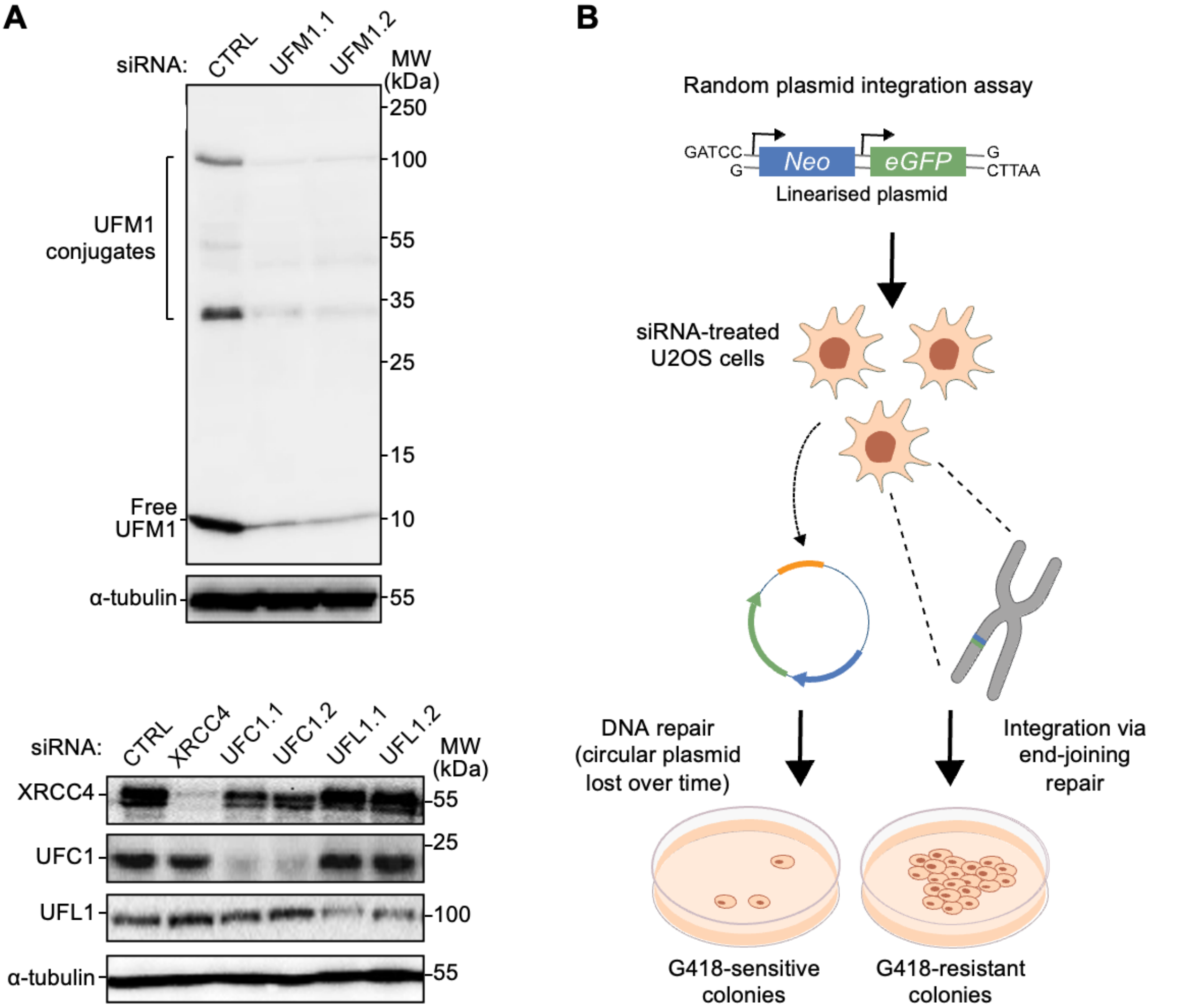


**Figure S3. Random plasmid integration information for evaluating UFM1 pathway components in end-joining repair.** (**A**) siRNA depletion efficiencies assessed by immunoblotting of two independent siRNAs targeting UFM1 (UFM1.1, UFM1.2), UFC1 (UFC1.1, UFC1.2) and UFL1 (UFL1.1, UFL1.2), as well as a positive control, siXRCC4. (**B**) Schematic overview of random plasmid integration assay. Cells were transfected with a linearised plasmid harbouring a neomycin resistance cassette (*Neo*) conferring resistance to geneticin (G418), along with a *GFP* reporter to assess transfection efficiency. Successful G418-resistant colony formation requires genomic integration of the *Neo* cassette via end-joining repair, enabling quantification of surviving colonies as a functional readout of end-joining repair efficiency.

Abbreviations: α: anti; MW: molecular weight; Neo: neomycin resistance cassette.

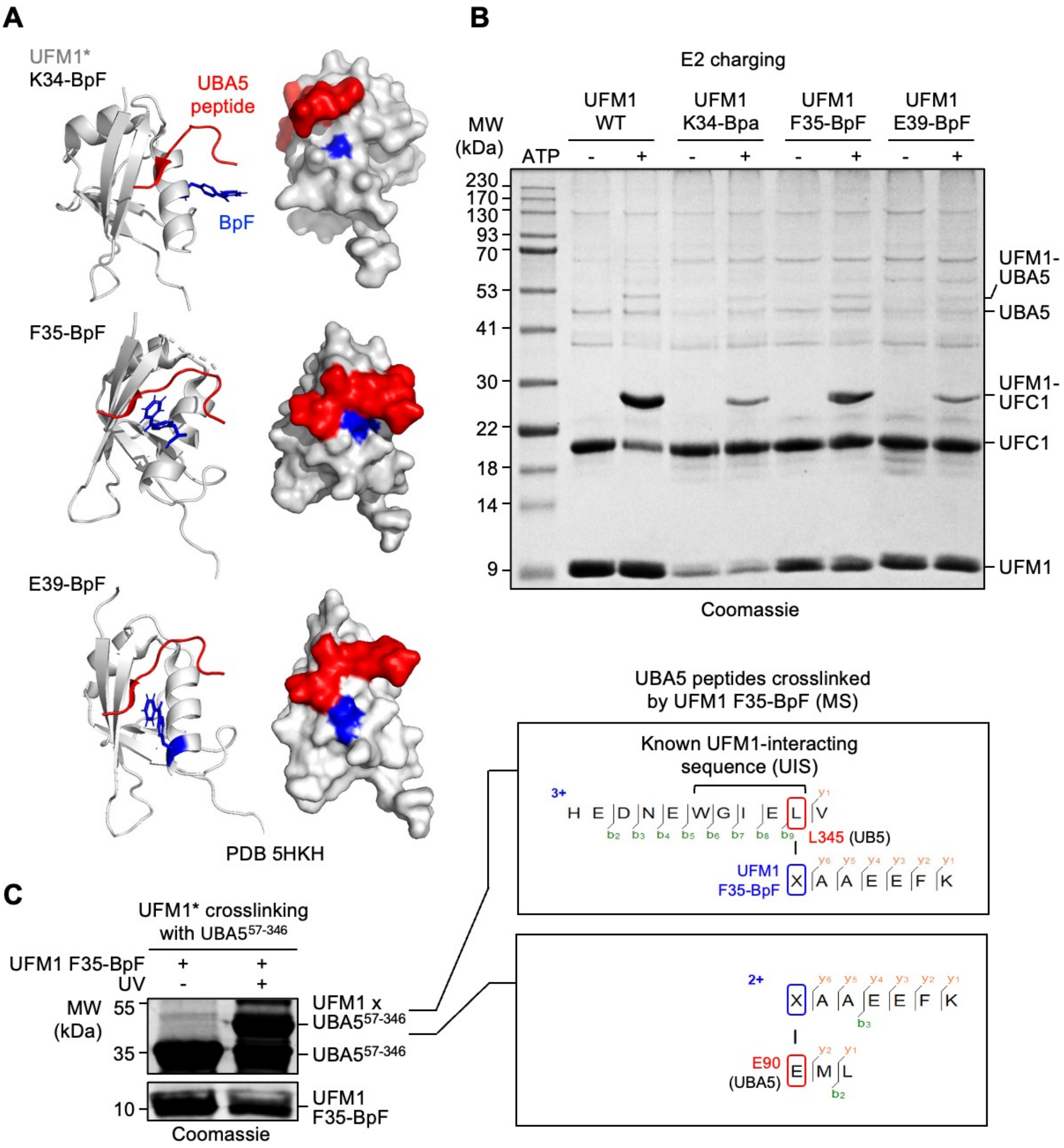


**Figure S4. Establishment of a photo-crosslinkable UFM1 probe for capturing weak and transient interactors with its α-β groove.** (**A**) Ribbon and surface representation of BpF incorporation into UFM1 (PBD 5HKH) at K34, F35, and E39 relative to binding of the UFM1-interacting sequence (UIS, amino acids 338-346) from UBA5. (**B**) E2 charging assays using UFM1, wildtype (WT) or mutants incorporating BpF at K34, F35, or E39, indicating the functional integrity of UFM1 F35-BpF (same gel as in Figure 2C). (**C**) Left: photo-crosslinking reaction (365 nm, 2 h) of UFM1 F35-BpF with recombinant UBA5$^{57-346}$. Right: two crosslinked peptides obtained from the reaction displayed on the left by mass spectrometry (MS), corresponding to the UIS, and the adenylation domain, of UBA5.

Abbreviations: MS: mass spectrometry; MW: molecular weight, UIS: UFM1-interacting sequence; WT: wildtype.

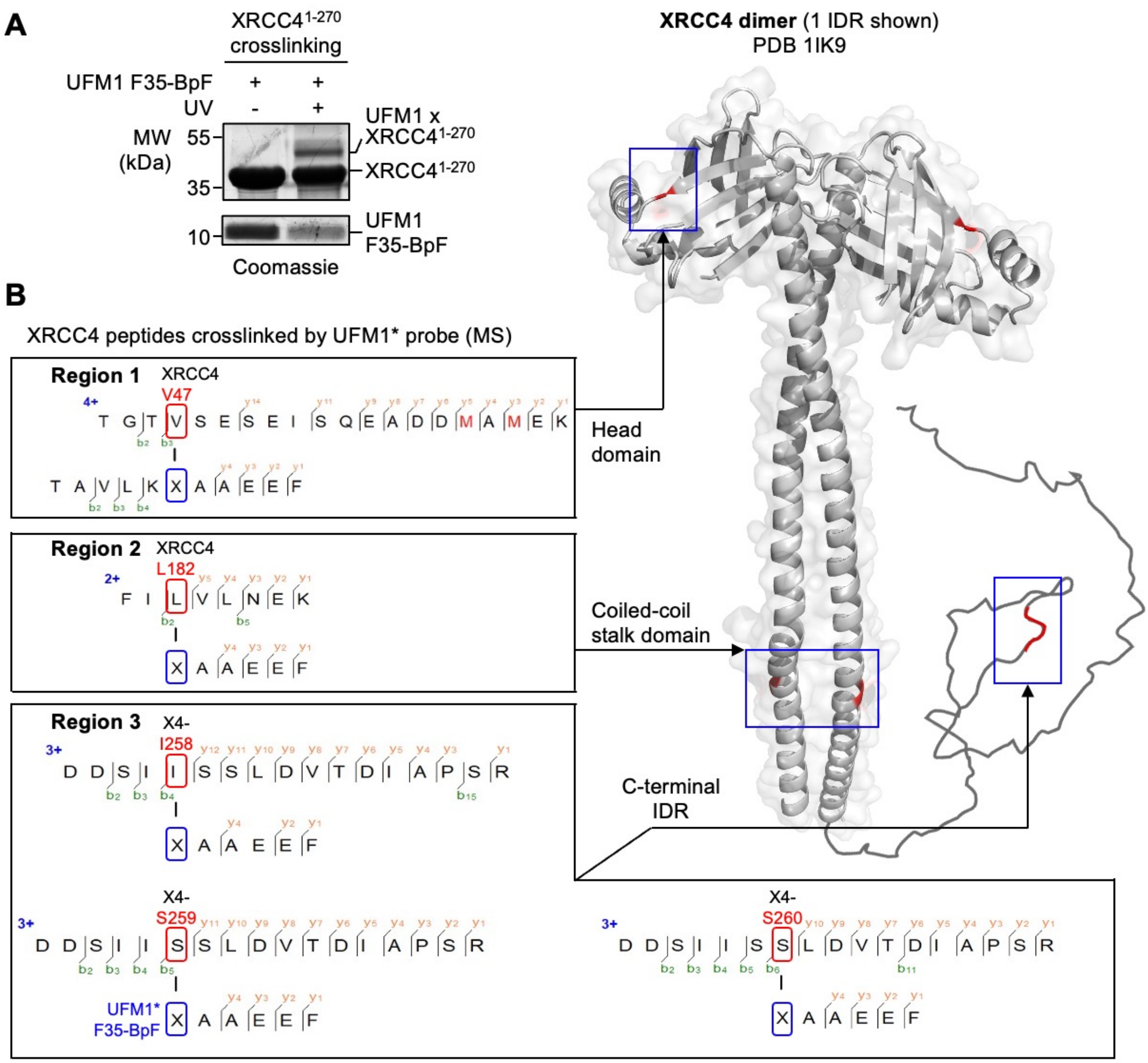


**Figure S5. Identification of UFM1 crosslinking regions on XRCC4.** (**A**) Photo-crosslinking reaction (365 nm, 2 h) of UFM1 F35-BpF with recombinant XRCC4$^{1\text{-}270}$. (**B**) Five crosslinked peptides obtained from the reaction in (A) and identified by mass spectrometry (MS), corresponding to three distinct regions on XRCC4 in the head domain (Region 1), the coiled-coil stalk (Region 2), and the C-terminal intrinsically disordered region (IDR, Region 3), as depicted on the right (PDB 1IK9). Crosslinked residues, valine 47 (V47), leucine 182 (L182), isoleucine 258 (I258) and serines 259 and 260 (S259, S260), are highlighted in red, F35-BpF in blue.

Abbreviations: IDR: intrinsically disordered region; MS: mass spectrometry; MW: molecular weight; UFM1*: UFM1 F35-BpF; X4: XRCC4.

### Supplemental Figure S6

**A**

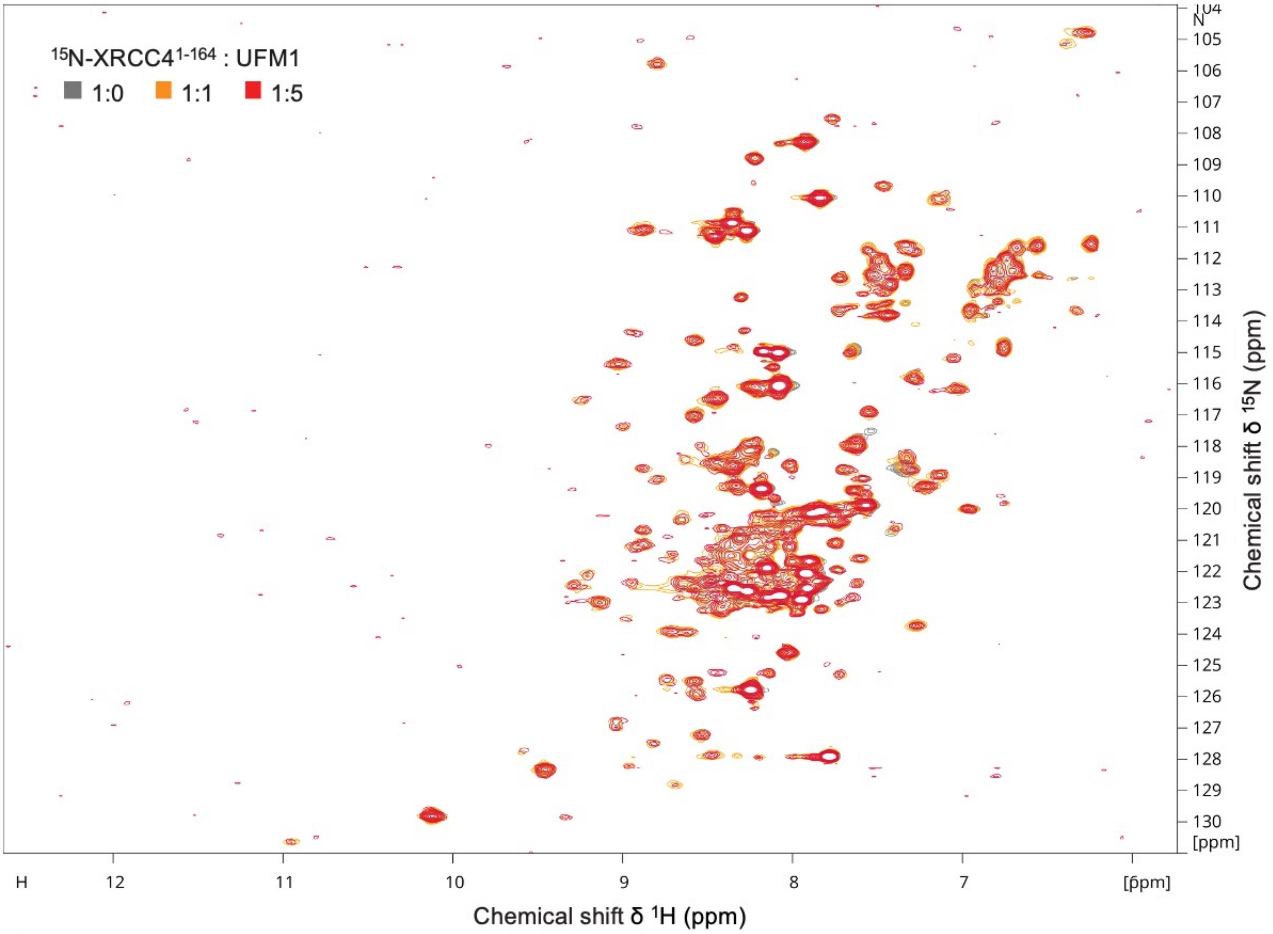


**B**

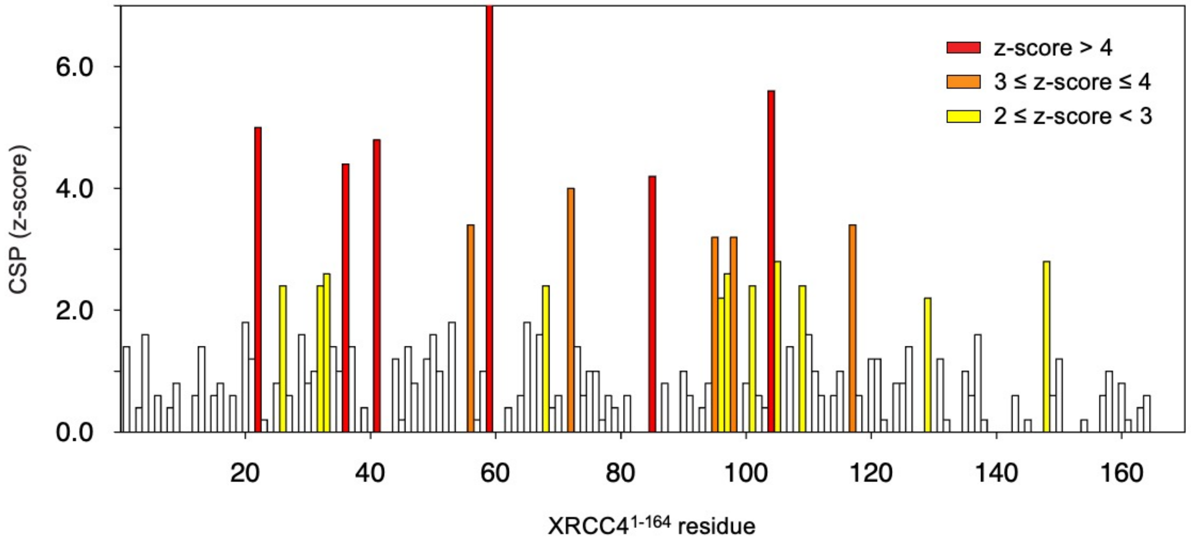


**Figure S6. NMR spectra and perturbations of XRCC4$^{1-164}$ binding to UFM1.** (**A**) Overlay of $^{1}$H-$^{15}$N BEST-TROSY spectra of XRCC4$^{1-164}$ alone (grey) and after adding 1 (orange) or 5 (red) equivalents of UFM1. (**B**) Z-scores of chemical shift perturbations (CSPs) extracted from the spectra shown in (A).

### Supplemental Figure S7

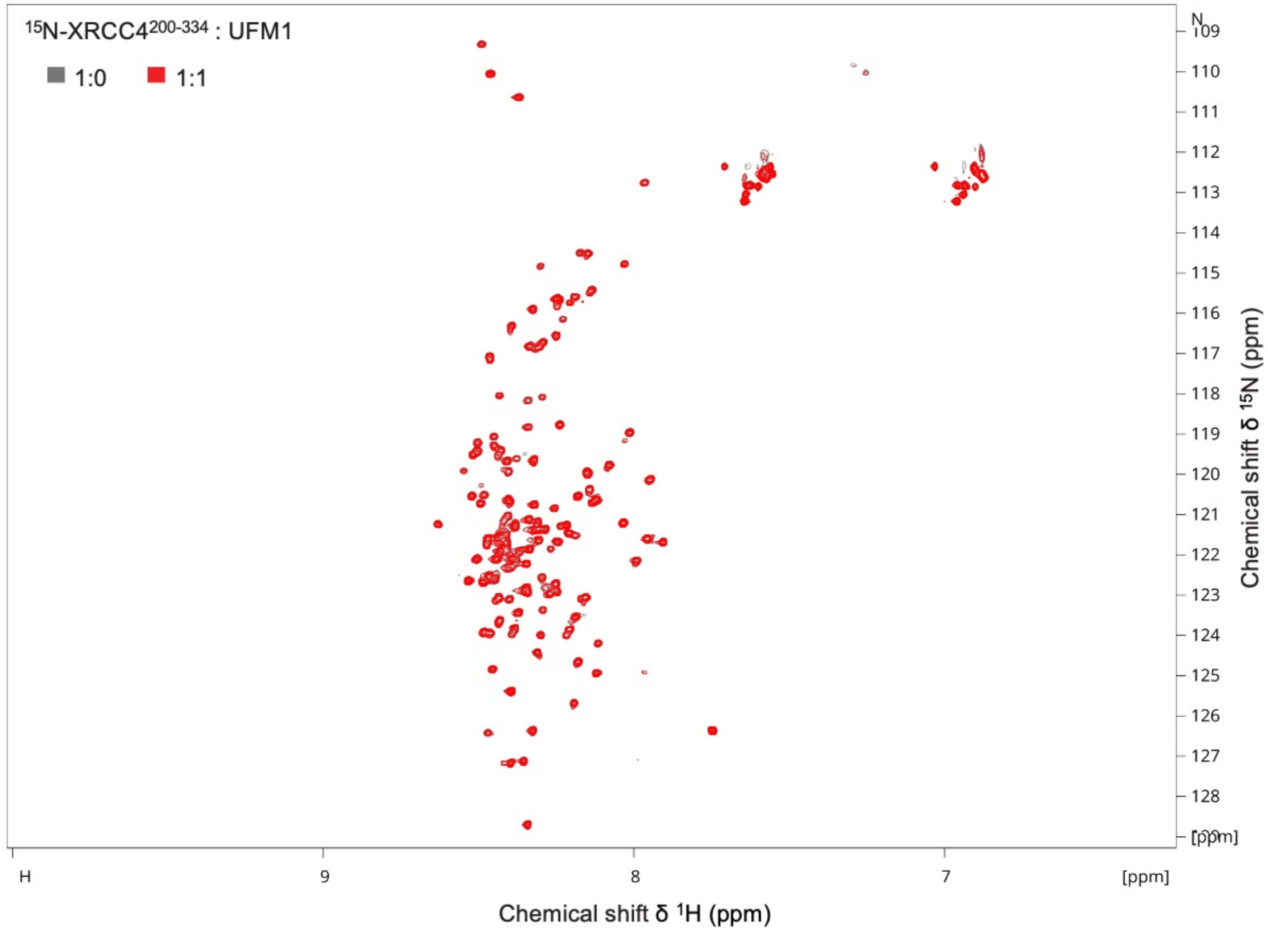


**Figure S7. NMR spectra of XRCC4$^{200\text{-}334}$ binding to UFM1.** Overlay of $^{1}H$-$^{15}N$ HSQC spectra of XRCC4$^{200\text{-}334}$ alone (grey) and in the presence of 1 equivalent of UFM1 (red).

#### Supplemental Figure S8

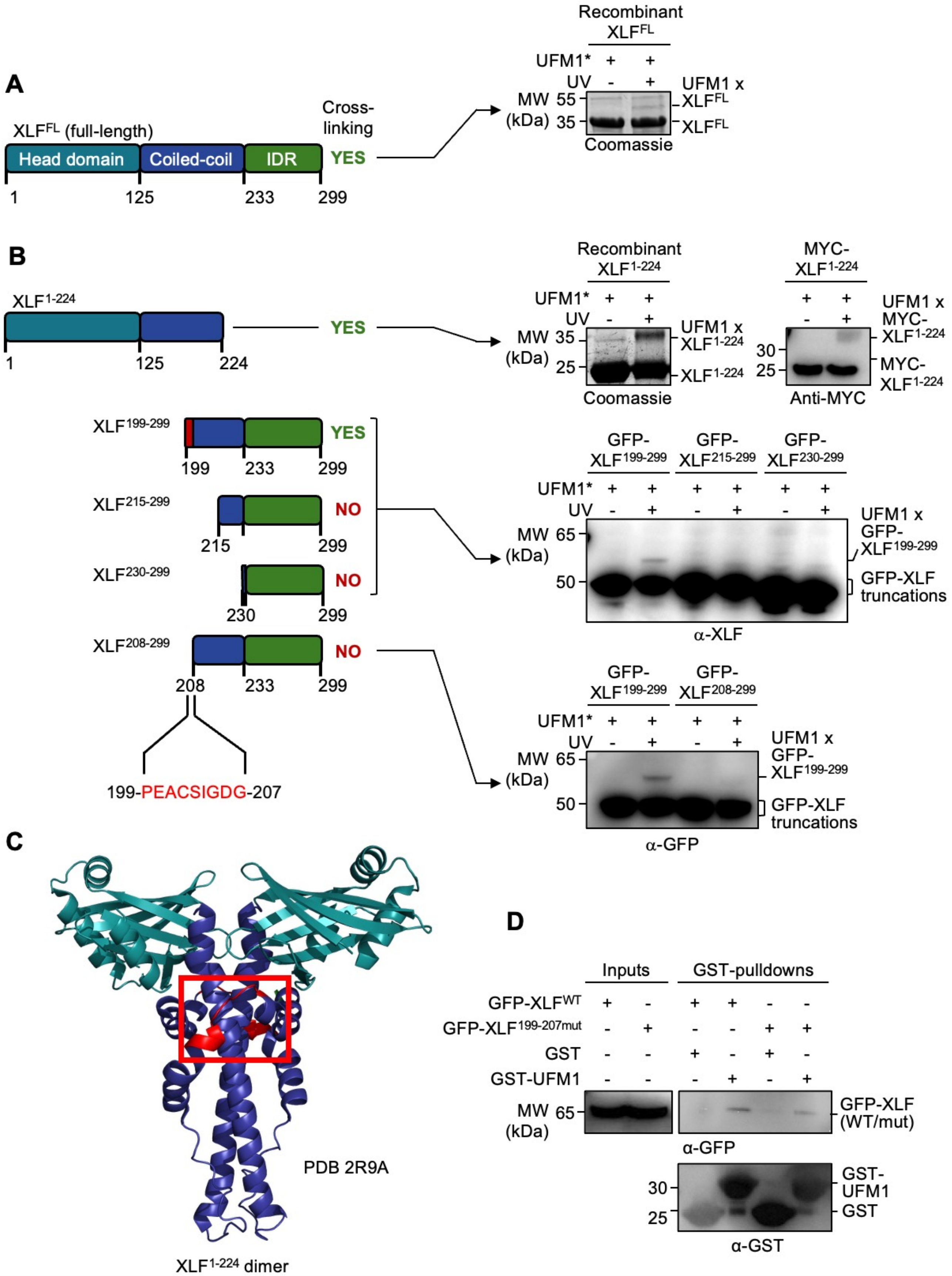

**Figure S8**. **Photo-crosslinking validation of XLF as a UFM1-interacting protein**. (**A**) Full-length purified XLF photo-crosslinks to the UFM1 F35-BpF probe (UFM1*). (**B**) Photo-crosslinking of UFM1* with $XLF^{1-224}$, either purified from *E. coli* or in HEK293T cell lysate following ectopic expression (top right two panels). Photo-crosslinking of UFM1* with HEK293T cell lysates transiently transfected with GFP-tagged XLF truncation mutants highlights a sequence (199-PEACSIGDG-207) that is required for binding to UFM1* (middle and bottom right panels). Schematics of XLF and the mutants used are shown on the left. (**C**) Structural representation of $XLF^{1-224}$ dimer (PDB 2R9A), highlighting the location of the UFM1-interacting region (red). (**D**) GST-UFM1 pulldowns with HEK293T cell lysates ectopically expressing GFP-tagged full-length XLF – wildtype ($XLF^{WT}$) or 199-207 mutated to GGGCGGGS ($XLF^{199-207mut}$) – highlighting reduced binding to UFM1 when this UFM1-interacting sequence is altered.

Abbreviations: α: anti; MW: molecular weight; UFM1*: UFM1 F35-BpF.

#### Supplemental Figure S9

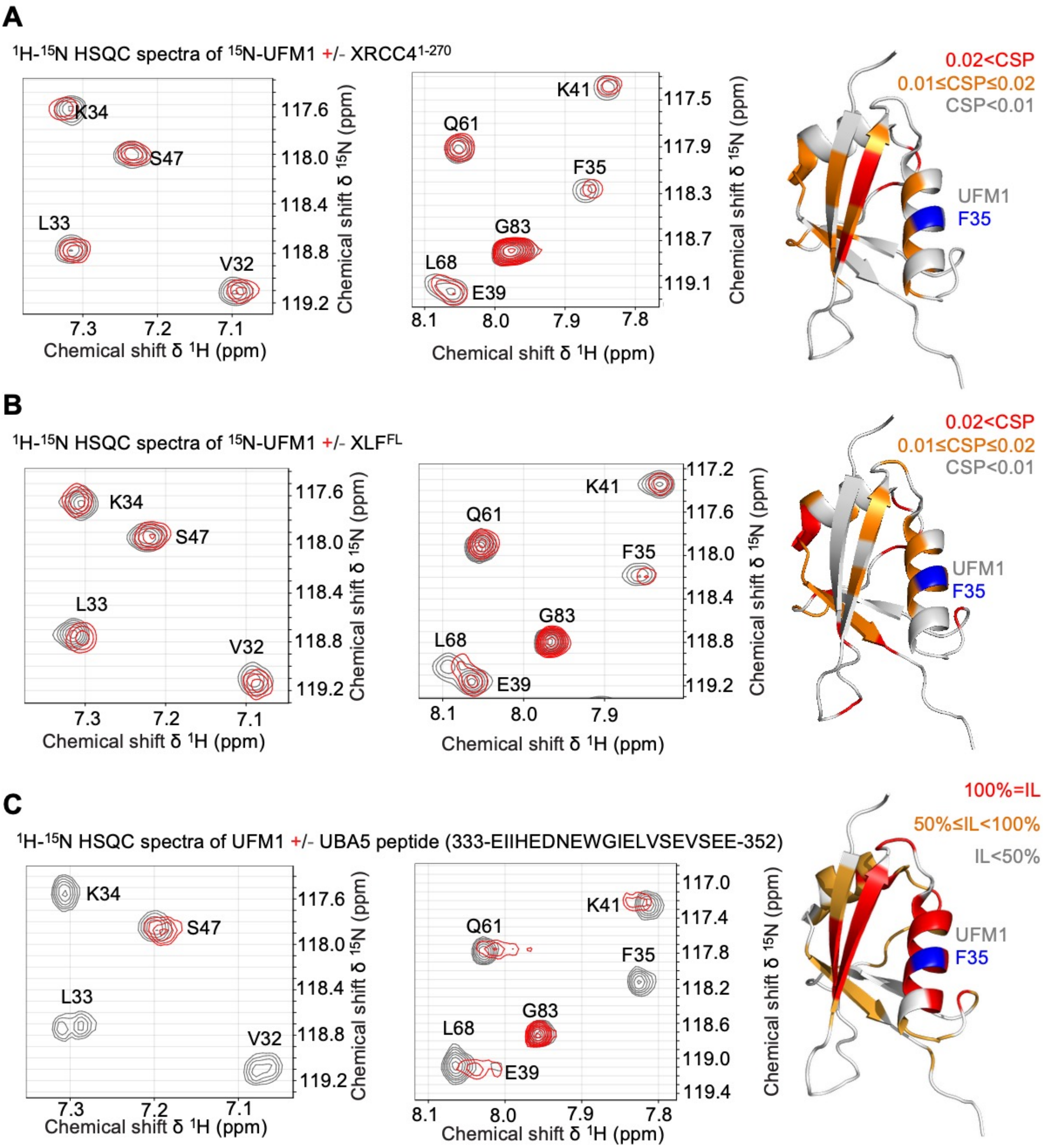


**Figure S9. XLF and XRCC4 target the α-β groove of UFM1.** Left: $^1$H-$^{15}$N HSQC spectra of $^{15}$N-UFM1 (grey), overlayed with the spectra after the addition of 1 equivalent (red) of XRCC4$^{1-270}$ (**A**), XLF$^{FL}$ (**B**), or a UBA5 peptide containing the UFM1 interacting sequence (UIS), as indicated (**C**), showing key residues affected. Right: Corresponding ribbon structures of UFM1 (PBD 5HKH), highlighting the most affected residues by chemical shift perturbation (CSP, XRCC4$^{1-270}$ and XLF$^{FL}$) or intensity loss (IL, UBA5 peptide). Colour code ranges from red (residues most affected) over orange (least affected) to grey (not affected), as indicated. F35-BpF is highlighted in blue.

Abbreviations: CSP: chemical shift perturbation; FL: full length; IL: intensity loss; UIS: UFM1-interacting sequence.

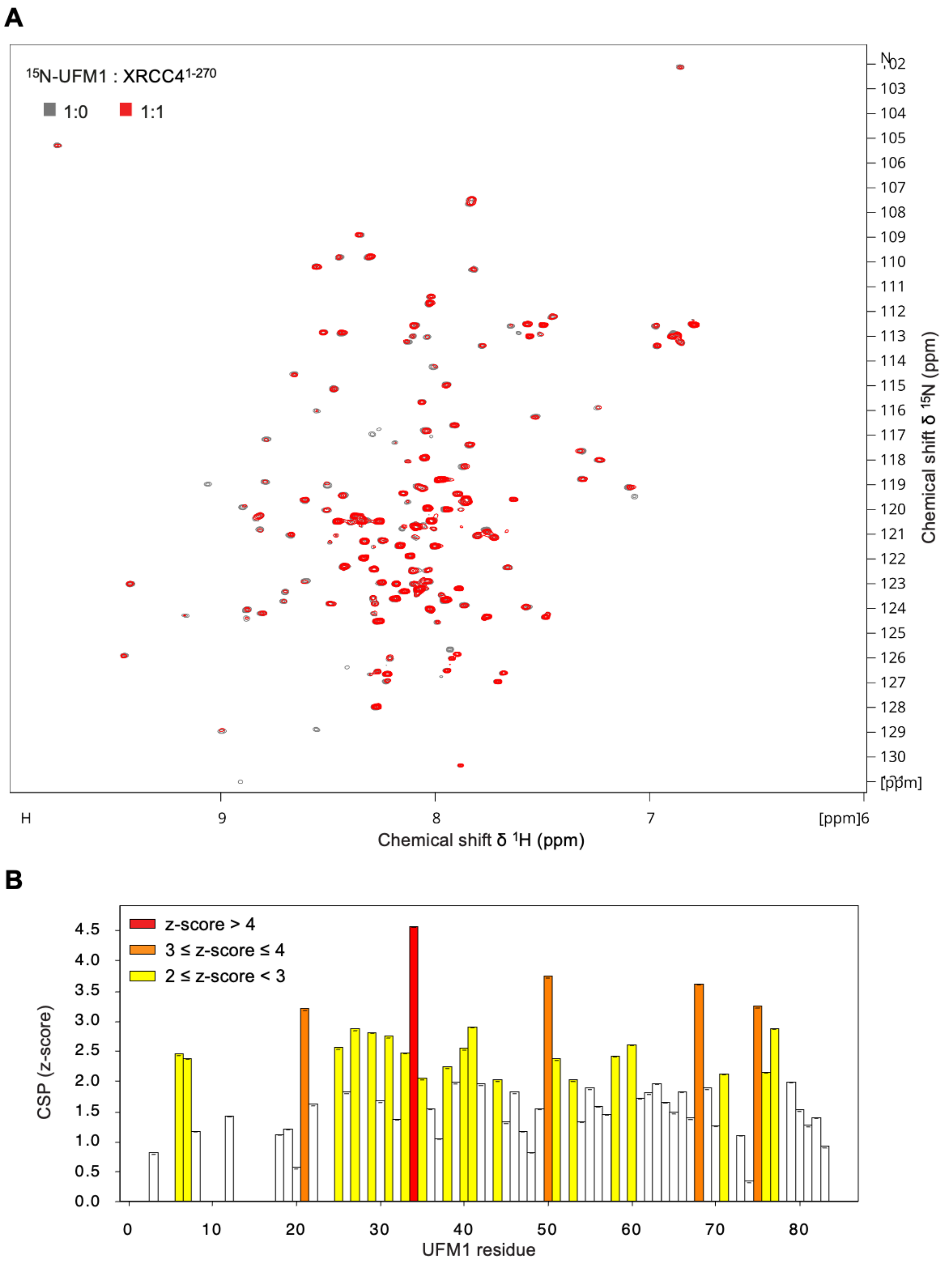


**Supplementary Figure S10. NMR spectra and perturbations of UFM1 binding to XRCC4$^{1-270}$.** (**A**) Overlay of $^{1}$H-$^{15}$N HSQC spectra of UFM1 alone (grey) and after adding 1 (red) equivalent of XRCC4$^{1-270}$. (**B**) Z-scores of chemical shift perturbations (CSPs) extracted from the spectra shown in (A).

### Supplemental Figure S11

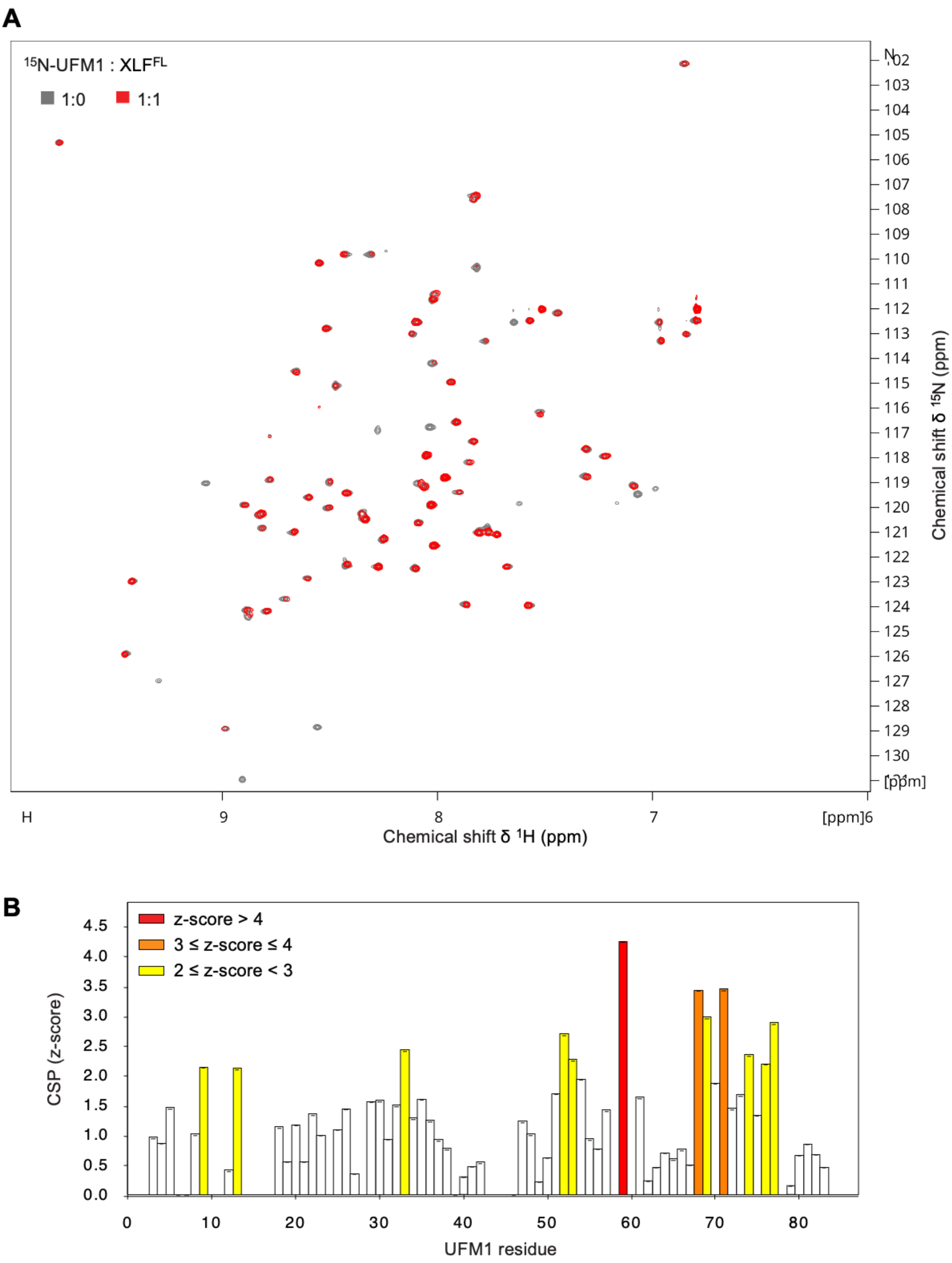


**Supplementary Figure S11. NMR spectra and perturbations of UFM1 binding to XLF$^{FL}$.** (**A**) Overlay of $^{1}$H-$^{15}$N HSQC spectra of UFM1 alone (grey) and after adding 1 (red) equivalent of full-length XLF (XLF$^{FL}$). (**B**) Z-scores of chemical shift perturbations (CSPs) extracted from the spectra shown in (A).

#### Supplemental Figure S12

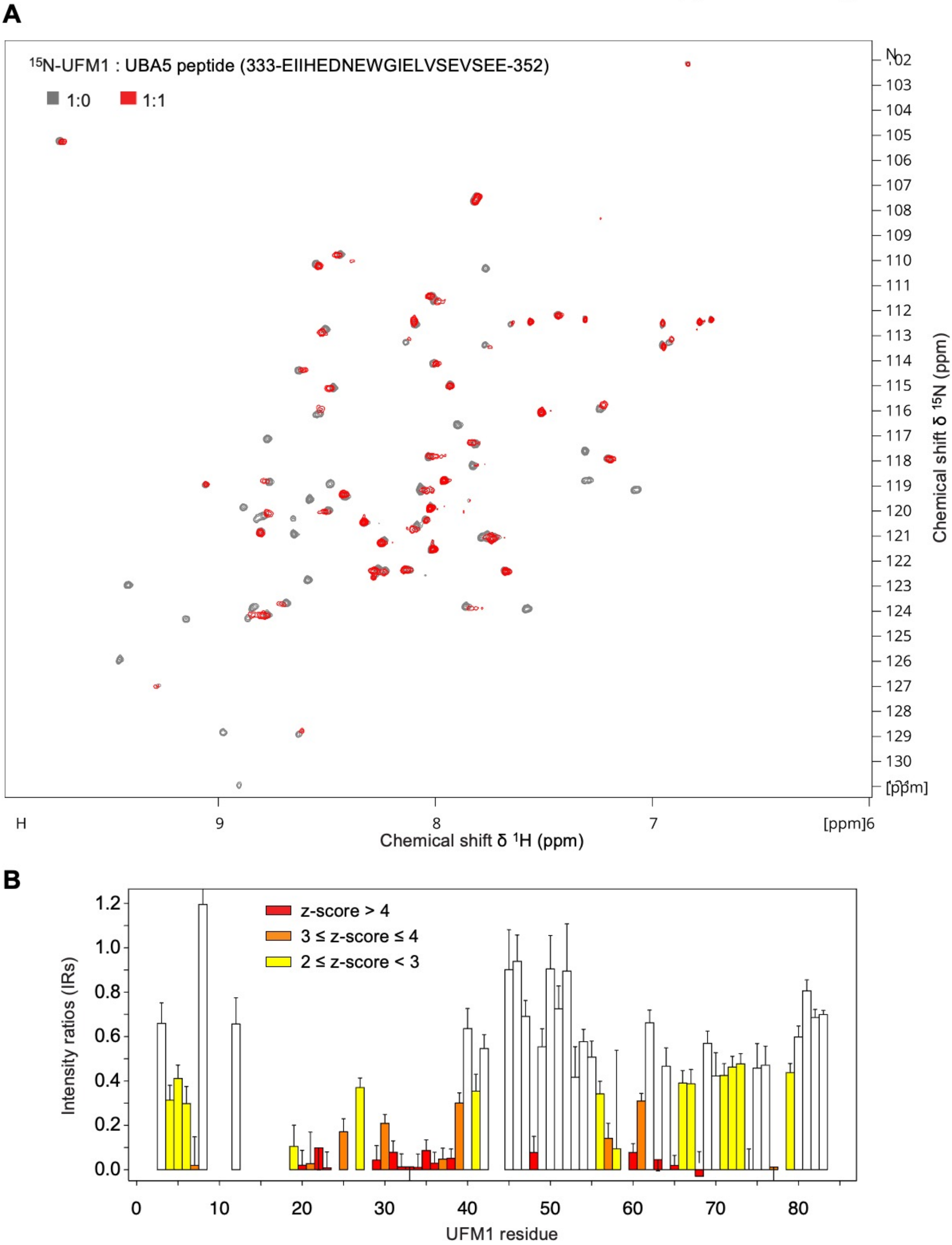


**Supplementary Figure S12. NMR spectra and intensity ratios of UFM1 binding to a UBA5 peptide.** (**A**) Overlay of $^{1}$H-$^{15}$N HSQC spectra of UFM1 alone (grey) and after adding 1 (red) equivalent of UBA5 peptide. (**B**) Intensity ratios extracted from the spectra shown in (A). Z-scores relate to chemical shift perturbations. Error bars represent 1 standard deviation propagated from the baseline deviation.

### Supplemental Figure S13

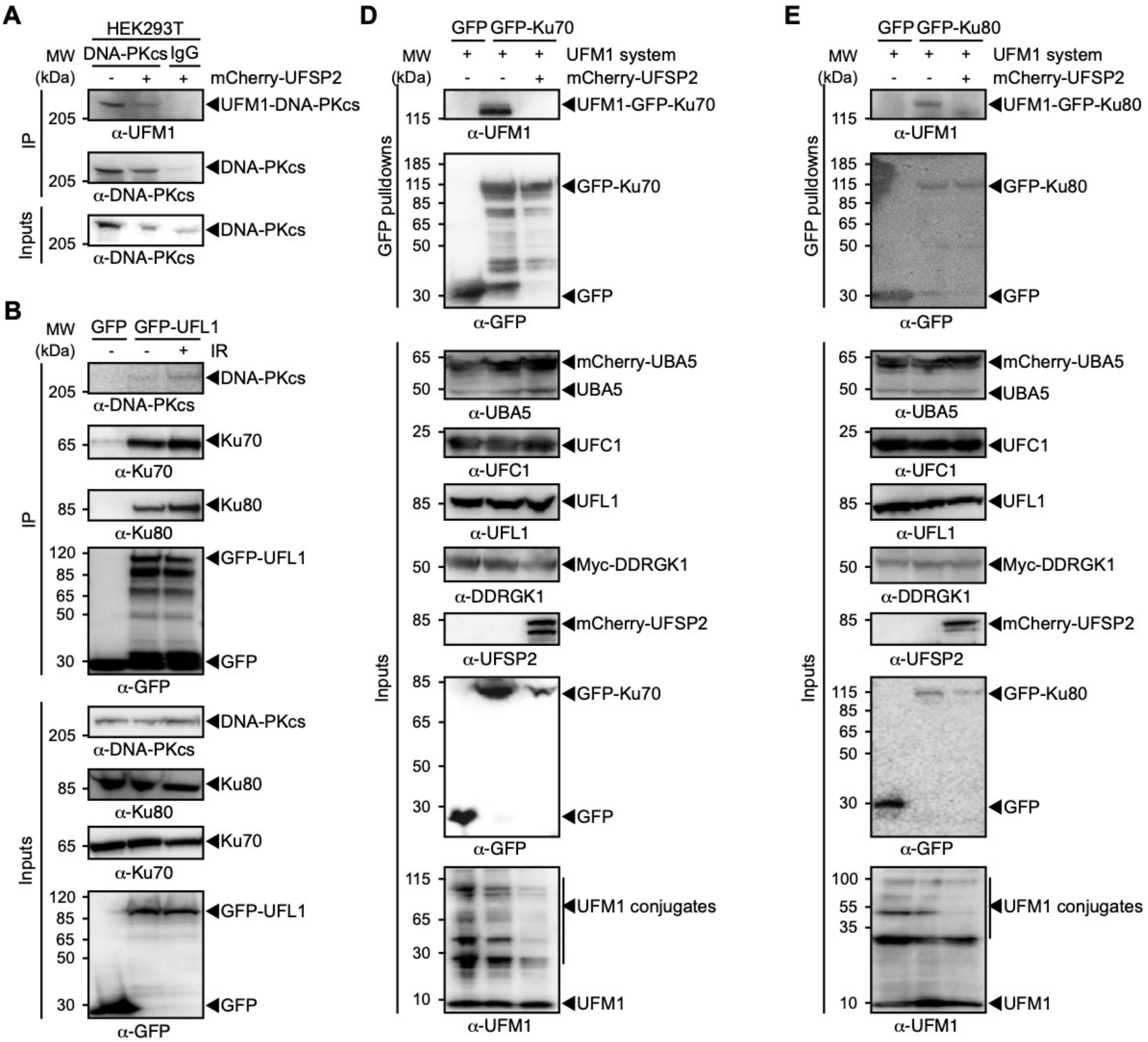


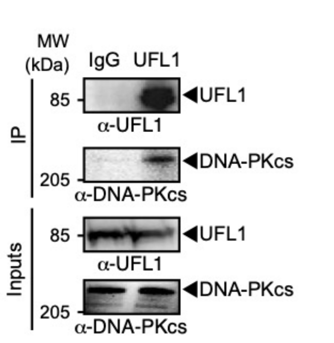


**Figure S13. UFMylation of DNA-PK complex components.** (**A**) Immunoprecipitation (IP) of DNA-PKcs from whole cell extracts, showing a slowly migrating band detectable by a-UFM1 consistent with UFMylation of DNA-PKcs, reduced in the presence of ectopically expressed deUFMylase UFSP2. Whole cell extracts were obtained from HEK293T cells. (**B**) Interaction between UFL1 and the DNA-PK holoenzyme, as shown by GFP-UFL1 pulldown co-precipitating endogenous DNA-PKcs, Ku70 and Ku80. Whole cell extracts were obtained from HEK293T cells, treated, or not, with ionising radiation (IR, ~30 min post 10 Gy). (**C**) Immunoprecipitation (IP) of endogenous UFL1 showing co-precipitation with endogenous DNA-PKcs in whole cell extracts obtained from HEK293T cells. (**D**) GFP-pulldowns of Ku70 from whole cell extracts, showing an a-UFM1-detectable band migrating at a molecular weight consistent with Ku70 UFMylation, which was reduced in the presence of ectopically expressed deUFMylase UFSP2. Whole cell extracts were obtained from HEK293T cells, ectopically expressing GFP-Ku70 and UFM1 system components (UBA5, UFC1, UFL1, DDRGK1) together, or not, with UFSP2. (**E**) As (D) but for GFP-Ku80.

Abbreviations: α: anti; IP: immunoprecipitation; IR: ionising radiation, MW: molecular weight; WT: wildtype.
